# Long-term (1990-2019) monitoring of tropical moist forests dynamics

**DOI:** 10.1101/2020.09.17.295774

**Authors:** C. Vancutsem, F. Achard, J.-F. Pekel, G. Vieilledent, S. Carboni, D. Simonetti, J. Gallego, L. Aragao, R. Nasi

## Abstract

Accurate characterization of the tropical moist forests changes is needed to support conservation policies and to better quantify their contribution to global carbon fluxes. We document - at pantropical scale - the extent of these forests and their changes (degradation, deforestation and recovery) over the last three decades. We estimate that 17% of the tropical moist forests have disappeared since 1990 with a remaining area of 1060 million ha in 2019, from which 8.5% are degraded. Our study underlines the importance of the degradation process in such ecosystems, in particular as precursor of deforestation and in the recent increase of the tropical moist forest disturbances. Without reduction of the present disturbance rates, undisturbed forests will disappear entirely in large tropical humid regions by 2050. Our study suggests reinforcing actions to prevent the first disturbance scar that leads to forest clearance in 45% of the cases.

## INTRODUCTION

Tropical moist forests (TMF) have a huge environmental value. They play an important role in biodiversity conservation, terrestrial carbon cycle, hydrological regimes, indigenous population subsistence and human health (**1–5**). They are increasingly recognized as an essential element of any strategy to mitigate climate change (**6, 7**). Deforestation, and degradation compromise the functioning of tropical forests as an ecosystem, lead to biodiversity loss (**1, 4, 5, 8, 9**) and reduced carbon storage capacity (**10–17**). Deforestation and fragmentation are increasing the risk of virus disease outbreaks (**18–20**).

For humanity wellbeing, sustainable economic growth and conservation of the remaining TMF constitute one of the largest challenges and shared responsibility. A consistent, accurate and geographically explicit characterization of the long-term disturbances at the pantropical scale is a prerequisite for elaborating a coherent territorial planning towards Sustainable Development Goals (SDGs) and the Nationally determined contributions (NDCs) of the Paris Agreement (2015). Advances in remote-sensing, cloud computing facilities, and free access to the Landsat satellite archive (**21–23**), enable systematic monitoring and consistent dynamic characterization of the entire TMF across a long period. Global maps have been derived to quantify tree cover loss since 2000 (**24–25**) and to identify remaining intact forest landscapes (**17**). However, detailed spatial information on the long-term dynamics of tropical moist forests and particularly on forest degradation and post-disturbances development stages is still missing to accurately estimate the carbon loss associated with forest disturbances (**2, 13, 15**) and assess their impact on biodiversity (**5, 8**).

## RESULTS AND DISCUSSION

Here we provide new information through a wall-to-wall mapping of tropical moist forest cover dynamics over a long-term period (January 1990 to December 2019) at 0.09 ha resolution (freely available from https://forobs.jrc.ec.europa.eu/TMF/) (see Materials and Methods). This validated dataset depicts the TMF extent and the related disturbances (deforestation and degradation), and post-disturbances recovery on an annual basis over the last three decades (see Supplementary Text on the annual change dataset, and fig. S1). A major innovation consists of characterizing the sequential dynamics of changes by providing transition stages from the initial observation period to the end of the year 2019, i.e. undisturbed forest, degraded forest, forest regrowth, deforested land, conversion to plantations, conversion to water, afforestation, and changes within the mangroves (**Figs. 1** **and** **2**, see Supplementary Text on the transition map and figs. S2 to S7), as well as the timing (dates and duration), recurrence and intensity of each disturbance.

**Fig. 1.**
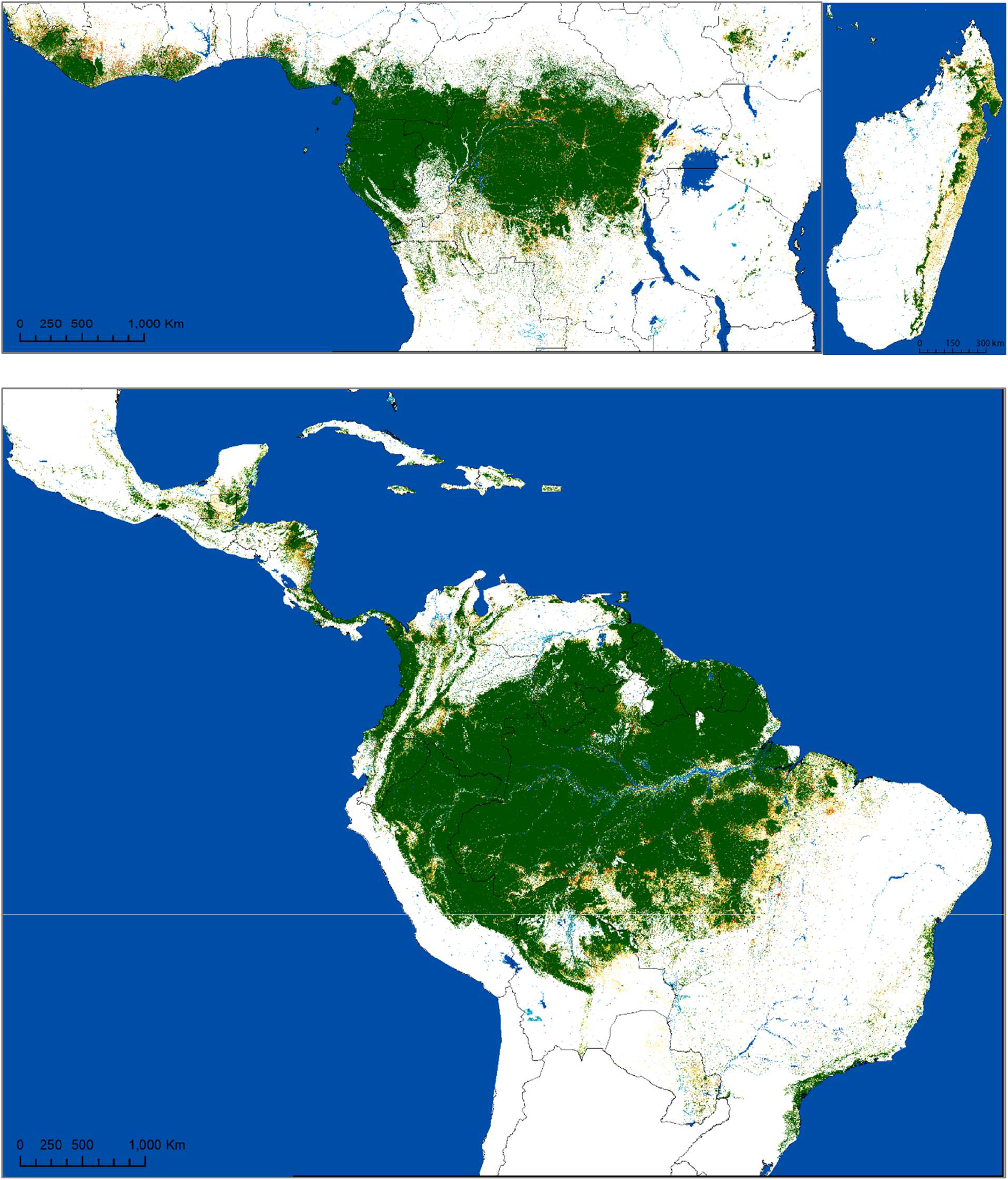
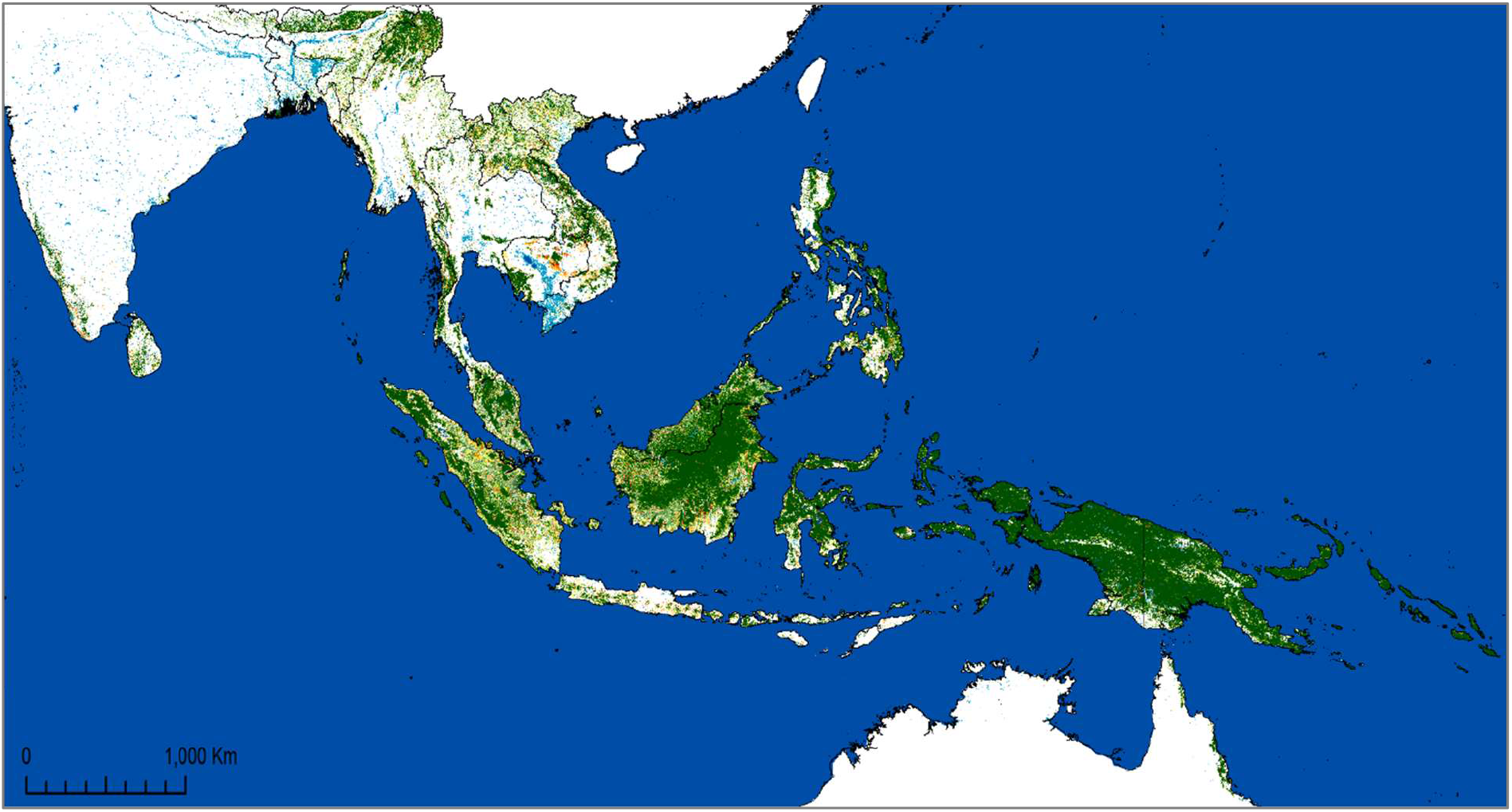
Map of tropical moist forests remaining in January 2020 and disturbances observed during the period 1990-2019. See legend in Figs. 2.

**Fig. 2.**
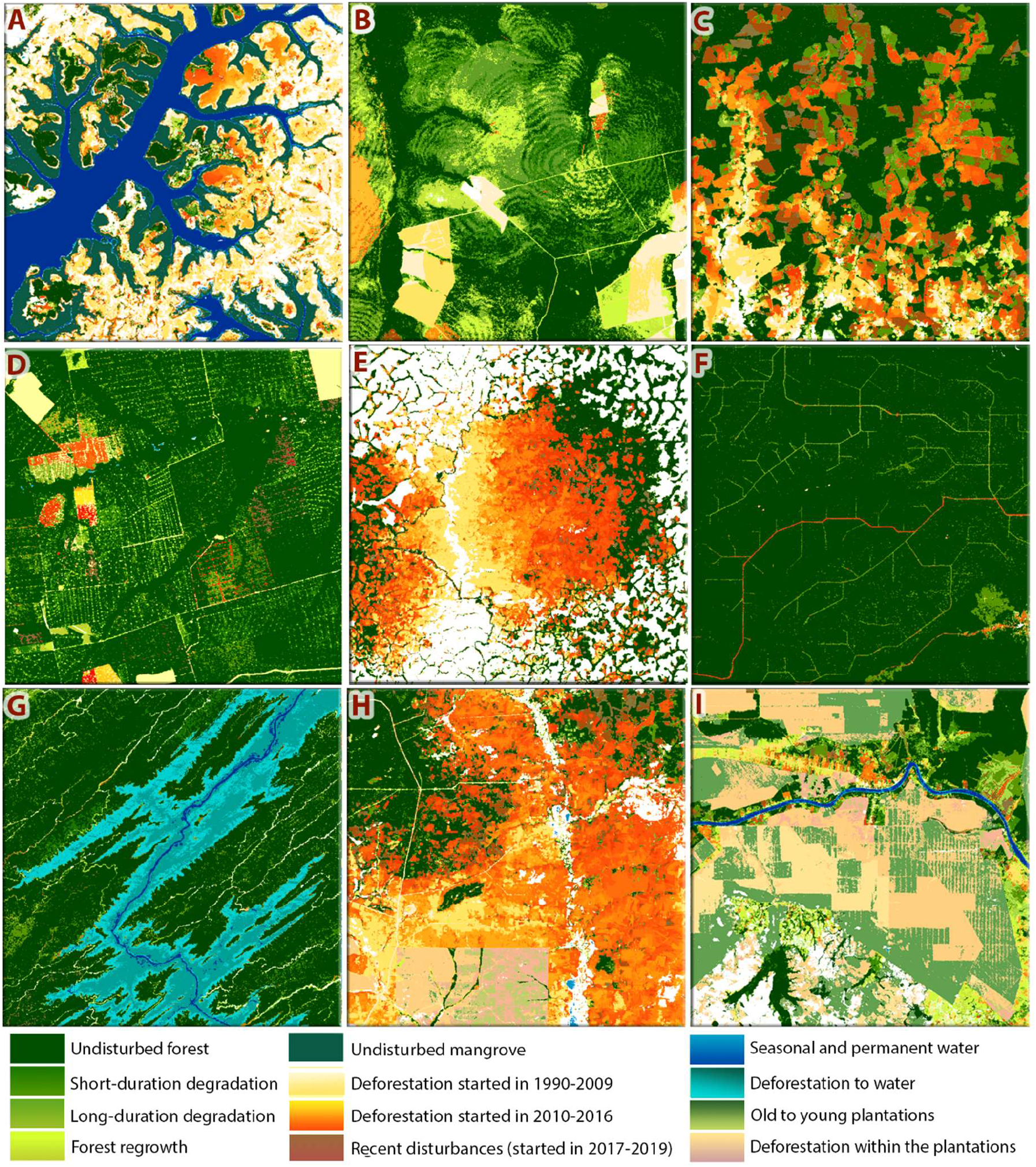
Examples of patterns of forest cover disturbances (deforestation and degradation) during the period 1990-2019: (A) Remaining Mangroves and the related changes in Guinea-Bissau (14.9°W, 11.1°N), (B) Fires in Mato-Grosso province of Brazil (53.8°W, 13°S), (C) Recent deforestation in Colombia (74.4°W, 0.7°N), (D) Logging in Mato-Grosso (54.5°W, 12°S), (E) Deforestation and degradation caused by the railway in Cameroon (13.4°E, 5.8°N) (F) Recent selective logging in Ouesso region of Republic of Congo (15.7°E, 1.4°N), (G) Deforestation for the creation of a dam in Malaysia (113.8°E, 2.4°S), (H) Massive deforestation in Cambodia (105.6°E, 12.7°N), and (I) Commodities in the Riau province of Indonesia (102°E, 0.4°N). The size of each box is 20 km × 20 km.

For the first time at the pantropical scale the occurrence and extent of the forest cover degradation is documented on an annual basis in addition to the deforestation. This has been achieved thanks to the analysis of each individual valid observation of the Landsat archive (see Data and Mapping method Sections) allowing to capture short-duration disturbances such as selective logging (**Fig. 2F**, fig. S3), fires (**Fig. 2B**), and severe weather events (hurricanes, dryness) (fig. S7).

The accuracy of the disturbance mapping is 91.4%. Uncertainties in the area estimates were quantified based on a sample-based reference in accordance with the latest statistical good practices (**26**) and indicates an underestimation of the forest disturbance areas by 11.8% (representing 38.4 million ha, with 15 million ha confidence interval at 95%) (see Section and Supplementary Text on the validation, figs. S8 to S10 and tables S1 to S4).

### Main results on degradation

The analysis of the yearly dynamics of TMF disturbances over the last 30 years underlines the importance of the degradation process in tropical moist forest ecosystems with the following key outcomes (**Tables 1**, **2** **and** **3**, the Trend analysis section in Materials and Methods, fig. S11):

i. During the last three decades, 195.1 million ha of TMF have disappeared and 106.5 million ha are in a degraded status (**Table 3**). This represents 8.4% of the 1059.6 million ha of forest area remaining in January 2020. Degraded forests represent 33% of the observed disturbances with high variability between regions and countries, ranging from 96% in Venezuela, 74% in Gabon, and 69% in Papua New Guinea to 21% in Brazil and Madagascar, and 13% in Cambodia (Table S6). 40.7% of the degraded forests are in Asia-Oceania (compared to 36.9% in Latin America and 22.3% in Africa) (**Table 3**).
ii. 84.5% of the degraded forests (i.e. 90 million ha) are resulting from short-term disturbances (observed over less than 1-year duration, mostly due to selective logging, natural events and light-impact fires), from which 30 million ha have been degraded repeatedly 2 or 3 times over the last 30 years (observed each time along a short-term period). The remaining 15.5% (16.5 million ha) are mainly resulting from intense fires, with a disturbance duration between 1 to 2.5 years.
iii. 45.4% of the degradation (88.6 million ha) is a precursor of deforestation events occurring on average after 7.5 years (without significant variability between continents). This is particularly true for South-East Africa and South-East Asia that show respectively 60.4% (with 65% for Madagascar) and 53% (with 59% for Cambodia) of degraded forests becoming deforested in a second step (**Table 2**). These proportions are underestimated because 45.4% of recent degradation (e.g. in the last 7 years) will most likely be deforested in future years.
iv. A further 30.3% of the undisturbed forest areas (291.8 million ha) are potentially disturbance-edge-affected forests, i.e. located within 120 meters from a disturbance (see Materials and Methods). This proportion indicates a higher forest fragmentation proportion in Asia (45.2%) compared to other continents (25.6% and 28.9% respectively in the Americas and Africa).
v. 82.8% of the TMF mapped as degraded in December 2019 corresponds to short-term disturbances that have never been identified so far at the pan-tropical scale. Over the period covered by the Global Forest Change (GFC) product (**24**), i.e. 2001-2019, 21.2 million ha have been captured as a tree cover loss compared to 86 million ha detected as degraded forests by our study during the same period (see Section on the comparison with the GFC dataset, **Fig. 4** and table S5).
vi. We show that the annual rate of degradation is highly related to climatic conditions (**Figs. 3** **and** **4**, fig. S11). Whereas the trends in deforestation rates seem to be related to changes in national territorial policies, degradation rates usually show peaks during drought periods and do not seem to be impacted by forest conservation policies. The drought conditions that occurred during strong and very strong El Niño southern oscillation (ENSO) events of 1997-1998 and 2015-2016 were optimal for forest fires (**27–29**) and resulted in a strong increase of forest degradation (**28**). The impact of these fires in 2015-2016 is particularly strong and visible in all regions except in South-East Africa.

**Table 1.** Areas (in million ha) of (a) undisturbed tropical moist forests (TMF) and (b) Undisturbed and degraded TMF for years 1990, 1995, 2000, 2005, 2010, 2015 and 2020 (on first January) by sub-region and continent, and relative decline (in %) over intervals of 30 years (1990-2020), and 10 years (1990-2000, 2000-20100, 2010-2020). The values appearing in grey color indicate values derived from an average percentage of invalid pixel observations over the baseline TMF domain higher than 40%.

| Sub-region | Area of Undisturbed TMF on 1st January (Mha) |  |  |  |  |  |  | Decline (% of the forest) |  |  |  |
| --- | --- | --- | --- | --- | --- | --- | --- | --- | --- | --- | --- |
|  | 1990 | 1995 | 2000 | 2005 | 2010 | 2015 | 2020 | [1990-2020] | [1990-2000] | [2000-2010] | [2010-2020] |
| West-Africa | 34.6 | 34.1 | 32.8 | 27.4 | 23.9 | 20.6 | 15.6 | 55.0 | 5.0 | 27.1 | 35.0 |
| Central-Africa | 223.1 | 221.5 | 216.1 | 207.2 | 201.1 | 193.7 | 184.7 | 17.2 | 3.1 | 6.9 | 8.2 |
| South-East Africa | 15.7 | 15.0 | 12.5 | 10.1 | 8.9 | 7.6 | 6.4 | 59.2 | 20.7 | 28.6 | 27.9 |
| Central-America | 34.5 | 32.3 | 27.4 | 24.1 | 21.7 | 19.6 | 16.2 | 53.0 | 20.8 | 20.6 | 25.3 |
| South-America | 670.6 | 655.4 | 628.8 | 600.9 | 583.2 | 568.9 | 548.2 | 18.2 | 6.2 | 7.3 | 6.0 |
| Continental SE Asia | 73.3 | 67.2 | 57.7 | 50.2 | 44.4 | 39.9 | 34.2 | 53.3 | 21.2 | 23.2 | 22.9 |
| Insular SE Asia | 237.9 | 229.5 | 207.8 | 192.9 | 180.8 | 170.5 | 159.1 | 33.1 | 12.6 | 13.0 | 12.0 |
| **Continent** | **1990** | **1995** | **2000** | **2005** | **2010** | **2015** | **2020** | **[1990-2020]** | **[1990-2000]** | **[2000-2010]** | **[2010-2020]** |
| Africa | 273.4 | 270.6 | 261.4 | 244.7 | 234.0 | 221.9 | 206.7 | 24.4 | 4.4 | 10.5 | 11.7 |
| Latin-America | 705.1 | 687.7 | 656.1 | 625.0 | 604.9 | 588.5 | 564.4 | 19.9 | 6.9 | 7.8 | 6.7 |
| Asia-Oceania | 311.1 | 296.7 | 265.6 | 243.2 | 225.2 | 210.4 | 193.3 | 37.9 | 14.6 | 15.2 | 14.2 |
| **Total** | **1267.1** | **1232.4** | **1160.6** | **1090.4** | **1041.5** | **998.2** | **964.4** | **23.9** | **8.4** | **10.3** | **7.4** |

**(b)**
| Sub-region | Area of TMF (undisturbed and degraded) at the end of the year (Mha) |  |  |  |  |  |  | Decline (% of the forest) |  |  |  |
| --- | --- | --- | --- | --- | --- | --- | --- | --- | --- | --- | --- |
|  | 1990 | 1995 | 2000 | 2005 | 2010 | 2015 | 2020 | [1990-2020] | [1990-2000] | [2000-2010] | [2010-2020] |
| West-Africa | 34.6 | 34.3 | 33.3 | 29.8 | 27.4 | 25.1 | 22.1 | 36.0 | 3.6 | 17.8 | 19.3 |
| Central-Africa | 223.1 | 222.3 | 218.6 | 212.7 | 208.9 | 204.4 | 199.9 | 10.4 | 2.0 | 4.4 | 4.3 |
| South-East Africa | 15.7 | 15.2 | 12.9 | 11.0 | 10.1 | 9.2 | 8.5 | 45.9 | 17.8 | 21.8 | 15.9 |
| Central-America | 34.5 | 32.8 | 29.0 | 26.8 | 25.3 | 24.0 | 22.1 | 35.9 | 16.0 | 12.8 | 12.5 |
| South-America | 670.6 | 657.7 | 635.5 | 613.7 | 601.0 | 592.4 | 581.6 | 13.3 | 5.2 | 5.4 | 3.2 |
| Continental SE Asia | 73.3 | 69.1 | 61.3 | 55.7 | 52.0 | 49.2 | 46.4 | 36.6 | 16.3 | 15.2 | 10.6 |
| Insular SE Asia | 237.9 | 232.6 | 218.4 | 208.9 | 201.2 | 194.9 | 190.2 | 20.0 | 8.2 | 7.9 | 5.5 |
| **Continent** | **1990** | **1995** | **2000** | **2005** | **2010** | **2015** | **2020** | **[1990-2020]** | **[1990-2000]** | **[2000-2010]** | **[2010-2020]** |
| Africa | 273.4 | 271.7 | 264.8 | 253.5 | 246.4 | 238.7 | 227.7 | 16.7 | 3.1 | 7.0 | 7.6 |
| Latin-America | 705.1 | 690.5 | 664.5 | 640.5 | 626.3 | 616.4 | 599.2 | 15.0 | 5.8 | 5.7 | 4.3 |
| Asia-Oceania | 311.1 | 301.7 | 279.7 | 264.5 | 253.2 | 244.1 | 232.8 | 25.2 | 10.1 | 9.5 | 8.1 |
| **Total** | **1267.1** | **1241.4** | **1186.5** | **1136.0** | **1103.4** | **1076.6** | **1059.6** | **16.4** | **6.4** | **7.0** | **4.0** |

**Table 2.** Average annual losses of undisturbed tropical moist forest areas (in million ha) between 1990 and 2020 over intervals of 5 year, 30 year (1990-2020), 20 year (2000-2020), and 10 years (1990-2000,2000-2010, 2010-2020) by sub-region and continent: (a) annual losses due to deforestation and degradation, (b) annual losses due to deforestation, (c) annual losses due to degradation, (d) annual losses due to direct deforestation, (e) annual degradation before deforestation, (f) annual losses due to deforestation followed by regrowth and (g) average percentage of invalid observations over the baseline TMF domain. The values appearing in grey color indicate values derived from an average percentage of invalid observations higher than 40%.

|  | Annual loss of Undisturbed TMF areas (Mha) |  |  |  |  |  |  |  |  |  |
| --- | --- | --- | --- | --- | --- | --- | --- | --- | --- | --- |
| Sub-region | [1990-1995] | [1995-2000] | [2000-2005] | [2005-2010] | [2010-2015] | [2015-2020] | [1990-2020] | [1990-2000] | [2000-2010] | [2010-2020] |
| West-Africa | 0.10 | 0.24 | 1.08 | 0.70 | 0.67 | 1.01 | 0.6 | 0.2 | 0.9 | 0.8 |
| Central-Africa | 0.33 | 1.07 | 1.79 | 1.22 | 1.49 | 1.79 | 1.3 | 0.7 | 1.5 | 1.5 |
| South-East Africa | 0.13 | 0.52 | 0.47 | 0.24 | 0.26 | 0.24 | 0.3 | 0.3 | 0.4 | 0.3 |
| Central-America | 0.45 | 0.99 | 0.66 | 0.47 | 0.43 | 0.67 | 0.6 | 0.7 | 0.6 | 0.6 |
| South-America | 3.03 | 5.33 | 5.56 | 3.56 | 2.85 | 4.14 | 4.1 | 4.2 | 4.6 | 4.1 |
| Continental SE Asia | 1.21 | 1.90 | 1.50 | 1.17 | 0.89 | 1.14 | 1.3 | 1.6 | 1.3 | 1.2 |
| Insular SE Asia | 1.68 | 4.32 | 2.98 | 2.43 | 2.07 | 2.28 | 2.6 | 3.0 | 2.7 | 2.5 |
| Continent | [1990-1995] | [1995-2000] | [2000-2005] | [2005-2010] | [2010-2015] | [2015-2020] | [1990-2020] | [1990-2000] | [2000-2010] | [2010-2020] |
| Africa | 0.56 | 1.82 | 3.34 | 2.15 | 2.42 | 3.04 | 2.2 | 1.2 | 2.7 | 2.6 |
| Latin-America | 3.48 | 6.32 | 6.22 | 4.03 | 3.27 | 4.81 | 4.7 | 4.9 | 5.1 | 4.8 |
| Asia-Oceania | 2.89 | 6.22 | 4.48 | 3.60 | 2.96 | 3.42 | 3.9 | 4.6 | 4.0 | 3.7 |
| Total | 6.93 | 14.37 | 14.04 | 9.78 | 8.66 | 11.27 | 10.8 | 10.6 | 11.9 | 11.1 |

b) annual loss due to deforestation (with or without prior degradation)
|  | Total deforestation on an annual basis by period (Mha) |  |  |  |  |  |  |  |  |  |
| --- | --- | --- | --- | --- | --- | --- | --- | --- | --- | --- |
| Sub-region | [1990-1995] | [1995-2000] | [2000-2005] | [2005-2010] | [2010-2015] | [2015-2020] | [1990-2020] | [1990-2000] | [2000-2010] | [2010-2020] |
| West-Africa | 0.06 | 0.19 | 0.71 | 0.48 | 0.45 | 0.60 | 0.4 | 0.1 | 0.6 | 0.5 |
| Central-Africa | 0.17 | 0.74 | 1.18 | 0.76 | 0.90 | 0.91 | 0.8 | 0.5 | 1.0 | 0.9 |
| South-East Africa | 0.10 | 0.45 | 0.38 | 0.18 | 0.18 | 0.14 | 0.2 | 0.3 | 0.3 | 0.2 |
| Central-America | 0.34 | 0.76 | 0.45 | 0.29 | 0.26 | 0.37 | 0.4 | 0.6 | 0.4 | 0.3 |
| South-America | 2.57 | 4.44 | 4.35 | 2.54 | 1.73 | 2.16 | 3.0 | 3.5 | 3.4 | 1.9 |
| Continental SE Asia | 0.84 | 1.55 | 1.13 | 0.74 | 0.56 | 0.54 | 0.9 | 1.2 | 0.9 | 0.6 |
| Insular SE Asia | 1.05 | 2.83 | 1.91 | 1.53 | 1.27 | 0.94 | 1.6 | 1.9 | 1.7 | 1.1 |
| Continent | [1990-1995] | [1995-2000] | [2000-2005] | [2005-2010] | [2010-2015] | [2015-2020] | [1990-2020] | [1990-2000] | [2000-2010] | [2010-2020] |
| Africa | 0.33 | 1.38 | 2.26 | 1.42 | 1.54 | 1.65 | 1.43 | 0.86 | 1.84 | 1.59 |
| Latin-America | 2.91 | 5.21 | 4.80 | 2.83 | 1.99 | 2.53 | 3.38 | 4.06 | 3.82 | 2.26 |
| Asia-Oceania | 1.89 | 4.38 | 3.04 | 2.27 | 1.83 | 1.48 | 2.48 | 3.14 | 2.65 | 1.65 |
| Total | 5.14 | 10.97 | 10.10 | 6.52 | 5.35 | 5.66 | 7.29 | 8.06 | 8.31 | 5.51 |

c) annual loss due to degradation (followed or not by deforestation)
|  | Total degradation on an annual basis by period (Mha) |  |  |  |  |  |  |  |  |  |
| --- | --- | --- | --- | --- | --- | --- | --- | --- | --- | --- |
| Sub-region | [1990-1995] | [1995-2000] | [2000-2005] | [2005-2010] | [2010-2015] | [2015-2020] | [1990-2020] | [1990-2000] | [2000-2010] | [2010-2020] |
| West-Africa | 0.08 | 0.16 | 0.87 | 0.50 | 0.35 | 0.61 | 0.4 | 0.1 | 0.7 | 0.5 |
| Central-Africa | 0.28 | 0.83 | 1.40 | 0.91 | 0.92 | 1.24 | 0.9 | 0.6 | 1.2 | 1.1 |
| South-East Africa | 0.09 | 0.27 | 0.28 | 0.14 | 0.13 | 0.13 | 0.2 | 0.2 | 0.2 | 0.1 |
| Central-America | 0.29 | 0.64 | 0.49 | 0.34 | 0.27 | 0.45 | 0.4 | 0.5 | 0.4 | 0.4 |
| South-America | 1.25 | 2.29 | 2.61 | 1.83 | 1.59 | 2.54 | 2.0 | 1.8 | 2.2 | 2.1 |
| Continental SE Asia | 0.88 | 1.23 | 1.03 | 0.81 | 0.49 | 0.78 | 0.9 | 1.1 | 0.9 | 0.6 |
| Insular SE Asia | 1.16 | 2.80 | 1.98 | 1.39 | 1.03 | 1.65 | 1.7 | 2.0 | 1.7 | 1.3 |
| Continent | [1990-1995] | [1995-2000] | [2000-2005] | [2005-2010] | [2010-2015] | [2015-2020] | [1990-2020] | [1990-2000] | [2000-2010] | [2010-2020] |
| Africa | 0.45 | 1.26 | 2.56 | 1.55 | 1.40 | 1.98 | 1.53 | 0.86 | 2.05 | 1.69 |
| Latin-America | 1.54 | 2.93 | 3.10 | 2.16 | 1.85 | 2.99 | 2.43 | 2.24 | 2.63 | 2.42 |
| Asia-Oceania | 2.04 | 4.03 | 3.00 | 2.21 | 1.51 | 2.43 | 2.54 | 3.04 | 2.61 | 1.97 |
| **Total** | **4.03** | **8.23** | **8.66** | **5.92** | **4.77** | **7.40** | **6.50** | **6.13** | **7.29** | **6.09** |

d) annual loss due to direct deforestation (without prior degradation)
|  | Annual direct deforestation by period (Mha) |  |  |  |  |  |  |  |  |  |
| --- | --- | --- | --- | --- | --- | --- | --- | --- | --- | --- |
| Sub-region | [1990-1995] | [1995-2000] | [2000-2005] | [2005-2010] | [2010-2015] | [2015-2020] | [1990-2020] | [1990-2000] | [2000-2010] | [2010-2020] |
| West-Africa | 0.02 | 0.08 | 0.21 | 0.19 | 0.32 | 0.40 | 0.2 | 0.0 | 0.2 | 0.4 |
| Central-Africa | 0.05 | 0.24 | 0.38 | 0.31 | 0.57 | 0.56 | 0.4 | 0.1 | 0.3 | 0.6 |
| South-East Africa | 0.05 | 0.24 | 0.19 | 0.10 | 0.13 | 0.10 | 0.1 | 0.1 | 0.1 | 0.1 |
| Central-America | 0.16 | 0.34 | 0.17 | 0.13 | 0.16 | 0.22 | 0.2 | 0.2 | 0.2 | 0.2 |
| South-America | 1.78 | 3.05 | 2.95 | 1.73 | 1.26 | 1.61 | 2.1 | 2.4 | 2.3 | 1.4 |
| Continental SE Asia | 0.33 | 0.66 | 0.48 | 0.36 | 0.41 | 0.36 | 0.4 | 0.5 | 0.4 | 0.4 |
| Insular SE Asia | 0.52 | 1.53 | 1.00 | 1.03 | 1.04 | 0.62 | 1.0 | 1.0 | 1.0 | 0.8 |
| Continent | [1990-1995] | [1995-2000] | [2000-2005] | [2005-2010] | [2010-2015] | [2015-2020] | [1990-2020] | [1990-2000] | [2000-2010] | [2010-2020] |
| Africa | 0.11 | 0.56 | 0.78 | 0.60 | 1.02 | 1.06 | 0.69 | 0.34 | 0.69 | 1.04 |
| Latin-America | 1.94 | 3.39 | 3.12 | 1.86 | 1.42 | 1.82 | 2.26 | 2.66 | 2.49 | 1.62 |
| Asia-Oceania | 0.85 | 2.19 | 1.48 | 1.39 | 1.45 | 0.99 | 1.39 | 1.52 | 1.44 | 1.22 |
| **Total** | **2.90** | **6.14** | **5.38** | **3.86** | **3.89** | **3.87** | **4.34** | **4.52** | **4.62** | **3.88** |

e) annual degradation before deforestation
|  | Annual degradation before deforestation by period (Mha) |  |  |  |  |  |  |  |  |  |
| --- | --- | --- | --- | --- | --- | --- | --- | --- | --- | --- |
| Sub-region | [1990-1995] | [1995-2000] | [2000-2005] | [2005-2010] | [2010-2015] | [2015-2020] | [1990-2020] | [1990-2000] | [2000-2010] | [2010-2020] |
| West-Africa | 0.04 | 0.11 | 0.50 | 0.28 | 0.14 | 0.20 | 0.2 | 0.1 | 0.4 | 0.2 |
| Central-Africa | 0.12 | 0.50 | 0.79 | 0.45 | 0.33 | 0.35 | 0.4 | 0.3 | 0.6 | 0.3 |
| South-East Africa | 0.06 | 0.21 | 0.19 | 0.08 | 0.05 | 0.04 | 0.1 | 0.1 | 0.1 | 0.0 |
| Central-America | 0.18 | 0.42 | 0.28 | 0.16 | 0.10 | 0.16 | 0.2 | 0.3 | 0.2 | 0.1 |
| South-America | 0.79 | 1.40 | 1.40 | 0.81 | 0.47 | 0.55 | 0.9 | 1.1 | 1.1 | 0.5 |
| Continental SE Asia | 0.51 | 0.89 | 0.65 | 0.38 | 0.15 | 0.18 | 0.5 | 0.7 | 0.5 | 0.2 |
| Insular SE Asia | 0.53 | 1.31 | 0.91 | 0.50 | 0.23 | 0.31 | 0.6 | 0.9 | 0.7 | 0.3 |
| Continent | [1990-1995] | [1995-2000] | [2000-2005] | [2005-2010] | [2010-2015] | [2015-2020] | [1990-2020] | [1990-2000] | [2000-2010] | [2010-2020] |
| Africa | 0.22 | 0.82 | 1.48 | 0.82 | 0.52 | 0.59 | 0.74 | 0.52 | 1.15 | 0.55 |
| Latin-America | 0.98 | 1.82 | 1.68 | 0.97 | 0.57 | 0.71 | 1.12 | 1.40 | 1.33 | 0.64 |
| Asia-Oceania | 1.05 | 2.19 | 1.56 | 0.88 | 0.38 | 0.49 | 1.09 | 1.62 | 1.22 | 0.44 |
| **Total** | **2.24** | **4.83** | **4.72** | **2.66** | **1.47** | **1.79** | **2.95** | **3.54** | **3.69** | **1.63** |

f) annual deforestation followed by a regrowth
| Total deforestation followed by regrowth on an annual basis by period (Mha) |  |  |  |  |  |  |  |  |  |  |
| --- | --- | --- | --- | --- | --- | --- | --- | --- | --- | --- |
| Sub-region | [1990-1995[ | [1995-2000[ | [2000-2005[ | [2005-2010[ | [2010-2015[ | [2015-2020[ | [1990-2020[ | [1990-2000[ | [2000-2010[ | [2010-2020[ |
| West-Africa | 0.00 | 0.00 | 0.01 | 0.04 | 0.06 | 0.03 | 0.0 | 0.0 | 0.0 | 0.0 |
| Central-Africa | 0.02 | 0.04 | 0.07 | 0.10 | 0.13 | 0.06 | 0.1 | 0.0 | 0.1 | 0.1 |
| South-East Africa | 0.00 | 0.01 | 0.02 | 0.03 | 0.03 | 0.01 | 0.0 | 0.0 | 0.0 | 0.0 |
| Central-America | 0.05 | 0.09 | 0.07 | 0.07 | 0.05 | 0.02 | 0.1 | 0.1 | 0.1 | 0.0 |
| South-America | 0.21 | 0.40 | 0.50 | 0.49 | 0.37 | 0.20 | 0.4 | 0.3 | 0.5 | 0.3 |
| Continental SE Asia | 0.10 | 0.20 | 0.24 | 0.23 | 0.14 | 0.06 | 0.2 | 0.2 | 0.2 | 0.1 |
| Insular SE Asia | 0.11 | 0.33 | 0.44 | 0.42 | 0.30 | 0.15 | 0.3 | 0.2 | 0.4 | 0.2 |
| Continent | [1990-1995[ | [1995-2000[ | [2000-2005[ | [2005-2010[ | [2010-2015[ | [2015-2020[ | [1990-2020[ | [1990-2000[ | [2000-2010[ | [2010-2020[ |
| Africa | 0.02 | 0.06 | 0.10 | 0.17 | 0.21 | 0.10 | 0.11 | 0.04 | 0.13 | 0.15 |
| Latin-America | 0.26 | 0.48 | 0.58 | 0.56 | 0.43 | 0.22 | 0.42 | 0.37 | 0.57 | 0.32 |
| Asia-Oceania | 0.21 | 0.53 | 0.68 | 0.65 | 0.44 | 0.21 | 0.45 | 0.37 | 0.66 | 0.32 |
| Total | 0.50 | 1.06 | 1.36 | 1.37 | 1.08 | 0.53 | 0.98 | 0.78 | 1.37 | 0.80 |

g) average percentage of invalid observations over the TMF domain per period and per year
| Average % of Invalid observations (over the total forest domain, per period) |  |  |  |  |  |  |  |
| --- | --- | --- | --- | --- | --- | --- | --- |
| Sub-region | [1982-1990[ | [1990-1995[ | [1995-2000[ | [2000-2005[ | [2005-2010[ | [2010-2015[ | [2015-2020[ |
| West-Africa | 98.1 | 87.1 | 82.5 | 40.0 | 2.8 | 0.4 | 0.0 |
| Central-Africa | 99.4 | 94.7 | 80.8 | 37.2 | 6.7 | 2.9 | 0.4 |
| South-East Africa | 97.9 | 80.8 | 20.7 | 3.9 | 1.3 | 0.8 | 0.1 |
| Central-America | 50.1 | 12.4 | 4.3 | 1.2 | 0.4 | 0.1 | 0.0 |
| South-America | 30.5 | 1.2 | 0.6 | 0.2 | 0.0 | 0.0 | 0.0 |
| Continental SE Asia | 54.5 | 14.6 | 1.3 | 0.4 | 0.0 | 0.0 | 0.0 |
| Insular SE Asia | 34.3 | 16.3 | 4.2 | 0.8 | 0.1 | 0.1 | 0.0 |
| Continent | [1982-1990[ | [1990-1995[ | [1995-2000[ | [2000-2005[ | [2005-2010[ | [2010-2015[ | [2015-2020[ |
| Africa | 99.1 | 92.9 | 77.6 | 35.7 | 5.9 | 2.4 | 0.3 |
| Latin-America | 31.3 | 1.6 | 0.8 | 0.2 | 0.1 | 0.0 | 0.0 |
| Asia-Oceania | 29.5 | 12.0 | 3.7 | 1.3 | 0.3 | 0.2 | 0.0 |
| Total | 21.2 | 31.9 | 23.1 | 10.1 | 2.3 | 0.8 | 0.1 |

| Average % of Invalid observations (over the total forest domain, per year) |  |  |  |  |  |  |  |  |
| --- | --- | --- | --- | --- | --- | --- | --- | --- |
| Sub-region | 1982 | 1990 | 1995 | 2000 | 2005 | 2010 | 2015 | 2019 |
| West-Africa | 100.0 | 90.8 | 84.3 | 71.8 | 5.9 | 0.7 | 0.2 | 0.0 |
| Central-Africa | 100.0 | 97.7 | 87.6 | 67.8 | 10.2 | 3.7 | 1.9 | 0.0 |
| South-East Africa | 100.0 | 92.1 | 41.5 | 8.4 | 1.5 | 1.0 | 0.5 | 0.0 |
| Central-America | 69.4 | 17.0 | 5.8 | 1.8 | 0.7 | 0.2 | 0.0 | 0.0 |
| South-America | 55.6 | 1.7 | 0.8 | 0.3 | 0.1 | 0.0 | 0.0 | 0.0 |
| Continental SE Asia | 55.2 | 49.1 | 2.0 | 1.0 | 0.0 | 0.0 | 0.0 | 0.0 |
| Insular SE Asia | 34.6 | 31.2 | 6.5 | 1.9 | 0.2 | 0.1 | 0.0 | 0.0 |
| Continent | 1982 | 1990 | 1995 | 2000 | 2005 | 2010 | 2015 | 2019 |
| Africa | 100.0 | 93.5 | 71.2 | 49.3 | 5.9 | 1.8 | 0.9 | 0.0 |
| Latin-America | 62.5 | 9.4 | 3.3 | 1.0 | 0.4 | 0.1 | 0.0 | 0.0 |
| Asia-Oceania | 44.9 | 40.2 | 4.3 | 1.4 | 0.1 | 0.1 | 0.0 | 0.0 |
| Total | 69.1 | 47.7 | 26.2 | 17.3 | 2.1 | 0.7 | 0.3 | 0.0 |

**Table 3.** Total areas and proportions of tropical moist forest disturbances (deforestation without regrowth, regrowth after deforestation, forest degradation) and reforestation areas (initially other land cover) over the period 1990-2020 for each sub-region and continent (areas in million ha and proportions in percentage).

|  | Disturbed areas (Mha) |  |  |  | % of Tot disturbances |  |  | % of Undisturbed Forest in 1990 |  |  |  |  |
| --- | --- | --- | --- | --- | --- | --- | --- | --- | --- | --- | --- | --- |
| Sub-region | Deforestation | Regrowth | Degradation | Total | Deforestation | Regrowth | Degradation | Deforestation | Regrowth | Degradation | Total | Reforestation<br>(from other LC) |
| West-Africa | 11.8 | 0.7 | 6.5 | 19.0 | 61.9 | 3.7 | 34.5 | 34.0 | 2.0 | 18.9 | 55.0 | 0.4 |
| Central-Africa | 21.2 | 2.1 | 15.1 | 38.4 | 55.2 | 5.4 | 39.4 | 9.5 | 0.9 | 6.8 | 17.2 | 1.1 |
| South-East Africa | 6.7 | 0.5 | 2.1 | 9.3 | 71.9 | 5.7 | 22.4 | 42.5 | 3.4 | 13.3 | 59.2 | 0.2 |
| Central-America | 10.6 | 1.8 | 5.9 | 18.3 | 58.0 | 9.7 | 32.3 | 30.7 | 5.1 | 17.1 | 53.0 | 0.7 |
| South-America | 78.1 | 10.9 | 33.4 | 122.4 | 63.8 | 8.9 | 27.3 | 11.6 | 1.6 | 5.0 | 18.2 | 4.0 |
| Continental SE Asia | 22.0 | 4.9 | 12.2 | 39.1 | 56.2 | 12.4 | 31.4 | 30.0 | 6.6 | 16.7 | 53.3 | 2.0 |
| Insular SE Asia | 38.9 | 8.7 | 31.1 | 78.8 | 49.4 | 11.1 | 39.5 | 16.4 | 3.7 | 13.1 | 33.1 | 1.6 |
| Continent | Deforestation | Regrowth | Degradation | Total | Deforestation | Regrowth | Degradation | Deforestation | Regrowth | Degradation | Total | Reforestation<br>(from other LC) |
| Africa | 39.6 | 3.3 | 23.8 | 66.7 | 59.4 | 4.9 | 35.6 | 14.5 | 1.2 | 8.7 | 24.4 | 1.6 |
| Latin-America | 88.7 | 12.6 | 39.3 | 140.7 | 63.1 | 9.0 | 27.9 | 12.6 | 1.8 | 5.6 | 19.9 | 4.7 |
| Asia-Oceania | 60.9 | 13.6 | 43.4 | 117.8 | 51.7 | 11.5 | 36.8 | 19.6 | 4.4 | 13.9 | 37.9 | 3.6 |
| Total | 189.2 | 29.5 | 106.5 | 325.2 | 58.2 | 9.1 | 32.7 | 14.9 | 2.3 | 8.4 | 25.7 | 10.0 |

**Fig. 3.**
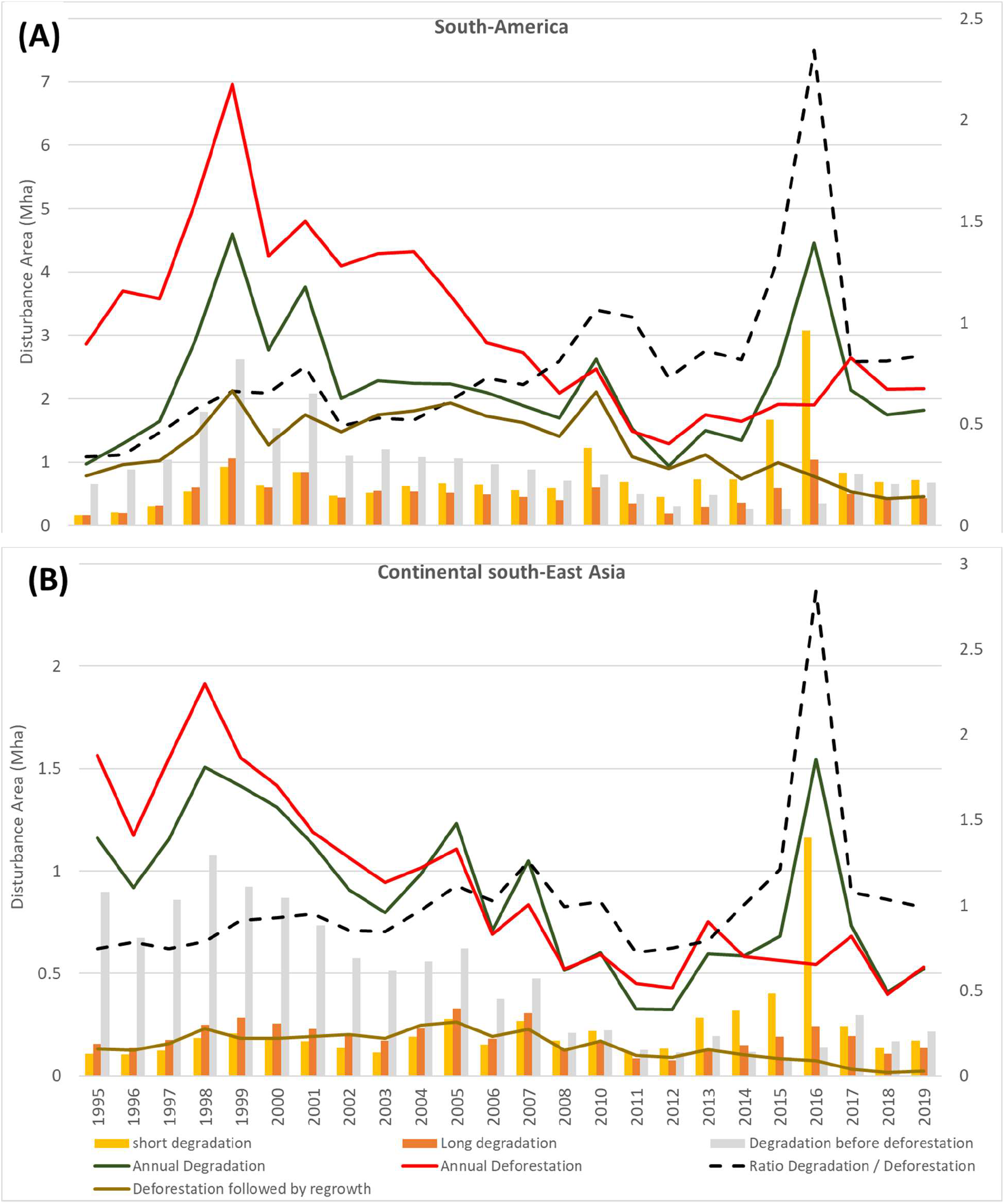
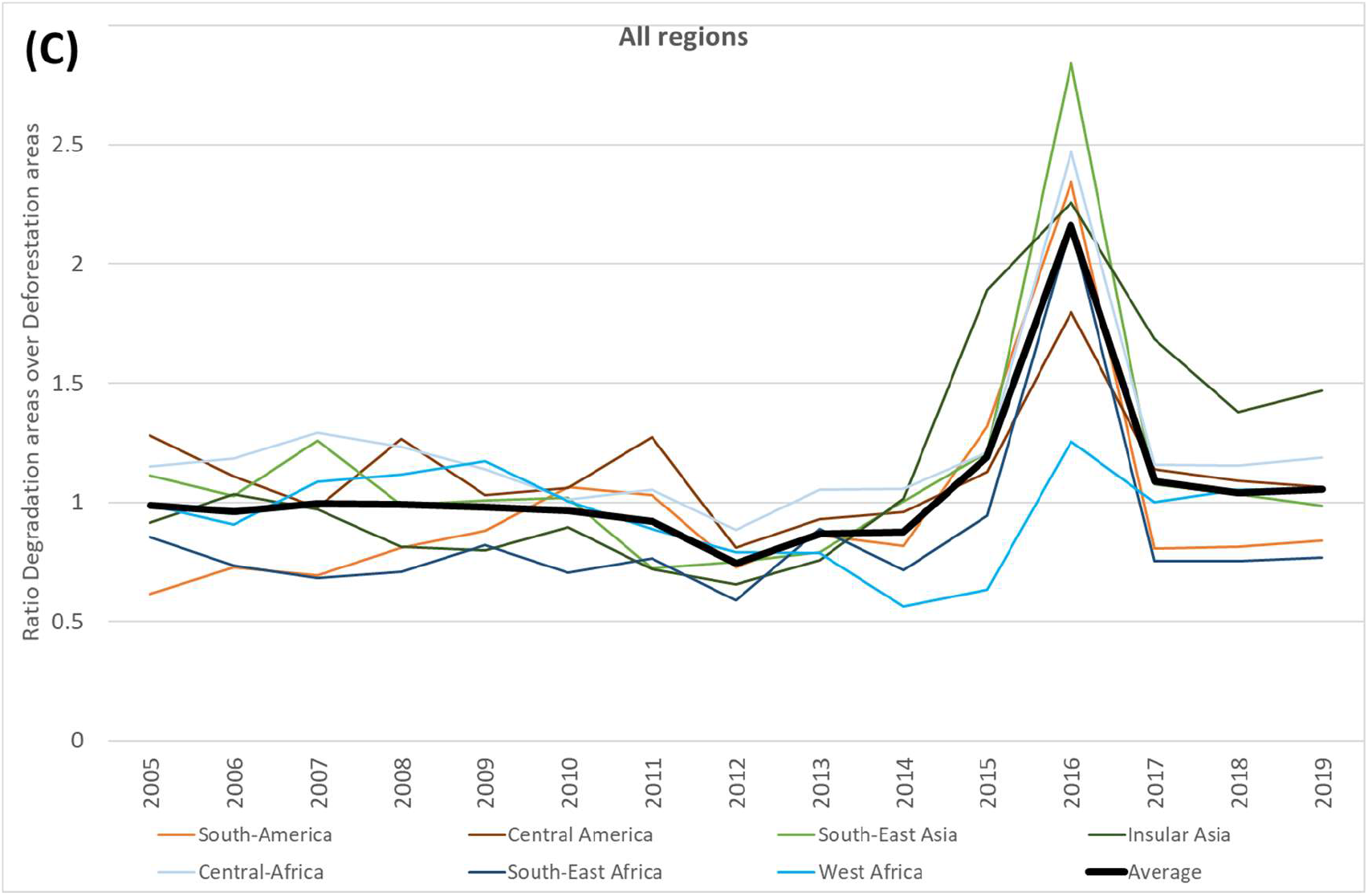
Evolution of annual deforestation and degradation (A) over the last 25 years in South America, and (B) in continental South-East Asia regions, and (C) over the last 15 years in all regions.

**Fig. 4.**
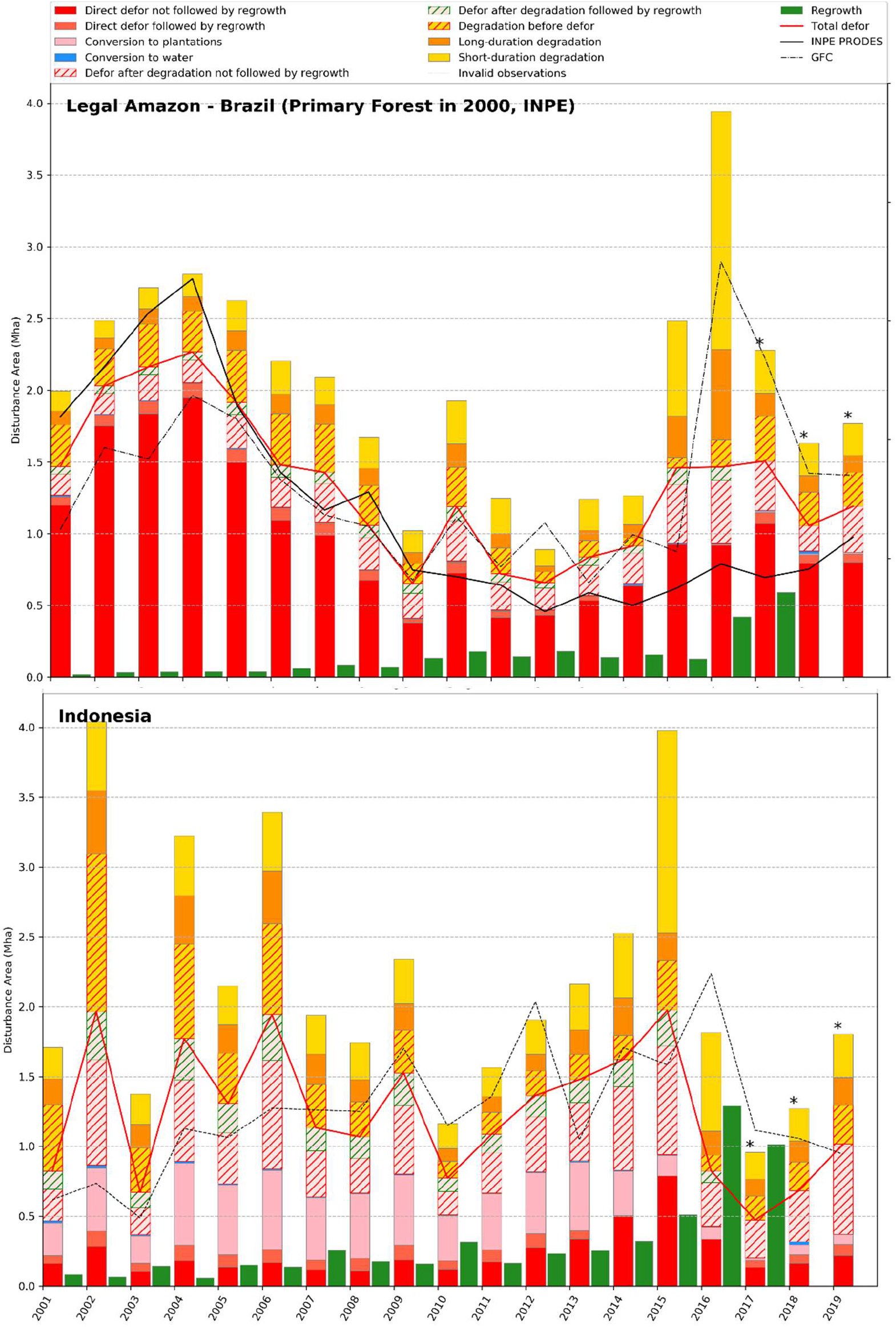
Dynamics of annual disturbed areas from 2001 to 2019 for (A) the Amazônia Legal region of Brazil within the primary forest extent in 2000 from INPE and (B) Indonesia using the entire TMF extent (undisturbed and degraded) in 2000. (x-axis in years and y-axis in million ha) in comparison with GFC loss and the PRODES data for the Amazônia Legal region of Brazil. * The average proportions of disturbance types within total disturbances over the period 2005-2014 is used to distribute the disturbance types for years 2017 to 2019.

Our results stress the paramount importance of (i) integrating measures for reducing degradation in forest conservation and climate mitigation programs, and (ii) considering forest degradation as risk factor of deforestation and as an indicator of climate change and climate oscillations. We anticipate that a better knowledge of forest degradation processes and its resulting fragmentation will help to assess accurately the anthropogenic impact on the tropical ecosystem services and the effects on biosphere-atmosphere-hydrosphere feedbacks. Future policies will have to account for this finding.

### Main results on deforestation and post-deforestation regrowths

Deforestation in TMF cover is documented in an unprecedented comprehensive manner: (i) by covering a 30-year period of analysis, (ii) by mapping deforestation occurring after degradation and deforestation followed by a regrowth, (iii) by identifying specific forest conversion to commodities or water (**Figs. 2G** **and** **2I**), (iv) by including changes within the mangroves (**Fig. 2A**), and (v) by documenting each deforestation event at the pixel level by its timing (date and duration), intensity, recurrence and when appropriate, start date and duration of post-disturbance regrowth.

Overall, 17.2% of the initial TMF area (i.e. 207.4 over 1267.1 million ha), have disappeared since 1990, down to 1059.6 million ha of TMF in January 2020 (**Tables 1**, **2** **and** **3**). We report a rate of gross loss of TMF area for the entire pan-tropical region varying from 5.5 to 7.7 million ha / year with the period (**Table 4**). Comparison with previous studies results in the following outcomes:

i. Estimations reported by FAO national statistics (**30**) and the sample-based estimations from Tyukavina et al. (**31**) for the natural tropical forest – that includes both moist and dry forest types - are higher by 0.9% and 27% respectively, compared to our TMF deforestation rates (excluding the conversion to tree plantations to get closer to the natural forest definition of these two studies) for the same period (**Table 4**). At the continental scale, Tyukavina et al. (**31**) shows lower estimates than our study for Africa (−23%) and for Asia (−4%), and higher estimates for Latin America (+16%).
ii. Comparison with GFC loss (**24**) (see Section on the Comparison with the GFC dataset and **Fig. 4**) shows a lower deforestation rate (−33%) compared to our study for the period 2000-2012 over the same forest extent (using our TMF extent for the year 2000) (**Table 4**). Underestimation of GFC loss has been documented by previous studies (**31, 33**). Tyukavina et al. (**31**) reported an underestimation of GFC loss of 19.4% considering the entire forest cover (moist and deciduous) loss during the period 2001-2012, with a larger underestimation for Africa (−39.4%) compared to other continents (−13% for Latin America and −5.7% for Asia). The ranking of this underestimation by continent is consistent with the ranking observed in our study (first Africa, second Latin America and third Asia). The differences with GFC loss are explained by three specific assets of our approach: (i) the use of single-date images enabling the detection of short-duration disturbance events (i.e. visible only during a few weeks from space) compared to the use of annual syntheses, (ii) a dedicated algorithm for TMF enabling the monitoring of seven forest cover change classes compared to the global monitoring of forest clearance, and (iii) a cloud masking and quality control optimized for equatorial regions enabling a more comprehensive analysis of the Landsat archive.
iii. Comparison with the Brazilian PRODES data (**29**) using their primary forest extent (**Fig. 4**) shows a similar decrease of annual deforestation rates between the 2000’s and the last decade that can be related to a set of economic and public policy actions (**28**). Differences in the deforestation rates are observed (i) during the period 2001-2004 with a higher deforestation rate for PRODES (2.32 million ha/year) compared to our study (2 million ha/year) and to GFC loss (1.53 million ha/year), and (ii) in the last ten years with a lower average deforestation rate for PRODES compared to our study and GFC loss (0.67, 1.1 and 1.34 million ha/year respectively) (**Table 4**). These differences are accentuated in the last five years (0.77, 1.33, and 1.76 million ha/year respectively). Discrepancies in area estimates between our product and the PRODES data are explained by (i) difference in minimum mapping units (0.09 ha compared to 6.25 ha in PRODES), and (ii) impacts of strong fires that are captured in our study (deforestation followed by forest regrowth) and in GFC loss but are discarded in the PRODES approach (because not considered as deforestation).

**Table 4.** Comparison of estimates of annual deforested areas (in million ha / year) from previous studies and our study, over the tropical belt, over the three continents and Brazil.

| Source |  | Hansen et al. 2013 |  | Tyukavina et al. 2015 | Keenan et al. 2015 | PRODES-INPE | This Study |  |  |
| --- | --- | --- | --- | --- | --- | --- | --- | --- | --- |
| Forest extent |  | Whole TMF (undisturbed and degraded) | Primary forest from INPE | Natural forests * | All tropical forests (evergreen & deciduous) | Primary forest | Tropical moist forest | TMF excluding the tree plantations | Primary forest from INPE |
| Pan-tropical region | 2001-2010 | 4.67 |  |  | 7.24 |  | 7.72 |  |  |
|  | 2001-2012 | 4.80 |  | 6.5 ± 0.7 |  |  | 7.19 | 6.44 |  |
|  | 2001-2015 | 5.07 |  |  | 6.66 |  | 6.95 |  |  |
|  | 2010-2019 | 6.87 |  |  |  |  | 5.51 |  |  |
|  | 2001-2019 | 5.79 |  |  |  |  | 6.66 |  |  |
| Africa | 2001-2012 | 0.73 |  | 1.21 ± 0.4 |  |  | 1.60 | 1.57 |  |
|  | 2001-2019 | 1.28 |  |  |  |  | 1.64 |  |  |
| Latin America | 2001-2012 | 2.19 |  | 3.7 ± 0.5 |  |  | 3.25 | 3.19 |  |
|  | 2001-2019 | 2.41 |  |  |  |  | 2.93 |  |  |
| Asia - Oceania | 2001-2012 | 1.89 |  | 1.6 ± 0.4 |  |  | 2.34 | 1.67 |  |
|  | 2001-2019 | 2.10 |  |  |  |  | 2.09 |  |  |
| Brazil | 2001-2010 | 1.61 | 1.35 |  |  | 1.65 | 2.55 |  | 1.57 |
|  | 2001-2012 | 1.54 | 1.26 | 2.1 ± 0.3 |  | 1.47 | 2.32 | 2.27 | 1.42 |
|  | 2010-2019 | 1.64 | 1.34 |  |  | 0.67 | 1.63 |  | 1.04 |
|  | 2001-2019 | 1.64 | 1.35 |  |  | 1.19 | 2.10 |  | 1.31 |

This study documents – in an unprecedented manner - the extent and age of post-deforestation regrowths (young secondary forests that are regenerating after human or natural disturbance) for the entire pan-tropical domain. These secondary forests grow rapidly in tropical moist conditions and absorb large amounts of carbon, whereas they were poorly documented. We show that 13.5% of the deforested areas (i.e. 29.5 Million ha) are regrowthing in a subsequent stage, with 33% of these secondary forests aged more than 10 years at the end of 2019 (**Table 3**). The proportion of secondary forests within the total deforestation is higher in Asia (18.3%) compared to Latin America (12.3%) and Africa (7.9%). The disturbance events followed by a *forest regrowth* are including intense fires and these are accentuated by drought conditions. This is well visible for South America (**Fig. 3**) for years 1997-1998 and 2010. Additionally, 10 Million ha are characterized as evergreen vegetation regrowth of areas initially classified as non-forest cover, i.e. that can be considered as forestation (i.e. afforestation and reforestation) aged of more than 10 years.

This study confirms that most of the deforestation caused by the expansion of oil palm and rubber and assigned to the commodity classes in our study (see Supplementary Text on ancillary datasets, **Figs. 2I** **and** **3B**, Fig. S11 and table S6) is concentrated in Asia with 18.3 million ha (representing 86% of the entire TMF conversion to plantations), and more specifically in Indonesia (57.4%) and Malaysia (23.8%).

### Deforestation and degradation trends

The evolution of the deforestation and degradation over the last three decades show the highest peaks of annual disturbances in Latin America and Southeast Asia during the period 1995-2000 with 6.3 million ha/year and 6.2 million ha/year respectively. The ENSO of 1997-1998 may - at least partially - explain these peaks of forest disturbances, in particular for Indonesia and Brazil where such peaks are manifest in the annual change trends with the highest proportion of degradation events over the total disturbance areas (**Figs. 3** **and** **5**, fig. S11). Between 2000-2004 and 2015-2019, the disturbance rates decreased by half in South-America and by 45% in South-East Africa and continental South-Est Asia. Brazil - that accounts for 29% of the remaining world’s TMF - largely contributed to this reduction (from 4.3 million ha/year down to 2.1 million ha/year) (**Figs. 3**, **4**, **and** **5**, **Table 4**, table S6 and fig. S11).

**Fig. 5.**
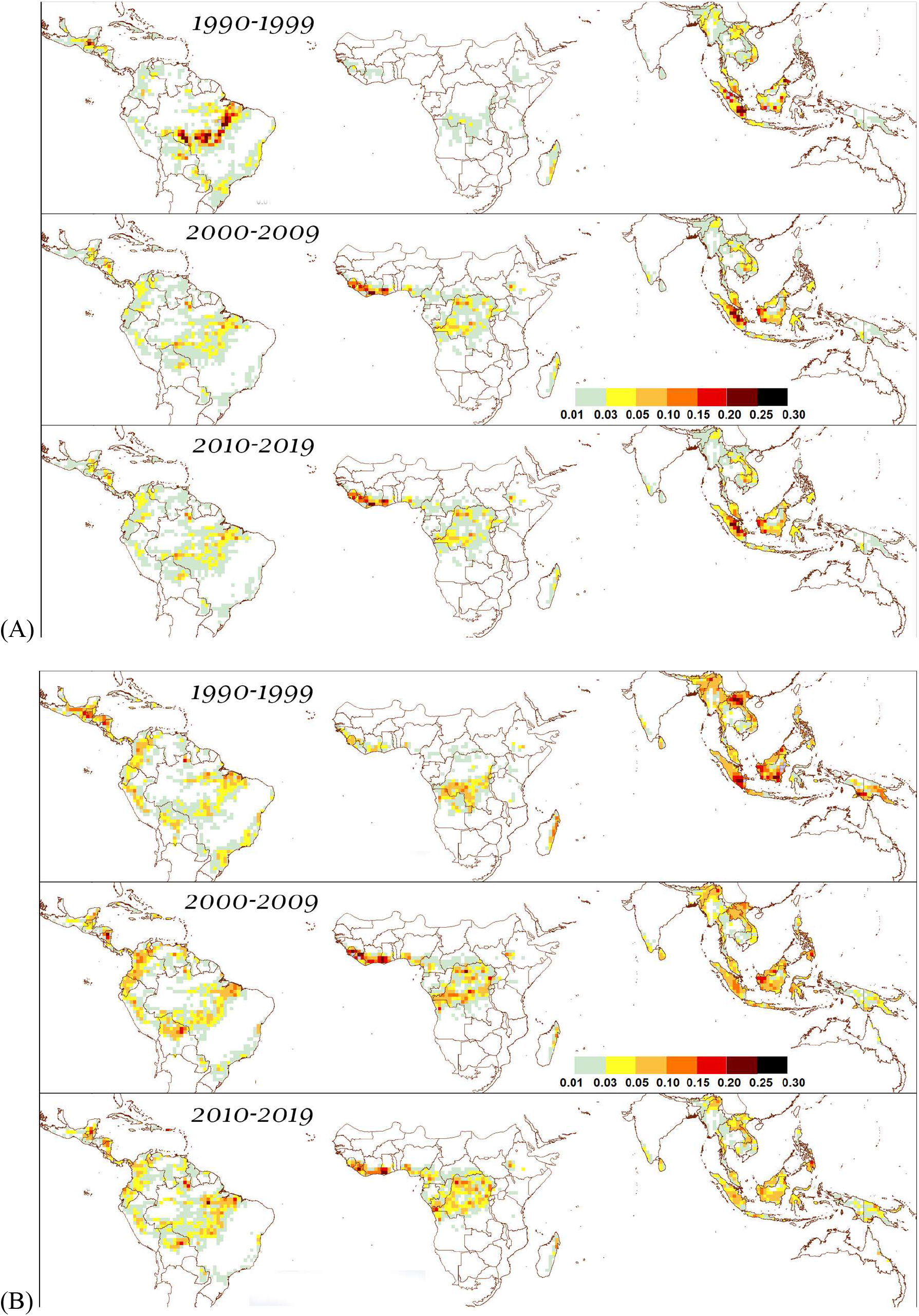
Evolution of hotspots of deforestation (A) and degradation (B) during the last three decades (total deforested or degraded area per box of 1° latitude × 1° longitude size – scale in million ha).

In the recent years, our study shows a dramatic increase of disturbances rates (deforestation and degradation) (+ 2.1 million ha/year for the last 5 years compared to the period 2005-2014) to reach a level close to that of the early 2000s (**Tables 2** **and** **3**) with the highest increases observed in West Africa and Latin America (48% higher). Degradation is the main contributor of this recent increase (average increase of 38% whereas annual deforestation decreased by 5%) caused notably by specific climatic conditions in 2015-2016 (**29**) (**Figs. 3** **and** **5**). Asia-Oceania region shows a lower increase of degradation rate (31%) compared to Africa (34%) and Latin America (49%) and a much higher decrease of deforestation rate (28%) compared to Africa (5%) and Latin America (12%).

### Undisturbed TMF decline and projections

Since 1990, the extent of undisturbed TMF has declined by 23.9% with an average rate of loss of 10.8 million ha/year. The decline of undisturbed TMF is particularly dramatic for Ivory Coast (81.5% of their extent in 1990), Mexico (73.7%), Ghana (70.8%), Madagascar (69%), Vietnam (67.8%), Angola (67.1%), Nicaragua (65.8%), Lao People’s Democratic Republic (PDR) (65.1%), and India (63.9%) (table S6). If the average rates of the period 2010-2019 would remain constant over a short or medium term future (see Materials and Methods, fig. S12), undisturbed TMF would disappear by 2026-2029 in Ivory Coast and Ghana, by 2040 in Central America and Cambodia, by 2050 in Nigeria, Lao PDR, Madagascar and Angola, and by 2065 for all the countries of continental Southeast Asia and Malaysia. By 2050, 15 countries -including Malaysia (the 9^th^ country with the biggest TMF forest) - will lose more than 50% of their undisturbed forests (table S6).

## CONCLUSION

It is now possible to monitor deforestation and degradation in tropical moist forests consistently over a long historical period and at fine spatial resolution. The mapping of forest transition stages will allow to derive more targeted indicators to measure the achievements in forest, biodiversity, health and climate policy goals from local to international levels (**34**). Our study shows that tropical moist forests are disappearing at much faster rates than what was previously estimated and underlines the precursor role of forest degradation in this process. These results should alert decision makers on the pressing need to reinforce actions for preserving tropical forest, in particular by avoiding the first scar of degradation that is most likely leading to forest clearance later on.

## MATERIALS AND METHODS

### Study area and Forest types

Our study covers the tropical moist forests, which include the following *formations* (**35**): the lowland evergreen rain forest, the montane rain forest, the mangrove forest, the swamp forest, the tropical semi-evergreen rain forest, and the moist deciduous forest. Evergreenness varies from permanently evergreen to evergreen seasonal (mostly evergreen but with individual trees that may lose their leaves), semi-evergreen seasonal (up to about one third of the top canopy can be deciduous, though not necessarily leafless at the same time), and moist deciduous (dominant deciduous species with evergreen secondary canopy layer).

We do not intent to map specifically intact or primary forest as the Landsat observation period is too short to discriminate never-cut primary forest from second growth naturally recovered forest older than the observation period. However, by documenting all the disturbances observed over the last three decades, the remaining undisturbed TMF in 2019 is getting closer to the primary forest extent. Whereas our entire TMF - that includes undisturbed and degraded forests - in 1990 and 2019 are comparable, our undisturbed forest of 1990 and 2019 should be carefully compared.

Our study area covers the following Global Ecological Zones (**36**): ‘Tropical rainforest’, ‘Tropical moist forest’, ‘Tropical mountain system’ and ‘Tropical dry forest’ (fig. S13) and stops at the borders of China, Pakistan, Uruguay, and USA. The TMF are located mostly in the tropical moist and humid climatic domains but also include small areas of gallery forests in the tropical dry domain.

### Data

The Landsat archive is the only free and long-term satellite image record suited for analysing vegetation dynamics at fine spatial resolution. We used the entire L1T archive (orthorectified top of atmosphere reflectance) acquired between July 1982 and December 2019 from the following Landsat sensors: Thematic Mapper (TM) onboard Landsat 4 and 5, Enhanced Thematic Mapper-plus (ETM+) onboard Landsat 7 and the Operational Land Imager (OLI) onboard Landsat 8 (**23**, **37–39**). Landsat 4 was launched in July 1982 and collected images from its TM sensor until December 1993. Landsat 5 was launched in March 1984 and collected images until November 2011. Landsat 7 was launched in April 1999 and acquired images normally until May 2003 when the scan line corrector (SLC) failed (**40**). All Landsat 7 data acquired after the date of the SLC failure have been used in our analysis. Landsat 8 began operational imaging in April 2013.

The Landsat archive coverage presents large geographical and temporal unevenness (**37, 41**). The main reason for the limited availability of images for some regions is that Landsat 4 and 5 had no onboard data recorders, and links with data relay satellites failed over time; cover was therefore often limited to the line of sight of receiving stations (**39**). Commercial management of the programme from 1985 to the early 1990s led to data acquisitions being acquired mostly when pre-ordered (**37**). From 1999 onwards, the launch of Landsat 7 and its onboard data recording capabilities, associated with the continuation of the Landsat 5 acquisitions, considerably improved global coverage.

In the tropical regions, Africa is particularly affected by the limited availability of image acquisitions, especially in the first part of the archive. From a total of around 1 370 860 Landsat scenes that were available for our study area, only 265 098 scenes were located in Africa (in comparison, 573 589 and 532 173 scenes were respectively available in South America and Asia). The most critical area is located around the Gulf of Guinea, with an overall average number of valid observations (i.e. without clouds, hazes, sensor artefacts and geo-location issues) over the full archive (fig. S14) of fewer than 50 per location (pixel) and with the first valid observations starting mostly at the end of the 1990s (fig. S15). Small parts of Ecuador, Colombia, Salomon Islands and Papua New Guinea present a similar low number of total valid observations, often with an earlier first valid observation around the end of the 1980s. Apart from these regions, the first valid observation occurs mostly within periods 1982-1984, 1984-1986, or 1986-1988 for Latin America, Africa and Southeast Asia, respectively.

The average number of annual valid observations (fig. S16) shows a stepped increase during the 38-year period for the three continents, with two major jumps: in 1999 with the launch of Landsat 7, and in 2013 with the launch of Landsat 8. There is also a clear drop in 2012 for Southeast Asia and Latin America with the decommissioning of Landsat 5 in November 2011, and a small drop in 2003 as a consequence of the Landsat 7 SLC off issue. There are major differences between Africa and the two other continents: Africa has significantly fewer valid observations, in particular during the period 1982-1999, and a much larger increase in number of observations from 2013. The geographical unevenness of the first year of acquisition constrains the monitoring capability period. Our method accounts for this constraint notably by recording the effective duration of the archive at the pixel level (see next subsection).

Data quality issues affecting the Landsat collection were addressed by excluding pixels where (i) detector artefacts occur (manifested as random speckle or striping), (ii) one or more spectral bands are missing (typically occurring at image edges) or (iii) scene geo-location is inaccurate.

### Mapping method

In order to map the area dynamics (extent and changes) of the TMF over a long period, we developed an expert system that exploits the multispectral and multitemporal attributes of the Landsat archive to identify the main change trajectories over the last 3 decades and uses ancillary information to identify sub-classes of forest conversion (see Supplementary Text on Ancillary data). The inference engine of our system is a procedural sequential decision tree, where the expert knowledge is represented in the form of rules. Techniques for big data exploration and information extraction, namely visual analytics (**42**) and evidential reasoning (**43**), were used similarly to a recent study dedicated to global surface water mapping (**41**). The advantages of these techniques for remotely sensed data analysis are presented in this previous study (**41**), notably for accounting for uncertainty in data, guiding and informing the expert’s decisions, and incorporating image interpretation expertise and multiple data sources. The expert system was developed and operated in the Google Earth Engine (GEE) geospatial cloud computing platform (**22**).

The mapping method includes four main steps described hereafter: (i) single-date multi-spectral classification into three classes, (ii) analysis of trajectory of changes using the temporal information and production of a ‘transition’ map (with seven classes) (**Figs. 1** **and** **2**, figs. S2 to S7), (iii) identification of sub-classes of transition based on ancillary datasets (see Supplementary Text on Ancillary datasets) and visual interpretation, (iv) production of annual change maps (fig. S1).

In the first step, each image of the Landsat archive was analysed on a single-date basis (through a multi-spectral classification), whereas previous large-scale studies used annual syntheses or intra-annual statistics such as the mean and standard deviation of available Landsat observations (**44–50**). Classification of individual images is challenging but presents three main advantages: it allows (i) to capture the disturbance events that are visible only over a short period from space, such as logging activities, (ii) to record the precise timing of the disturbances and the number of *disruption observations*, and (iii) to detect the disturbance at an early stage, i.e. even if the disturbance is starting at the end of the year, it is detected and counted as a disturbance for this year whereas other approaches notably based on composites will detect the disturbance with a delay of one year. A *disruption observation* is defined here as an absence of tree foliage cover within a 0.09 ha size Landsat pixel. The number of *disruption observations* constitutes a proxy of disturbance intensity. Each pixel within a Landsat image was initially assigned through single-date multi-spectral classification to one of three following classes: (i) *potential moist forest cover*, (ii) *potential disruption*, and (iii) *invalid observation* (cloud, cloud shadow, haze and sensor issue).

The temporal sequence of classes (i) and (ii) was then used to determine the seven transition classes, described in the second step of the mapping approach. However, not all pixels could be unambiguously spectrally assigned to one of the three single-date classes because the multi-spectral cluster hulls of such classes are overlapping in the multidimensional feature-space. In cases of spectral confusion, evidential reasoning was used to guide class assignment by taking into consideration the temporal trajectory of single-date classifications, as spectral overlap between land cover types may occur only at specific periods of the year. For instance, pixels covered by deciduous forests, grassland or agriculture, may behave – from a spectral point of view – as *potential moist forest cover* during the humid seasons and as *potential disruptions* during the dry seasons, and, consequently, can be assigned to the *other land cover* transition class. Disturbed moist forests (degraded or deforested) are appearing as *potential moist forest cover* at the start of the archive and as *potential disruption* assignments later.

For the three initial classes (*potential moist forest cover*, *potential disruption*, and *invalid observation*), multispectral clusters were defined first by establishing a spectral library capturing the spectral signatures of the land cover types and atmosphere perturbations that are present over the pan-tropical belt and targeted for these three classes: (i) moist forest types, (ii) deciduous forest, logged areas, savannah, bare soil, irrigated and non-irrigated cropland, evergreen shrubland and water (for the *potential disruption* class) and (iii) clouds, haze, cloud shadows (for the *invalid observations*). A total sample of 38 326 sampled pixels belonging to 1 512 Landsat scenes (L5, L7 and L8), were labelled through visual interpretation. The HSV (*hue*, *saturation*, *value*) transformation of the spectral bands - well adapted for satellite image analysis (**41, 52**) - were used to complement the spectral library. These components were computed using a standard transformation (**52**) for the following Landsat band combination: short-wave infrared (SWIR2), near infrared (NIR) and red. The stability of *hue* to the impacts of atmospheric effect is particularly desirable for identifying *potential disruption* in the humid tropics. The sensitivity of *saturation* and *value* to atmospheric variability is mainly used to detect *invalid* observations (haze). *Value* is particularly useful for identifying cloud shadows. The thermal infrared band (TIR) was relevant to detect *invalid* observations (clouds, haze) and bare soil, and the Normalized Difference Water Index (NDWI) to identify irrigated areas. The information held in the spectral library was analyzed through visual analytics to extract equations describing class cluster hulls in the multidimensional feature-space (fig. S17). An exploratory data analysis tool designed in a previous study (**41**) was used to support the interactive analysis.

In the second step of the mapping approach, the temporal sequence of single-date classifications at pixel scale was analysed to first determine the initial extent of the TMF domain and then to identify the change trajectories from this initial forest extent (fig. S2). Long-term changes cannot be determined uniformly for the entire pan-tropical region because the observation record varies (see Data), e.g. the first year of observation (fig. S18) is c. 1982 for Brazil and c. 2000 along the Gulf of Guinea. We have addressed this geographic and temporal discontinuities of the Landsat archive by determining at the pixel level (i) a reference initial period (baseline) for mapping the initial TMF extent and (ii) a monitoring period for detecting the changes. The data gaps at the beginning of the archive were tackled by requiring a minimum period of four years with a minimum of three valid observations per year or a minimum of five years with two valid observations per year from the first available valid observation. Hence, lower is the annual number of valid observations, higher is the length of the initial period. This minimizes the risk of inclusion of non-forest cover types (such as agriculture) and deciduous forests in the baseline when there are few valid observations over a short period. In addition, we have reduced the commission errors in our baseline by accounting for possible confounding with commodities, wetlands, bamboo, and deciduous forest (see Supplementary Text on ancillary datasets and specific tropical forest types). From our initial TMF extent, we identified seven main transition classes (fig. S2) which are defined thereafter. The first year of the monitoring period (that follows the initial period) is represented at fig. S18; it starts at the earliest in year 1987 (mostly for South-America) and, for very limited cases, at the latest in 2016 (e.g. Gabon).

Although no ecosystem may be considered truly undisturbed, because some degree of human impact is present everywhere (**54**), we define the undisturbed moist forests (class 1) as tropical moist (evergreen or semi-evergreen) forest coverage without any disturbance (degradation or deforestation) observed over the Landsat historical record (see Section on the Study area and forest types). Our TMF baseline may include old *forest regrowth* (old secondary forests) or previously degraded forests forest as the Landsat observation period is too short to discriminate never-cut primary forest from second growth naturally recovered forest older than the observation period. This class includes two sub-classes of bamboo-dominated forest (class 1a) and undisturbed mangrove (class 1b).

A *deforested land* (class 2) is defined as a permanent conversion from moist forest cover to another land cover whereas a *degraded forest* (class 3) is defined as a moist forest cover where disturbances were observed over a short time period. Here we assumed that the duration of the disturbance (and consequently the period over which we detect the disturbance with satellite imagery) is a proxy of the disturbance impact, i.e. higher is the duration of the detected disturbance, higher is the impact on the forest, and higher is the risk to have a permanent conversion of the TMF. By considering short-term disturbances we include logging activities, fires and natural damaging events such as wind breaks and extreme dryness periods. Hence, we are getting closer to the most commonly accepted definition of the degradation (**54**) that considers a loss of productivity, a loss of biodiversity, unusual disturbances (droughts, blowdown), and a reduction of carbon storage.

The threshold applied on the duration parameter used to separate *degraded forests* from *deforested land* is based on our knowledge of the impacts of human activities and of natural or human-induced events such as fires. We identified empirically two levels of degradation: (class 3a) degradation with short-duration impacts (observed within a 1-year maximum duration), which includes the majority of logging activities, natural events and light fires, and (class 3b) degradation with long-duration impacts (between one and 2.5 years) which mainly corresponds to strong fires (burned forests). Most of the degradation (50%) are observed over less than six-month durations (fig. S19). All disturbance events for which the impacts were observed over more than 2.5 years (900 days) were considered as deforestation processes, with 68% of such deforestation events observed over more than five years. When a deforestation process is not followed by a regrowth period at least over the last 3 years, it is considered as a *Deforested land*. Deforested land are also characterized by the recurrence of disruptions, i.e. the ratio between the number of years with at least one *disruption observation* and the total number of years between the first and last *disruption observations*. This information allowed to discriminate deforestation without prior degradation from deforestation occurring after degradation, the second one having a lower recurrence due to the period without any disruption between the degradation and deforestation phases (see Supplementary Text on annual change dataset).

*For the recent degradation and deforestation* (class 4) that initiated in the last three years (after year 2016) and that cannot yet be attributed to a long-term conversion to a non-forest cover, owing to the limited historical period of observation, specific rules were applied. Within this class, we separated degradation from deforestation, by taking a duration of minimum 366 days for the years 2017-2018 and a threshold of 10 disruptions for the last year (2019) to consider a *deforested land*. A *forest regrowth* (class 5) is a two-phase transition from moist forest to (i) *deforested land* and then (ii) vegetative regrowth. A minimum 3-years duration of permanent moist forest cover presence is needed to classify a pixel as *forest regrowth* (to avoid confusion with agriculture). The *other land cover* (class 6) includes savannah, deciduous forest, agriculture, evergreen shrubland and non-vegetated cover.

Finally, the *Vegetation regrowth* (*class 7*) consists of a transition from other land cover to vegetation regrowth and includes two sub-classes of vegetation regrowth according to the age of regrowth (between 3 and 10 years, and between 10 and 20 years) and a transition class from water to vegetation regrowth.

The third mapping step allowed to identify three sub-classes from the *deforested land* class. We geographically assigned deforestation to the conversion from TMF to tree plantations - mainly oil palm and rubber (class 2a), water surface (discriminating permanent and seasonal water)-mainly due to new dams (class 2b), and other land cover - agriculture, infrastructures, etc. (class 2c) using ancillary spatial datasets completed by visual interpretation of high-resolution (HR) imagery (see Supplementary Text on ancillary data). Finally, we have re-assigned disturbances when detected within two geographically specific tropical forest formations: (i) the bamboo dominated forest, and (ii) the semi-deciduous transition tropical forest (Supplementary Text on specific tropical forest formations).

Each disturbed pixel (degraded forest, deforested land, or forest regrowth) is characterized by the timing and intensity of the observed disruption events. The start and end dates of the disturbance allows identifying in particular the timing of creation of new roads or of logging activities and the age of forest regrowth or degraded forests. Three decadal periods have been used in the transition map to identify age sub-classes of degradation and forest regrowth: (i) before 2000, (ii) within 2000-2009 and (iii) within 2010-2019. The number of annual disruption observations combined with the duration, can be used as a proxy for the disturbance intensity and impact level.

In the last mapping step, we created a collection of 30 maps providing the spatial extent of the TMF and disturbance classes on a yearly basis, from 1990 to 2019, using dedicated decision rules (see Supplementary Text on the annual change dataset and thematic maps). These maps were used in our annual trend analysis-described in next subsection-to document the annual disturbances over the full period, with ten classes of transition for each annual statistic (**Figs. 3** **and** **4**, figs. S1 and S11): (i) degradation that occurs before deforestation, (iii) short-duration degradation not followed by deforestation, (iv) long-duration degradation not followed by deforestation, (v) direct deforestation (without prior degradation) not followed by *forest regrowth*, (vi) direct deforestation followed by *forest regrowth*, (viii) deforestation after degradation followed by *forest regrowth*, (viii) deforestation after degradation not followed by regrowth, (ix) forest conversion to water bodies and (x) forest conversion to tree plantations. The associated metadata information on *invalid observations* within the forest domain and the proportion of *invalid observations* over the forest domain area were also documented.

In order to produce a more conservative map of undisturbed forests by excluding potential missed areas impacted by logging activities, we created a disturbance buffer zone using a threshold distance of 120 m around disturbed pixels. This distance corresponds to the average observed distance between two logging desks (landing) and is consistent with the distances used in previous studies for assessing intact forests (**15**).

### Trend analysis

The areas of TMF and disturbance classes are reported yearly and at 5-year intervals between 1990 and 2019, by country, subregion and continent (**Tables 1**, **2**, **and** **3**, **Figs. 3** **and** **4** and fig. S11, Supplementary Text on Trend analysis), using the country limits from the Global Administrative Unit Layers dataset from the FAO (**53**). Area measurements were also computed for 1° × 1° cells of a systematic latitude–longitude grid in order to delineate hotspot areas of deforestation and degradation for the three decades (**Fig. 5**). For the three most recent years of the considered period (i.e. for 2017-2019), the proportions of disturbance types (degradation followed by deforestation, degradation not followed by deforestation and direct deforestation) were calibrated with historical proportions (2005-2014) of the three types of disturbances.

For countries with moist forest areas larger than 5 million ha in 1990 (i.e. for 32 countries), and for all sub regions, we analyzed the temporal dynamics of annual changes from 1990 to 2019 (fig. S11 and Supplementary Text on trend analysis).

### Validation

The performance of our classifier was assessed in term of errors of omission and commission at the pixel scale and the uncertainties in the area estimates derived from the transition map were quantified (see Supplementary Text on the validation). A stratified systematic sampling scheme was used to create a reference dataset of 5 250 sample plots of 3 × 3 pixels (0.81 ha plot size) (fig. S8). For each sample plot, Landsat images at several dates were visually interpreted, together with the most recent HR images available from the Digital Globe or Bing collections, to create the reference dataset. The dates of the Landsat images to be interpreted were selected to optimize the assessment of the performance of our classifier as follows (fig. S9); (i) at least one random date within three successive key periods to verify the consistency of the temporal sequencing and the classifier performance across the main sensors (L5, L7 and L8), (ii) for the disturbed classes, the two dates corresponding to the first and last *disruption observations* were selected to assess the commission errors, and (iii) for the undisturbed forest class, at least one random date during the Global Forest Change (GFC) loss year (if existing) to assess omission errors. It resulted into the interpretation of two to four Landsat images for each sample plot, with a total of 14 295 images.

The user, producer and overall accuracies, the confidence intervals of the estimated accuracies and the corrected estimates of undisturbed and disturbed forest areas with a 95% confidence interval on this estimation were computed in accordance with latest statistical good practices (**26**). The performance of our disturbance detection results into 9.4% omissions, 8.1% false detections and 91.4% overall accuracy (tables S2 and S3). In addition, the uncertainties of area estimates (forest cover and changes) have been assessed from a sample of 5119 reference plots. This accuracy assessment shows that a direct area measurement from the forest cover maps underestimates the forest area changes by 11.8% (representing 38.4 million ha, with 15 million ha confidence interval at 95%) (tables S4 and S5).

### Comparison with the Global Forest Change (GFC) dataset

We compared our transition classes with the GFC dataset (**24**) for the TMF domain (undisturbed and degraded forest) in 2000 and over the period 2001-2019, which is the common period between the two products.

We synthesized the GFC multiannual product into four classes of forest cover changes from the combination of the GFC annual layers of tree cover loss and gain over the period 2001-2019: (i) unchanged (no loss, no gain), (ii) at least one loss but no gain, (iii) at least one gain but no loss and (iv) at least one loss and one gain. A new version of the transition map with eight classes was created (through the combination with annual maps) to characterize the disturbances that occurred between 2001 and 2019: (i) undisturbed forest (at the end of 2019), (ii) old degradation or regrowth (initiated before 2001), (iii) old deforestation (before 2001), (iv) degradation initiated between 2001 and 2019, (v) direct deforestation initiated between 2001 and 2019, (vi) deforestation that follows a degradation and initiated between 2001 and 2019, (vii) regrowth initiated from 2001 (viii) other land cover.

A matrix of correspondences between the synthesized GFC map (four classes) and our reclassified transition map (eight classes) was then produced for each continent and for the pan-tropical region, where area estimates are compared (table S1). This comparison shows that our annual change dataset depicts 138.9 million ha of forest disturbances along the periods 2001-2019 that are not depicted in the GFC map (representing 59% of the total area of our disturbances). This finding is corroborated by previous studies (**33, 31**). In addition, 17.6 million ha and 3.2 million ha are depicted as a GFC loss whereas it is classified as old deforestation and degradation respectively (before 2001) in our TMF dataset. Amongst the disturbances that are not depicted by GFC, the highest disagreements concern the gradual processes such the degradation, the *forest regrowth* classes, and the deforestation that follows a degradation for which 75%, 67% and 59% respectively of our depicted areas are missing on the GFC map, whereas our direct deforestation class shows a good correspondence with the GFC map (60%). The disagreement between our dataset and the GFC map is even higher for the changes within the mangroves with 83% difference. Mangroves are a key ecosystem within the TMF. We also observed a lower agreement for the disturbance classes in Africa (38% of our disturbances are depicted by GFC) compared to other continents (40.9% and 43.3% for Asia and Latin America respectively). A higher underestimation of GFC loss in Africa compared to other continents has also been observed by Tyukavina et al. (**31**) using a sample-based analysis.

We observe higher discrepancies between GFC and our study for shorter and lower intensity events, i.e. (i) the average duration for the disturbances detected only by our approach is 6.7 years compared to 9.4 years for the disturbances captured by both approaches, and (ii) the average intensity (or total number of disruptions detected for each disturbance) for the disturbances detected only by our approach is 9.9 compared to 32.6 for the disturbances captured by both approaches.

The evolution of the discrepancies over time shows major differences between the period (2001-2010) where our annual change dataset depicts 61.4% more deforested areas, and the last decade (2010-2019) where GFC losses include all our deforestation areas and 5.7% of our degradation areas (**Table 4** **and** **Fig. 4**). This change in the last decade has also been observed in another study (**56**) and can be explained (i) by the differences of processing applied by GFC team before and after the year 2011 (https://earthenginepartners.appspot.com/science-2013-global-forest/download_v1.3.html), and (ii) by the inclusion of burned areas in the GFC loss (particularly for the dry period of 2015-2016) that are mainly classified as degradation in our TMF dataset.

### Projection of future forest cover

Temporal projections of future forest cover are provided for (i) undisturbed forest area and (ii) total forest area (undisturbed and degraded forests) per country (fig. S12. and table S8). We considered that the annual disturbed areas followed an independent log-normal distribution for each country, and we used a modified version of the Cox method to estimate the mean and the 95% confidence interval (**58**) of the distribution. We used these estimates on the last 10 years (period 2010-2019) to project disturbances over the period 2020-2050 under a business-as-usual scenario. Several metrics, with their uncertainties, have been produced: (i) forest area at the end of 2050, (ii) percentage of remaining forest area at the end of 2050 compared with forest area at the end of 2019 and (iii) year corresponding to full disappearance of forest cover.

### Known limitations and future improvements

Disturbances that affect less than the full pixel area (0.09 ha size), e.g. the removal of a single tree, are generally not included in our results because the impact of the spectral values of the pixel are not strong enough to be detected. However, in specific cases, where the impact on the forest canopy cover modifies significantly the spectral values within a single pixel, e.g. the opening of a narrow logging road (< 10 m wide) or the removal of several big trees, our approach can detect such disturbances.

We have addressed the geographic and temporal discontinuities of the Landsat archive (see Data and Mapping method) by determining at the pixel level (i) an initial period (baseline) of minimum four years (increasing when the annual number of valid observations is low) for mapping the initial TMF extent and (ii) a monitoring period for detecting the changes. This minimizes the risk of inclusion of non-forest cover types (such as agriculture) and deciduous forests in the baseline when there are few valid observations over a short period. This risk has been under-estimated by previous studies that did not use a long period of analysis and did not accounted for the number of valid observations.

The accuracy of the disturbance detections has been assessed in the validation exercise (see Validation section and Supplementary Text on the validation). The assignment of the disturbance types at any location improves as the number of valid observations increases. The meta-information documents (i) the annual number of valid observations (ii) the first year of valid observation (fig. S15) and (iii) the start year of the monitoring period (fig. S18) at each pixel location. This meta-information (in particular the number of valid observations) can be considered as a proxy measure of confidence. Hence our estimates of changes in the regions where the total number of valid observations is particularly low and/or the start year of the monitoring period is late (figs. S14, S15, S18ra), e.g. Gabon, Salomon Islands, La Reunion, should be considered with lower confidence. However, considering the geographic completeness of Landsat-8 coverage after year 2013 there is high confidence for the contemporary reported estimates.

Short-duration events are likely to be underestimated for regions with geographic and temporal discontinuities in the Landsat archive and/or with gaps caused by persistent cloud cover. This is the case of Africa which is poorly covered by Landsat acquisitions before year 2000 (fig. S16). In order to provide a more conservative estimate of the remaining undisturbed forested areas, we also produced another estimate of undisturbed forested areas using a buffer zone with a threshold distance of 120 m from the detected disturbed pixels to exclude the potentially edge-affected forest areas. Further contextual spatial analysis would be needed to better estimate the characteristics of fragmented areas.

For the first time at pan tropical scale, a fine spatial resolution and annual frequency, detailed information on the historical forest area changes within the plantation concessions of oil palm and rubber are provided through to the combination of ancillary information and dedicated visual interpretation (see Supplementary Text on ancillary datasets). Although some confusion between forests and old plantations may remain (in particular for plantations that are not included in the ancillary database of concessions or that cannot be easily identified visually on satellite imagery from a regular geometrical shape), such errors are expected to be limited due to the consideration of (i) a minimum duration for the initial period and (ii) a long observation period. Classes of tree plantations do not include all commodities such as coffee, tea and coconut, that are detected as deforested land (if initially TMF and converted in commodity during the monitoring period) or other land cover (if the concession was already established during the initial period).

Some isolated commission errors may remain in the bamboo-dominated TMF, wetlands and semi-deciduous forests as reference data were available on restricted areas (Supplementary Text on specific tropical forest types). These will be continuously improved as the reference information layers improve and based on the feedback of users and national authorities.

The L7 SLC-off issue may introduce some spatial inconsistencies owing to a higher number of valid observations outside the SLC-off stripes which allows more disruptions to be captured and leads – potentially - to a different transition class.

Efforts have been done to classify disturbances based on their characteristics (timing, recurrence and sequence) in order to fit to the land cover use. However, all the metrics used in this study are made freely available to the end-user to possibly apply different decision rules that would better fit to the specific user needs and constraints, e.g. threshold applied to discriminate deforestation from degradation may be different according to the selected definition of the degradation.

This approach can be automatically applied to future Landsat data (from 2020) and is intended to be adapted to Sentinel 2 data (available since 2015) towards a monitoring of tropical moist forests with higher temporal frequency and finer spatial resolution.

## SUPPLEMENTARY MATERIALS

This file contains Supplementary Text on ancillary data, on specific tropical forest formations, on the transition map, on the annual change dataset, on the validation, on the trend analysis, supplementary references, supplementary figures and supplementary tables.

## ACKNOWLEDGMENTS

The USGS and NASA provided the Landsat imagery. R. Moore and her team provided the Google Earth Engine. Noel Gorelick, Mike Dixon, and Chris Herwig provided support in GEE. Andrew Cottam provided support for the validation tool (adaptation from the water surface validation tool). Andreas Langner, Hans-Jurgen Stibig, Rene Beuchle, Astrid Verhegghen, Hugh Eva, Rosana Grecci, and Baudouin Desclee provided a useful feedback on the legend and products in the framework of the ReCaREDD (Reinforcement of Capacities for REDD+) and REDDCopernicus projects. We thank Alessandro Cescatti and Philippe Mayaux for reviewing earlier versions of the manuscript.

This study was funded by the Directorate-General for Climate Action of the European Commission (DG-CLIMA) in the framework of the *Roadless-For* pilot project (Making efficient use of EU climate finance: Using roads as an early performance indicator for REDD+ projects).

## CONTRIBUTIONS

CV was the principal investigator of this study. CV developed the expert system, implemented all the steps and analyzed the results. The manuscript was prepared by CV and FA, with the contributions of J-F.P., GV, JG, LA and RN. FA contributed to the analysis of the results. CV, FA and JG developed together the validation method. J-F.P contributed to the development of the expert system. GV realized the deforestation predictions, provided support with Python and gave a useful feedback on the maps produced. SC realized the validation exercise and contributed to the creation of the plantation database. DS gave a support for coding with GEE and Python. AM and DS realized the website.

## Notes

### Competing Interest Statement

The authors have declared no competing interest.

