## Supplementary information for "Long-term (1990-2019) monitoring of tropical moist forests dynamics"

### - Supplementary Text –

### ***Supplementary Information on ancillary datasets***

Three ancillary datasets were used to spatially attribute disturbances (i) to the conversion to commodities or tree plantations (mainly oil palm and rubber), (ii) to the conversion to water bodies (conversion mainly due to new dams), and (iii) to specific changes within the mangroves.

#### *1. Conversion to commodities or tree plantations*

In the transition map, commodities (oil palm) or tree plantations (rubber) appear either as deforested land or other land cover class when established during the monitoring period or before the initial period respectively. They can also appear as undisturbed forest or forest regrowth when established a long time before the initial period with a spectral signature similar to a forest or a forest regrowth, e.g. in the case of old oil palm plantations.

In order to reduce these commission errors we created a mask of commodities or tree plantations from two external data sources: (i) the planted tree datasets from the World Resources Institute (WRI) (**59, 60**), which cover 14 countries (Brazil, Cambodia, Colombia, Indonesia, Liberia, Malaysia, Peru, Chili, Gabon, Ghana, Argentina, Honduras, Guatemala, Australia) and several plantation types, from which we only used 'Large industrial plantations' and 'Clearing/very young plantation not mosaic', and (ii) the oil palm dataset from Duke University (**61**), which covers a few plantation zones in the tropics.

Both datasets have been checked visually against high-resolution (HR) images available from Google Earth Engine (GEE) (**22**) and areas that are validated by the photo-interpreter as covered by commodities and with a correct delineation are incorporated in the commodities mask. Commodities that are well identified on the HR images but with a wrong delineation have been manually re-delineated by the interpreter and incorporated in the commodities mask. Then we carried out a further control of the transition map to identify and delineate missing areas of commodities that are not identified from existing databases. This step was done through a systematic analysis within the transition map for all areas with specific geometric shapes corresponding to plantations. These commodities were delineated visually by the photo-interpreter and incorporated in the commodities mask.

This new class of commodities allows also assessing the area of conversion of moist forests to such commodities. All pixels of the commodities mask are reassigned to the classes of conversion to commodities, e.g. a pixel that was initially labelled as deforested is reclassified as "conversion to commodities" with three sub-classes: establishment before 2000, in 2000-2009, in 2010-2019.

*2. Conversion to water*

To identify conversion from moist forests to water bodies, which are usually due to the creation of a dam, we used the Global Surface Water (GSW) dataset derived from Landsat time series over the period 1984-2015 (**41**) and the GSW updates for the period 2016-2019. This allowed to create two additional classes of deforestation: (i) forest conversion to a permanent water body and (ii) forest conversion to a seasonal water body. The GSW time series also provided information on the start date of forest conversion (flooding) when the forest was directly flooded without prior clearance. We have also integrated the inter-annual variations of the water bodies in our annual change product by discriminating permanent water from seasonal water from other land cover classes.

#### *3. Changes within the mangroves*

We created a specific class of mangroves to assess the status and changes of this specific forest ecosystem. We used the Global Mangrove Watch (GMW) dataset (**62**) to create an initial map of mangroves. As the GMW dataset covers the years 1996 and 2006, we first produced a maximum extent mask of the mangroves during the period 1996-2006 and then reassigned the transition classes under the maximum extent mask to produce a map of changes in mangroves. As example of reassignment, “undisturbed TMF (class 10 of the transition map)” is reclassified as “undisturbed mangrove (class 12)”. Eight classes of changes within the mangroves (including degradation, regrowth and deforestation) have been documented with sub-classes for each period of disturbance.

***Supplementary Information on specific tropical forest types***

- The tropical moist forest areas with bamboo dominance might be mis-classified as disturbed forests due seasonal or occasional defoliation of the bamboos. Therefore, we use a specific approach to map (as a specific class 11 of the transition map) the bamboo-dominated forests (*pacales*) in two specific zones where they are present over large areas: the Brazilian state of Acre and of east Peru. We first created a spatial mask for reclassifying false disturbances in these two zones. The spatial mask is created from the South-America regional part of the Global Land Cover 2000 (GLC2000) map (**63**) combined with a visual interpretation of recent high-resolution imagery. Dedicated decision rules are then applied for pixels classified as disturbed forest cover within this mask based on the recurrence, intensity and distance from roads and rivers of the disruption events to separate false disturbances from real disturbances (e.g. disturbances within 120 m from roads are kept).

- Our tropical moist forest domain includes semi-deciduous forests that appear as evergreen forests along a full year. However, specific types of forest transitions between the humid and dry ecosystem domains, e.g. the Chiquitania forests of northern Bolivia (**63,64**) can behave alternatively as evergreen or seasonal forests during specific years depending on the yearly amount and distribution of precipitations (**64**). If these forests behave as evergreen at the beginning of the Landsat archive, temporary defoliation due to drier conditions in the second part of the archive may be mis-classified as a disturbance, e.g. as a degradation when defoliation is observed along less than a duration of 900 days. We apply here also dedicated classification rules for avoiding such misclassifications due to the variable seasonal character of this specific forest type. The rules are based on the recurrence and intensity of the disruption events and are combined with a spatial mask that is created from the South-America regional component of the GLC2000 map (**63**) and visual interpretation of high-resolution imagery.

- Some savanna wetlands (wet meadows or marshes) can be misclassified as forest or disturbed forests due the scarcity of dryness periods. Such errors are expected to be limited due to (i) the consideration of a minimum duration for the initial period and (ii) the use of the GWSE dataset that documents the water seasonality. A region that is regularly flooded since the beginning of the observation period is assigned to the permanent or seasonal water class and hence cannot be classified as a TMF. However, for regions with geographic and temporal discontinuities in the Landsat archive occasionally or never detected as water, and/or with gaps caused by persistent cloud cover, periods of dryness may be not well covered with Landsat imagery during the initial reference period. To avoid these commission errors, wetlands areas have been identified using the Global Wetland atlas (https://www2.cifor.org/global-wetlands/) and the GLC 2000 map (**63**), with a further visually check on HR imagery. When misclassification was detected, these areas have been visually delineated and re-assigned to the 'other land cover' class (# 90 in the transition map).

***Supplementary Information on the transition map***

The transition map shows the spatial distribution of the moist forest at 30 m resolution for year 2019 and allows the identification of small logging impacts such as skid trails and logging decks in concessions (fig. S3A-D) and small linear features such as gallery forests (fig. S4C).

Various types of deforestation and degradation events are mapped. In particular, impacts of logging activities are captured with different intensity levels, from selective logging impacts, which are mapped as degraded forests with short duration of disturbance detection (fig. S3A-D) to conversion of forest cover to another land cover (mainly pasture or crops) (fig. S4A and B) or vegetation regrowth.

Conversion to tree or shrub plantations occurred mainly for oil palm and rubber tree in Africa and Asia (fig. S5B-D).

Small-scale agriculture also contributes significantly to the conversion and degradation of forests. This is the case in both Madagascar and the Democratic Republic of the Congo (DRC), where shifting cultivation, small tree plantations, irrigated crops, forest regrowth and dense humid forests are often present together as a 'rural complex' in landscapes around villages (fig. S6C) or cover major parts of the landscape (Madagascar).

In Ivory Coast, most undisturbed forests which were remaining in 2008 have disappeared, except in a few remaining protected areas (fig. S4D).

Mining exploitation of metals and precious minerals is also a cause of deforestation, such as gold mining within the dense forest, often along river courses (fig. S6D). The infrastructure used for petrol extraction (drilling, pipelines) in Gabon also causes damage to forests.

Strong El Niño southern oscillation (ENSO) events cause droughts and subsequently fires, which can lead to long term degradation or be followed by full death of tree cover, such as in Cambodia, where a semi-evergreen forest dried up in 2016 (fig. S7B). The ENSO event that occurred in 2015-2016 caused extreme drought in the northern Brazilian Amazon and induced the burning of forest cover (fig. S7D).

Other extreme natural events with very short durations can also cause forest damage, for example Debbie cyclone in Australia in April 2017 (fig. S7A), Maria hurricane in Puerto Rico at the end of 2017 or Hudah cyclone in Madagascar in April 2000 (fig. S7C).

Transport infrastructure such as main roads and railways is well captured on the transition map within the dense humid forest (fig. S4C). The impacts of new dams are identified as conversion from undisturbed forest to seasonal or permanent water (fig. S6A). Finally, the significant differences in forest cover patterns between bordering countries illustrate differences in resource management policies (fig. S6B).

***Supplementary Information on the annual change dataset***

The annual change dataset is a collection of 30 maps depicting - for each year between 1990 and 2019 - the spatial extents of undisturbed forests and disturbances. The annual maps depict the thirteen following classes: (i) moist forest, (ii) moist forest before the establishment of a tree plantation, (iii) bamboo dominated moist forest, (iv) new degradation (disruptions detected for the first time during the considered year), (v) ongoing degradation (disruptions started before the considered year and are still detected), (vi) degraded forest (disruptions started before the considered year and are not detected anymore), (vii) new deforestation (disruptions detected for the first time during the considered year), (viii) ongoing deforestation (disruptions started before the considered year and are still detected), (ix) new regrowth (deforestation occurred the year before and disruptions are not detected anymore), (x) regrowthing (deforestation occurred at least one year before and disruptions are not detected anymore), (xi) water (permanent or seasonal), (xii) other land cover, and (xiii) invalid observations.

To obtain the annual classes, we combined the transition map with the following spatial layers: (i) number of disruption observations per year, (ii) first and last year of a disturbance period (YearMin and YearMax), (iii) recurrence of disruption observations (iv) Start Year of the archive (first year after the initial period), and (v) number of valid observations per year. The creation of annual maps is made from the following rules (where $\text{Year}_i$ stands for year 1990 to 2019):

- Deforestation that occurs after degradation is separated from direct deforestation using the recurrence value. The starting years of the disturbances (two in the case of a degradation before deforestation) are recorded (YearMin and YearMin2, respectively).
- A disturbance is classified as new disturbance in YearMin (for degradation or direct deforestation) or YearMin2 (for deforestation after degradation).

- A disturbance is classified as ongoing disturbance after YearMin or YearMin2 until YearMax.
- In the case of a degradation disturbance, a pixel is classified as degraded forest after YearMax.
- A disturbed pixel of the TMF domain is characterized into one of the three following timing periods: (i) when $Year_i$ is before StartYear, (ii) when $Year_i$ is after the StartYear and the pixel is within tree plantation areas, (ii) when $Year_i$ is after the StartYear and the pixel is outside tree plantation areas.
- In the case of a deforestation disturbance, a pixel is classified as new regrowth on YearMax + 1, and then as ongoing regrowth from YearMax + 2.
- A pixel is classified as permanent water or as seasonal water if located within the permanent water area or within the seasonal water area in the GWS annual dataset, respectively.
- A pixel is classified as other land cover for $Year_i$ if located outside the moist forest domain and with at least one valid observation available during $Year_i$.
- A pixel is classified as no data for $Year_i$ if no valid observations are available for $Year_i$.

In order to discriminate deforestation without prior degradation from deforestation occurring after degradation, we applied two conditions to consider that deforestation occurred after degradation: (i) a recurrence value lower than 58%, or (ii) a recurrence value lower than 70% with at least 6 years without any disruption events between the degradation and the deforestation disturbances. These conditions were determined empirically by analyzing various sequences of logged and deforested areas.

In the case of a degradation not followed by a deforestation, two temporal sequences can be potentially observed: (a) only one degradation disturbance is observed with less than 3 years duration and no other disruption events are detected until the end of the observation period, or (b) two degradation disturbances are observed and are separated by a break of a minimum 4 years period without disruption events.

Figure S1 illustrates the changes visible from this dataset looking at two different years for two regions where important deforestation and degradation occurred during the observation period: South of Cambodia and the Para region in Brazil.

***Supplementary Information on Trend analysis***

*Results at pan-tropical, continental and subregional levels*

Our analysis shows that Tropical Moist Forests (TMF) covered 1059.6 million ha in January 2020 including 964.4 million ha of undisturbed forest. Degraded TMF cover 106.5 million ha and are distributed as follows: 36.9% in Latin America, 40.7% in Asia-Oceania and 22.3% in Africa. The remaining TMF represent 60% of the extent of tropical natural forests reported by the FAO for the year 2015 (**30**).

At pan-tropical level, during the period 1990-2019, 189.2 M ha, 29.5 M ha and 106.5 M ha of undisturbed tropical moist forests were converted into non-forest cover, forest regrowth and degraded forests, representing 58.2%, 9.1% and 32.8% of the overall disturbances, respectively (**Table 3**). Hence the overall loss of TMF (i.e. forests converted to another vegetation type or to young forest regrowth) is of 218.7 million ha, representing 17.3% of the initial extent of TMF, or the extent of Saudi Arabia. In addition, the degraded forest area at the end of 2019 (106.5 million ha) represent 8.4% of the initial extent of undisturbed TMF. In Asia the sum of degraded forests and forest regrowths represents 48.5% of forest changes, whereas it represents only 41.4% and 36.6% of forest changes in Africa and Latin America respectively. The percentage of disturbance-edge-affected forests (of the initial extent of undisturbed forest) is also higher in Asia than in other continents, i.e. 43% vs c. 26%.

The extent of undisturbed tropical moist forests has declined by 23.9% since 1990 with a peak rate during the period 1995-1999 at 14.4 million ha/year (**Table 2**). This late 1990s peak rate happened in most regions with enough valid observations during the period 1990-1999, i.e. all regions except Central and West Africa where the peak happened in 2000-2004.

Among the three continents, Asia shows the largest relative decrease in undisturbed tropical moist forest cover since 1990, with 37.9% relative decrease compared to 24.4% and 19.9% decrease for Africa and Latin America respectively (**Table 1**). Continental Southeast Asia lost 53.3% of its undisturbed moist forest area, either through direct deforestation (33.3% of total disturbances) or deforestation after degradation (35.4%) or through degradation (31.3%) (**Table 2**). For this region, the decline of undisturbed forest was slightly faster before 2000 (1.6 and 1.3 million ha/year before and after 2000, respectively) (**Table 2**). However, the annual rate of loss of undisturbed forest cover slightly increased in the last five years (to reach 1.14 million ha/year) mainly due to degradation.

The Central and West Africa regions also show a faster decline during the last five years compared to the period 2005-2014 (c. 32% and 48% of increase respectively for Central and West Africa regions), with an overall loss of their undisturbed forests in 2000 (which covered 216 and 32.8 million ha, respectively) of 14.5% and 52.6% respectively.

Latin America shows the largest rate of loss of undisturbed forest among the three continents, in particular between 1990 and 2000 (ranging between 6.2 and 6.3 million ha/year), with a strong reduction after 2005 (from 6.2 million ha/year in 2000-2004 to 4 million ha/year in 2005-2019) mainly due to the decrease of the direct deforestation rate (from 3.1 million ha/year to 1.7 million ha/year). The degradation rate decreased from 3.1 million ha/year in 2000-2004 to 2.3 million ha/year in 2005-2019.

Central America lost 53% of its undisturbed forest area in 1990 (34.5 million ha).

South America experienced a peak in deforestation during the period 1995-1999 and then a progressive reduction from 4.4 million ha/year in 1995-1999 to 1.9 million ha/year in 2010-2015. The degradation rate slightly decreased from 2.6 million ha/year in 1995-2000 to 2.1 million ha/year in 2010-2015.

Asia shows the largest area of forest conversion to commodities (86% of all pan-tropical plantations), which represents 15.2% of continental disturbances, compared with 1.7% and 0.8% for Latin America and Africa, respectively.

The proportions of disturbance types present at the end of 2019 are the following: 3.6% of disturbances were long-duration degradation, 20.7% were short-duration degradation (one stage), 10.4% were short-duration in 2/3 stages, 53.8.3% were deforested, and 8.5% were forest regrowth after deforestation.

When we considered a buffer zone around pixels detected as disturbed forests (**15**), our estimate of edge-affected degraded forest area increased by a factor of 3.3 or 5.1 for threshold distances of 120 m or 240 m, respectively.

*Area estimates at national level*

The three Southeast Asia countries with the largest remaining areas of undisturbed moist forest are Indonesia, Papua New Guinea and Malaysia, with losses representing 42.1%, 17.7% and 49.1% of their forest area in 1990. Indonesia was ranked as the second largest country (after Brazil) for area of undisturbed tropical moist forest in the early 1990s but is now (in 2019) in third position behind Brazil and the DRC.

All countries in continental Southeast Asia have already lost more than half of their undisturbed moist forest (up to 68% for Vietnam, 65% for Lao PDR, and 61% for Cambodia and Myanmar) (Supplementary Table 2). In Cambodia, Laos, Myanmar and Vietnam, 42%, 29%, 26%, and 23%, respectively, of overall disturbances occurred during the last 10 years. Papua New Guinea is the Asian country with the lowest loss rate (0.24 million ha/year on average over the last 10 years) and is now ranked as the seventh country in the world for undisturbed TMF.

The majority of forest conversion to oil palm and rubber occurred in Asia (18.3 million ha from an overall conversion of 21.3 million ha), with the largest contribution from Indonesia and Malaysia (12.7 and 5.3 million ha, respectively representing 19% and 38% of their overall disturbances).

A few Asian countries were also impacted by the creation of new dams within the moist forest (1.4 million ha of forest conversion across the continent).

All Southeast Asian countries showed a decline in their disturbance rates after 2000 and most of them were particularly affected by the strong El Niño events of 1997-1998 and 2015-2016. The high peak in disturbances in 1997 in Indonesia was mostly caused by forest fires in Sumatra, which led to 3.5 million ha of new degradation (burned forest that regrown) and 1.7 million ha of direct deforestation (burned forest that were converted to other land cover). Papua New Guinea, Thailand, and Cambodia were the Asian countries most affected by the ENSO event of 2015-2016.

Five Latin American countries are among the 10 countries with the largest areas of undisturbed tropical moist forest: Brazil, Peru, Colombia, Venezuela and Bolivia, representing together more than half of the remaining moist tropical forests (496 million ha).

Disturbance rates decreased significantly in (2000-2014) compared to (1990-1999) mostly in Brazil and Mexico (reduction of 0.62 and 0.16 million ha/year, respectively), and increased in Colombia, Venezuela, and, Nicaragua and Ecuador (0.23, 0.17 and 0.09 million ha/year, respectively).

Between 1990 and 2019, the area of undisturbed tropical moist forest in the Legal Amazon declined by 17%, from 353 million ha in 1990 to 292 million ha in 2019. The annual disturbance rate decreased significantly after 2005 (**Fig. 4**), from 3.3 and 2.7 million ha/year in 1999 and 2003, respectively, down to 0.9 million ha/year in 2012. By 2015, Brazil had accomplished a 60% reduction in the deforestation rate of its Amazonian forests from the peak in 1995-2000. However, the annual deforestation rates of moist forests in the Brazilian Amazon have dramatically increased since 2015, to reach 3.9 million ha/year in 2016.

Our estimate of direct deforestation in Brazil is similar to Brazilian National Institute for Space Research (INPE) estimate for the humid domain of the Amazônia Legal, particularly from 2004 when INPE implemented a digital processing approach to mapping deforestation (**29**) (**Fig. 4A**).

Amongst the American countries, Mexico lost the highest proportion of undisturbed forest (a 74% reduction), followed by Nicaragua (66%).

In Brazil a total of 0.6 million ha of undisturbed forests have been converted into water bodies, including the creation of the Balbina, San Antônio and Belo Monte dams. Peru, Bolivia, and Venezuela lost 0.14, 0.11 and 0.10 million ha of undisturbed forests, respectively, through the conversion to water bodies.

In Latin America 2.5 million ha of moist forests were converted to tree plantations, the major part being located in Brazil (1.62 million ha) and minor contributions from Venezuela and Peru (0.07 and 0.04 million ha, respectively).

Peaks in disturbances are observed during strong El Niño events that led to severe droughts in South America, in particular in 1991-1992, 1997-1998, 2009-2010 and 2015-2016 (**Figs. 3 and 4**, and fig. S11). The warm and dry weather that occurs during El Niño years provides optimal conditions to cause and spread fires in the Amazonian forests (**28-29**). This is made worse by the fact that fires are used as a tool to clear areas of tropical moist forest for agriculture and can spread out of control.

DRC is the African country with the largest remaining extent of undisturbed moist forest at 105.8 million ha, being the second largest country area at tropical level. Gabon, Cameroon and the Republic of Congo have similar areas of remaining intact forests (between 19.8 and 23.4 million ha in 2019). The Republic of Congo and Gabon show very low rates of decline for the period 2000-2019 (0.03-0.1 million ha/year) compared to the DRC (1.4 million ha/year).

All African countries show an increase in annual rates of disturbances after 2009, except Madagascar and Angola.

The African countries with the largest reductions in undisturbed moist forest extent are Ivory Coast (81.5%), Ghana (70.8%), Madagascar and Angola (67%), Nigeria (47%) and Liberia (36%).

In West Africa, the disturbance rate shows a recent increase, with 1.1 million ha/year during the last five years (2015-2019) compared with 0.68 million ha/year for the period (2005-2014).

The major part of forest areas converted to tree plantations in Africa are located in DRC (0.08 million ha), Cameroon (0.07 million ha), and Gabon (0.04 million ha each).

### ***Supplementary Information on Validation***

#### ***Validation Method***

The validation approach includes three steps to produce an accuracy assessment: (i) the sampling design, (ii) the response design and (iii) the production of confusion matrices and estimates of uncertainties. The sampling design consisted of defining the spatial distribution of the sample within our study zone. The response design consisted of defining the protocol of measurements over the sample plots, including the selection of the dates of Landsat images to be interpreted.

*Sampling design*

The most frequent sampling approach for validating land cover maps is a stratified random sampling with strata defined from the classes of the map to be validated and with an independent random sample in each stratum (**65-67**). However, in our case, the transition map depicted temporal land cover changes that made this solution difficult to apply. Here we selected a stratified systematic sampling scheme, which provides unbiased estimators of accuracy, although it lead to a non-unbiased and more complex estimation of the variances. The main considerations for this choice are:

- Under spatial correlation decreasing with the distance, systematic sampling is more accurate than random sampling, i.e. the actual sampling variance is lower. However there is no unbiased estimator for the variance of systematic sampling and the usual random sampling estimator overestimates the systematic sampling variance, leading to a conservative accuracy assessment (**68,69**).
- If the sample size or the stratification are modified after the plot data collection start (e.g. because of changes in resources or improvement of the stratification), traditional systematic sampling with independent sampling may lead to a completely new sample (**69**). Our selected

stratified systematic sampling minimized this drawback by using a common pattern of ranked replicates for all strata.

- Bi-dimensional systematic sampling usually relies on a regular grid, which should be applied in principle on an equal-area projection. Although geographical coordinates do not correspond to an exact equal-area projection, the area distortion of geographical coordinates has a limited impact in tropical regions.

The sample was designed using three steps: (i) the preparation of a two-level systematic grid of potential sample points, (ii) the creation of a stratification layer by combining the transition map with an ancillary layer and (iii) the selection of a set of second-level replicates to reach the target number of sample plots per stratum and continent.

We first defined a grid of regular blocks of 1° × 1° latitude–longitude size that covered our study zone. A random location was selected within one block, then the set of points that occupied the same position in each block defined replicate 1 (fig. S10). The location of the second replicate was selected randomly within the 1° × 1° block among the locations that maximized the distance from replicate 1. The distance $d(1,2)$ between replicate 1 and 2 was the minimum distance between two points from each replicate that could belong to adjacent sampling blocks. For squared sampling blocks, there was only one location that reached the maximum distance for replicate 2. Replicates 1 and 2 constituted together a new systematic pattern following diagonal lines. Additional replicates could be added to intensify the sampling. To preserve a spatial distribution that was as homogeneous as possible, the location of each additional replicate was selected at random among those that maximized the distance to the previously selected replicates. Under the assumption that spatial correlation is higher at short distances, by maximising the distance between replicates, we reduced the redundancy of the information provided by the sample (**69**). The use of 1° × 1° blocks implies that the block size diminishes when moving away from the equator. Although this effect is limited within the tropics, we handled it by reducing the number of plots along each geographical parallel through fraction downgrading between replicates[66]. The parallel at latitude $\alpha$ has a relative length of approximately $\cos(\alpha)$ compared with the equator. Therefore, a portion of $[1 - \cos(\alpha)]$ plots belonging to replicate 1 was downgraded to replicate 2, a portion of $[2 \times (1 - \cos(\alpha))]$ was downgraded from replicate 2 to 3, and so on.

In the second step, we used the main classes of the transition map as core layers for the stratification, i.e. undisturbed, degraded forest, forest regrowth, deforested land, recent disturbances and other land cover. Moreover, in order to better assess potential omission errors in the mapping of disturbances, we added a supplementary (sub)stratum within the *undisturbed forest* stratum using the GFC dataset (**24**) as an ancillary spatial layer. To compensate for the shorter time coverage of the GFC dataset (compared with our dataset) we enlarged the GFC loss areas using a spatial buffer of 5 km, similarly to (**31**). This was intended as a proxy for GFC past deforestation (i.e. before 2000), as new deforestation often occurs close to places where deforestation has occurred in the past. This led to the division of the undisturbed forest stratum into two strata: stratum 1 (undisturbed forest outside the GFC loss buffer) and stratum 2 (undisturbed forest within the GFC loss buffer). Overall, this resulted in a total of seven strata (fig. S9).

Regarding the availability of information on the structure of the variance of the target variables across the strata, there is a variety of criteria that can be used to optimize the sampling allocation, for example the traditional Neyman's rule (**70**) or multivariate algorithms (**72**). As we were missing such knowledge on the variability of target variables per stratum, we allocated the same sample size (250 sample plots) to each stratum and each continent, leading to heterogeneous numbers of plots per replicate, stratum and continent. For example, for the large stratum 7 in Africa, all 197 sample plots from replicate 1 were allocated and 53 sample plots in replicate 2 were selected randomly to meet the target of 250 plots. For a smaller-sized stratum, higher-ranked replicates needed to be considered in order to find 250 plots (e.g. up to replicate 8 for stratum 2 in Africa). In spite of this sampling heterogeneity, the sampling algorithm ensured spatial regularity and avoided pairs of sample plots that were very close to each other. The overall sample consisted of 1 750 sample plots by continent (7 strata × 250 plots), i.e. 5 250 sample plots for the overall study area (fig. S8).

*Response design*

The reference dataset of land cover classes was created through visual expert interpretation of Landsat images at several dates and of recent higher resolution satellite images when available. Each reference sample plot was assessed over a box size of 3 × 3 Landsat pixels centered on one of the 5 250 sample points. For each sample plot, a sub-set of Landsat images was selected for visual interpretation. For each image and sample plot (0.81 ha size), the interpreter selected one of the following land cover labels: (i) *forest cover*, (ii) *mostly non-forest*, (iii) *minor non-forest*, or (iv) *invalid*. A *forest cover* label corresponds to mature trees or vegetation regrowth (mosaic of shrubs and trees), covering the full plot (9 pixels). The *mostly non-forest* label corresponds to a sample plot including at least five pixels with a *non-forest cover* (including disruption observations and other non-evergreen forest cover such as savanna, agriculture, and water surface). The interpreter assigned a *minor non-forest* label to the sample plot when one to four Landsat pixels with *non-forest* spectral signature were observed.

As it was not possible to interpret all the Landsat images available for each sample plot, we selected a subset of Landsat image dates for the visual interpretation, with the aim of optimizing the assessment of commission and omission errors and the resulting uncertainties (fig. S9), as follows:

- The Landsat images were selected from the full archive with at least one image within each of three key periods: (i) very recent years (2014-2017) corresponding to the acquisition period of Landsat 8 data, (ii) recent years (2007-2013) and (iii) historical period (before 2007).
- To assess the commission errors, we validated the detection of *disruption observations* from the same Landsat image that led to its detection. For each sample plot belonging to a disturbed stratum (i.e. strata 3-6) or to the *other land cover* stratum, the Landsat images corresponding to the dates of first and last *disruption observations* (or the dates of the non-evergreen forest cover observations for strata 7) were selected for visual interpretation.
- To assess omission errors (i.e. potential missed *disruption observations*), we validated the periods without *disruption observations* (green boxes in fig. S9) as follows. For stratum 1 (undisturbed forest with no GFC loss), three dates from the series of existing Landsat images were selected randomly, one for each of the three periods. For stratum 2 (undisturbed forest within the GFC loss buffer), three dates from the series of existing Landsat images were selected for visual interpretation: one date selected during the GFC loss year when the sample plot was covered by GFC loss pixels and two dates were selected randomly from Landsat images available during the two remaining periods. For strata 3-6, one date was selected randomly from available Landsat images during each forest/regrowth period (periods without *disruption observations*). In these cases (strata 3-6), the year just after or just before a period with *disruption observations* was

preferentially selected in order to validate the duration of the disturbance period; for example, for stratum 3 an image from 2007 was selected instead of a random selection from the period 2007-2013 (as the last disruption was observed before 2007).

This process of selection of Landsat images led to the selection of two to four images per sample plot and resulted in a total of 14 295 Landsat images to be visually interpreted.

*Interpretation interface/tool*

To interpret satellite images over the sample plots, a GEE web interface was developed to facilitate the photo-interpretation task by displaying (i) Landsat images at specific dates (see the subsection 'Response design' above), (ii) high-resolution (HR) images from the Digital Globe or Bing collections, and (iii) the sample box for each image.

For each sample plot and for each Landsat image, the expert validator had to select one class from the four land cover classes defined in the response design phase (*forest cover*, mostly *non-forest cover*, *minor non-forest cover*, or *invalid*). The expert validator did not have access to the results of our mapping approach (transition map or single-date classification) to avoid potential bias during this interpretation phase.

When an HR image was available, a more detailed land cover legend was used with the following classes: (i) *dense forest* (continuous tree cover with > 90% crown cover); (ii) *open forest* (non-continuous tree cover with > 50% crown cover); (iii) mostly *shrubland*; (iv) *forest/shrubland mosaic* (at least 10% of shrubs); (v) *minor non-forest* (10-50% non-forest cover); (vi) *mostly non-forest* (at least 50% non-forest cover); or (vii) *invalid* (no HR image available or clouds). Unfortunately, the exact dates of HR images are not usually available from GEE. Therefore, the HR interpretations were used in combination with the Landsat interpretations: (i) to support evidence in the validation process of the single-date classification algorithm, and (ii) to assess the accuracy of the transition classes and uncertainties of area estimates.

*Accuracy assessment of the single-date classification algorithm*

Our reference sample dataset was first used to assess the performance of the single-date classification algorithm in terms of errors of omission and commission. The accuracy was measured against the Landsat interpretations of the reference sample. The HR interpretations are provided in the detailed confusion matrix as complementary information to enable a better understanding of the commission and omission errors.

The HR interpretations were reclassified into five larger classes to make them comparable with the legend of the Landsat and single-date interpretations: (i) *forest,* (ii) *mostly non-forest*, (iii) *minor non-forest*, (iv) *shrub*, and (v) *invalid*. The *minor non-forest* and the *open forest* labels were grouped in one class (iii). The *mostly shrubland* and the *forest/shrubland mosaic* were grouped in one class (iv).

Using the full reference sample, a confusion matrix between the Landsat-based visual interpretations and the class values of our transition map was produced (for the three classes *forest cover*, *mostly non-forest*, and *minor non-forest)* from which a simplified 2-classes confusion matrix was derived with the forest and non-forest (including *mostly non-forest* and *minor non-forest*) classes. This was used to estimate overall accuracy and related omission and commission errors.

As the single-date classification was done using different sensors (TM, ETM+ and OLI sensors) onboard different satellites, we verified the consistency of classifier performance across the main sensors (L5, L7 and L8) by estimating the omission and commission errors for each sensor. Finally, the validation results were analysed by continent and for the different land cover strata, as well as for different disturbance intensities.

*Accuracy assessment of the transition map and uncertainties of area estimates*

Our reference dataset of sample plots was then used for the accuracy assessment of the transition map and for estimating errors in area estimates. For this accuracy assessment exercise, we considered the land cover classes of the transition map to produce four new classes at the scale of the sample plots (3 × 3 pixels) in order to make them comparable with the reference dataset: (i) *fully undisturbed forest* (all nine pixels of the transition map within the sample box were classified as undisturbed forest); (ii) *mostly undisturbed* (fewer than five pixels have changed), and (iii) *mostly changed*. The *mostly changed* class corresponds to sample plots with (i) at least five pixels that have changed from forest to non-forest or degraded forest, and (ii) fewer than five pixels that have been classified as other land cover.

From our reference dataset, we used the Landsat interpretations at different dates (from two to four dates) (fig. S9) combined with the HR interpretation to obtain the following potential classes for each reference sample plot:

(i) *undisturbed forest* (no interpretation of non-forest or mosaic forest/non-forest events) both on Landsat and HR (shrubland or mosaic forest/shrubland or invalid);

(ii) forest with *major or minor disturbance only on HR* (undisturbed forest on Landsat);

(iii) forest with *major disturbance*, i.e. with at least one interpretation of major disturbances (at least five pixels) on Landsat, whatever the HR interpretation;

(iv) forest with *minor disturbance*, i.e. with at least one interpretation of minor disturbances (fewer than five pixels) on Landsat, whatever the HR interpretation.

In addition, for the sample plots with disturbances identified either from the transition map or from the reference dataset of Landsat interpretations, we identified subclasses based on the number of Landsat images interpreted as disturbed. From the transition map, we defined three subclasses based on the number of disruption observations within the box and over the full 36-year period: (i) one disruption observation, (ii) between two and three disruption observations, and (iii) more than three disruption observations. For the reference dataset of Landsat interpretations, we identified four subclasses corresponding to the number of images that were interpreted as disrupted (including major and minor disturbances), i.e. 1, 2, 3 or 4.

The numbers of disruption observations in our sample were used to analyse the omission and commission errors between the transition map and the reference dataset.

To estimate the accuracy of the transition map, we used a simplified legend that allowed a good correspondence between the classes of the transition map and of the interpretations of the reference dataset. The simplified land cover legend included two target classes: (i) *undisturbed forests* and (ii) *forest changes*. From the transition map, a sample plot was considered *undisturbed forest* when the plot box was fully undisturbed (all nine pixels of the box) and was considered *forest changes* in the other cases, i.e. when the plot box contained at least one pixel of disturbance (i.e. including minor and major disturbances). From the reference dataset, a sample plot was considered *undisturbed* when there were no disturbance interpretations either on Landsat or on HR (or an invalid interpretation on HR) and *forest changes* in the other cases, i.e. when there was at least one disturbance interpretation either on Landsat or on HR.

The contributions of the sample plots were then weighted on the basis of the stratification used in the sampling phase (see the subsection 'Sampling design' above). Finally, the user, producer and overall accuracies, the omission and commission errors, the confidence intervals of the estimated accuracies and the corrected estimates of undisturbed and disturbed forest areas with a 95% confidence interval on this estimation were computed in accordance with the good practices recommended by Olofsson *et al*. (**26**).

#### ***Validation Results***

The validation was performed using a reference dataset of 5 119 sample plots with at least one valid Landsat interpretation per plot (1 705 for Africa, 1 693 for Asia and 1 700 for Latin America) and a total of 12 343 Landsat interpretations (3 823 for Africa, 4 215 for Asia and 4 305 for Latin America) distributed temporally (across the period 1982-2016) and across Landsat sensors (L5, L7, and L8). The 12 343 Landsat interpretations were compiled from the 5 119 sample plots using Landsat images at selected dates with no cloud presence, no sensor artefacts and no doubt about the visual interpretation (from one to four Landsat images per plot). Within the full sample of the 5 119 reference plots, 3 982 sample plots had one valid recent HR interpretation, 3% of the plots had one valid Landsat interpretation, 30% had two valid Landsat interpretations, 57% had three valid interpretations and 10% had four valid Landsat interpretations.

Two confusion matrices were produced from the reference sample dataset: one from all valid Landsat interpretations (12 343 in total), to assess the performance of the single-date classification algorithm, and another from the 5 119 sample plots, to assess the accuracy of the transition map.

##### *Performance of the single-date algorithm*

The confusion matrix for the single-date classification (tables S2 and S3) reports an overall accuracy of 91.4%, with omission and commission errors for *non-forest cover* detection of 9.4%

and 7.9%, respectively. At continental level, overall accuracy is higher for Africa (94.6%) than for Latin America (91.0%) or Asia (89.0%). Among the 577 plots that correspond to the omission errors, 86% were classified as forest changes or other land cover on the transition map. Moreover, 71% of these plots were confirmed as changes by the HR interpretations. This shows that most of the omissions of the single-date classification algorithm were 'temporary' omissions, as most of these disturbances were then confirmed from the full temporal Landsat time series. These probably correspond to omissions at the beginning of disturbance events. It is also important to mention that 87% of these omissions were plots with *minor non-forest* extent (fewer than five pixels interpreted as non-forest within the sample plot).

The accuracy matrices by continent (table S3) show a higher rate of omission errors for Asia (14.3%) than for Africa or Latin America (6.5% and 7.1%, respectively) but no significant differences were observed among sensors (9.8% for LT5, 9.7% for L8, 9.4% for L7).

These omission errors (in the single-date classification) mainly appear as other land cover class on the transition map (49% of these omissions), but also as mostly undisturbed (a spatial majority of undisturbed forest within the box) (25%), deforestation without regrowth (17%) and mostly degraded or regrowth (10% altogether). The 577 sample plots presenting omission errors are sites of intensive disturbances, as 71% of these errors concern sample plots where the total number of disturbances detected over the 37 years was greater than three.

Among the commission errors in the single-date classification corresponding to *forest cover* in Landsat interpretations and *non-forest cover* in the single-date classification (480 plots), only 22% were interpreted as *forest* using the HR images, whereas 24% were interpreted as mostly or *minor non-forest*, 35% as *shrubland* and 18% as *invalid*. Therefore, a large part of these 'false detections' of single-date disturbance events were observed as disturbances in the most recent HR images. This raises the issue of the potential limitations of the visual interpretation of Landsat images. Among the sample plots that correspond to these commission errors, 69% were classified in the transition map as spatial minor changes (fewer than five pixels within the 3 × 3 pixel box) and 31% as spatial major changes (more than five pixels within the 3 × 3 pixel box). In addition, of the total commission error plots, 67% concern deforestation and other land cover classes; degradation and regrowth represent 18%; and classes of mostly undisturbed forest (between five and eight pixels of undisturbed forest within the 3 × 3 pixel box) represent 15%.

More commission errors are observed for Latin America (13.2%) than for Asia (8.3%) or Africa (3.2%). These differences can be partly explained by the numbers of Landsat scenes that were processed: 540 634 scenes for Latin America, 482 965 scenes for Asia and 231 087 scenes for Africa. A larger number of scenes may result in a greater number of errors in the final product as a result of the presence of noise or artefacts within a minor part of the Landsat dataset, which cannot fully be eliminated.

Finally, more commission errors were observed for L8 (11.3%) than for L7 (8.2%) or L5 (7.3%) (table S3).

*Accuracy of the transition map and area estimates*

The accuracy matrix for the transition map shows an overall accuracy (stratum-weighted estimate) of 92.8% for the classes of the moist forest domain (tables S4 and S5). The omission and commission errors for the *forest changes* areas are 19% and 8.4%, respectively.

The commission errors concern mainly (66.3%) minor disturbances on our transition map (fewer than four disturbances detected over the observation period and fewer than five pixels within the sample plot). Of these commission errors, 24.2% concern major disturbances (more than five pixels within the plot with at least three detections over the full period).

The omission errors concern mainly (74.1%) minor disturbances that were identified only once from the valid Landsat images or disturbances that were identified only using the HR images (15.9%).

The accuracy matrix for the transition map allows us to produce reference-corrected area estimates (table S5). The correction shows that a direct area measurement from the transition map underestimates the forest area changes by 38.4 million ha (325.2 million ha derived from the map versus 363.6 million ha for the corrected estimate), representing a relative bias of 11.8%, with a confidence interval (95%) of this error estimation at 15 million ha.

**Supplementary Figures and Tables**

**Fig. S1** Subsets (25 X 34km) of the Annual Change layer at two different periods (1990 and 2015) for two regions: A, Cambodia (106 ˚ E, 12.5 ˚ N); B, Brazil - Para region (53˚ W, 6˚ S).

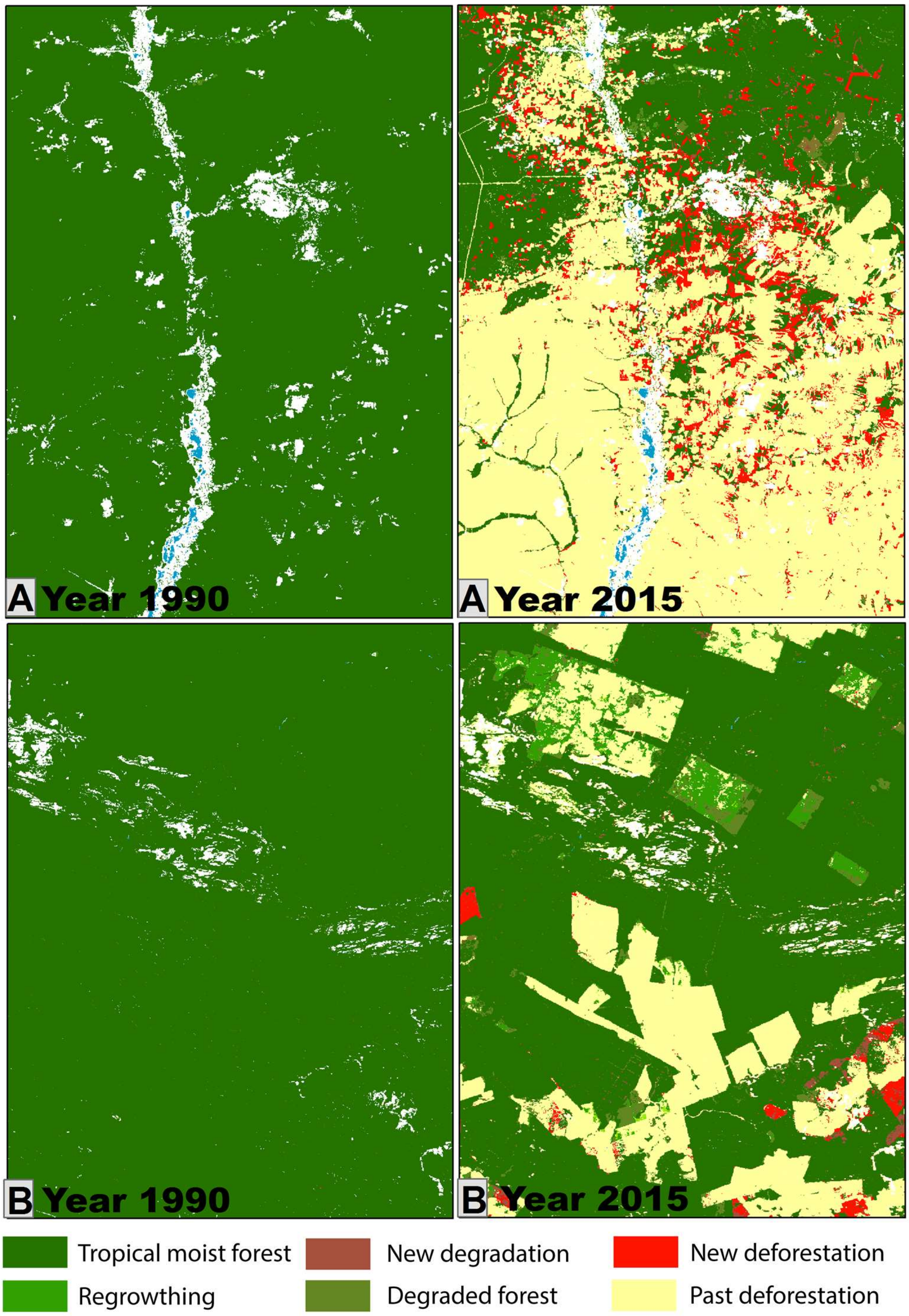

**Fig. S2** Methodological steps for the definition of the transition classes.

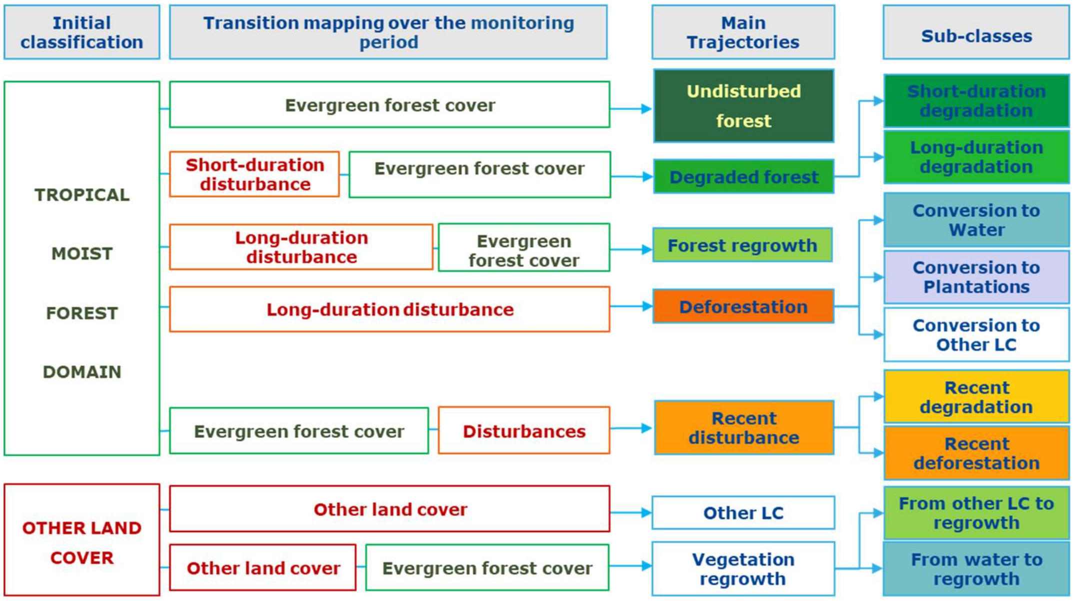

**Fig. S3** Subsets (10 km × 30 km) of the transition map capturing different types of logging areas: A, logging concession in Ouesso, the Republic of the Congo; B, selective logging in Para state, Brazil; C, logging network in Suriname; D, logging and deforestation in Papua New Guinea. Short-duration degradation (logging activities) appears in green and deforestation appears in red.

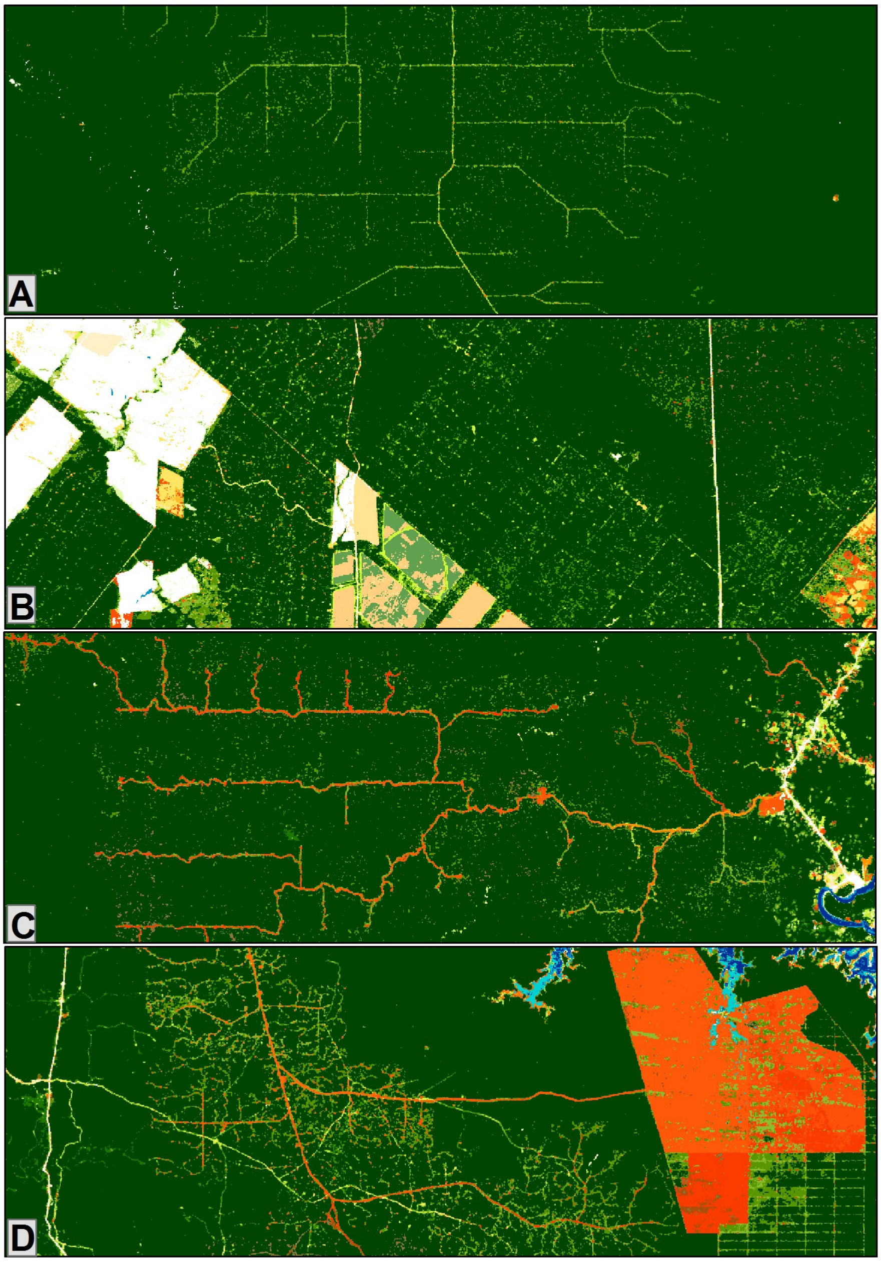

**Fig. S4** Subsets (18 km × 50 km) of the transition map capturing different types of deforestation processes: A, deforestation in the south of Porto Velho (Rondônia state, Brazil; B, deforestation in Roraima state, Brazil; C, deforestation and degradation due to the proximity of the railway in Cameroon; D, degradation and deforestation in a protected area in Ivory Coast.

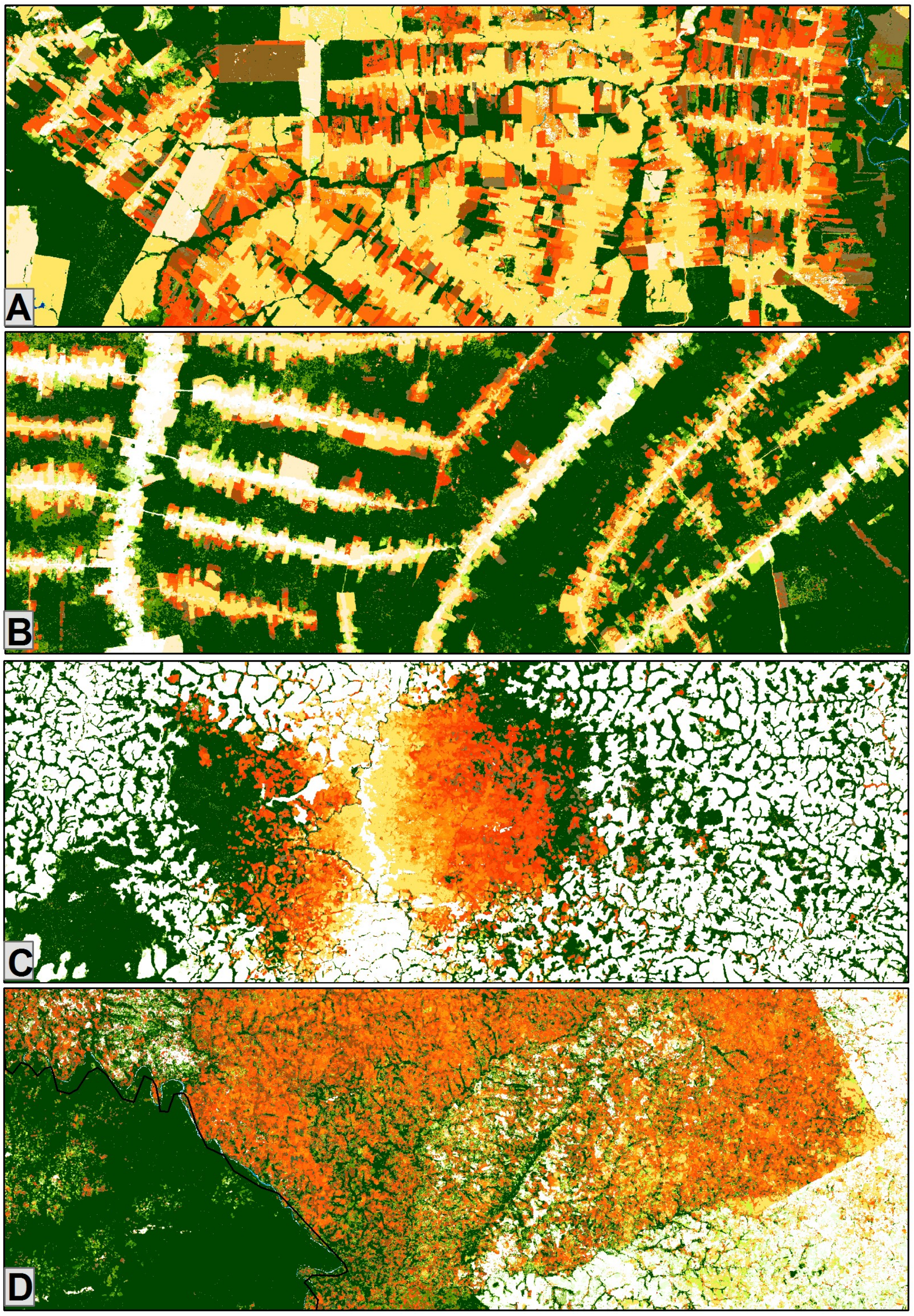

**Fig. S5** Subsets (18 km × 50 km) of the transition map capturing different tree plantation areas: A, cacao plantation in Venezuela; B, recent large oil palm plantation in Gabon (2015-2017); C, massive forest conversion to oil palm plantations in Cambodia; D, oil palm plantations in Indonesia.

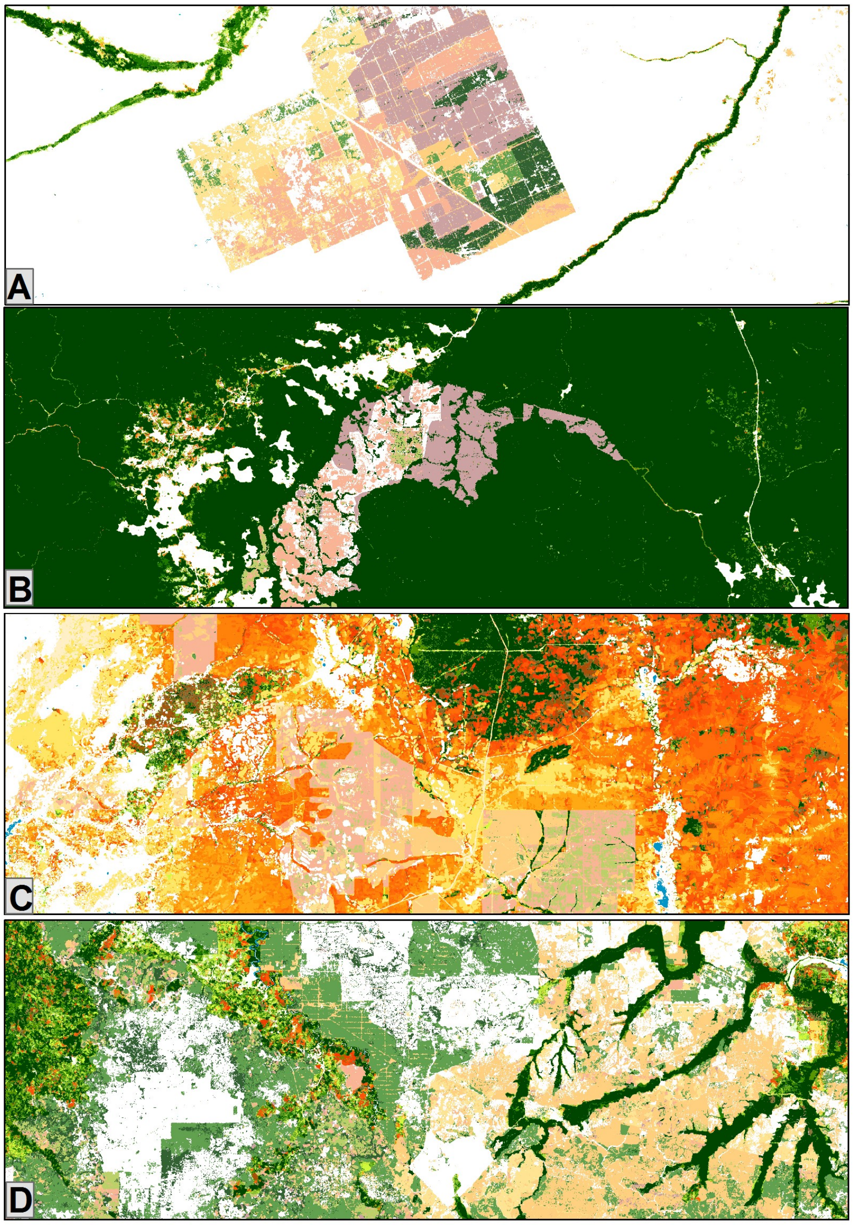

**Fig. S6** Subsets (18 km × 50 km) of the transition map: A, forest conversion to water body due to a new dam in Malaysia; B, road network in Sarawak, Malaysia, at the border with Kalimantan Indonesia; C, rural complex in the Democratic Republic of the Congo; D, Gold mining in Peru (Madre De Dios, Mazuco).

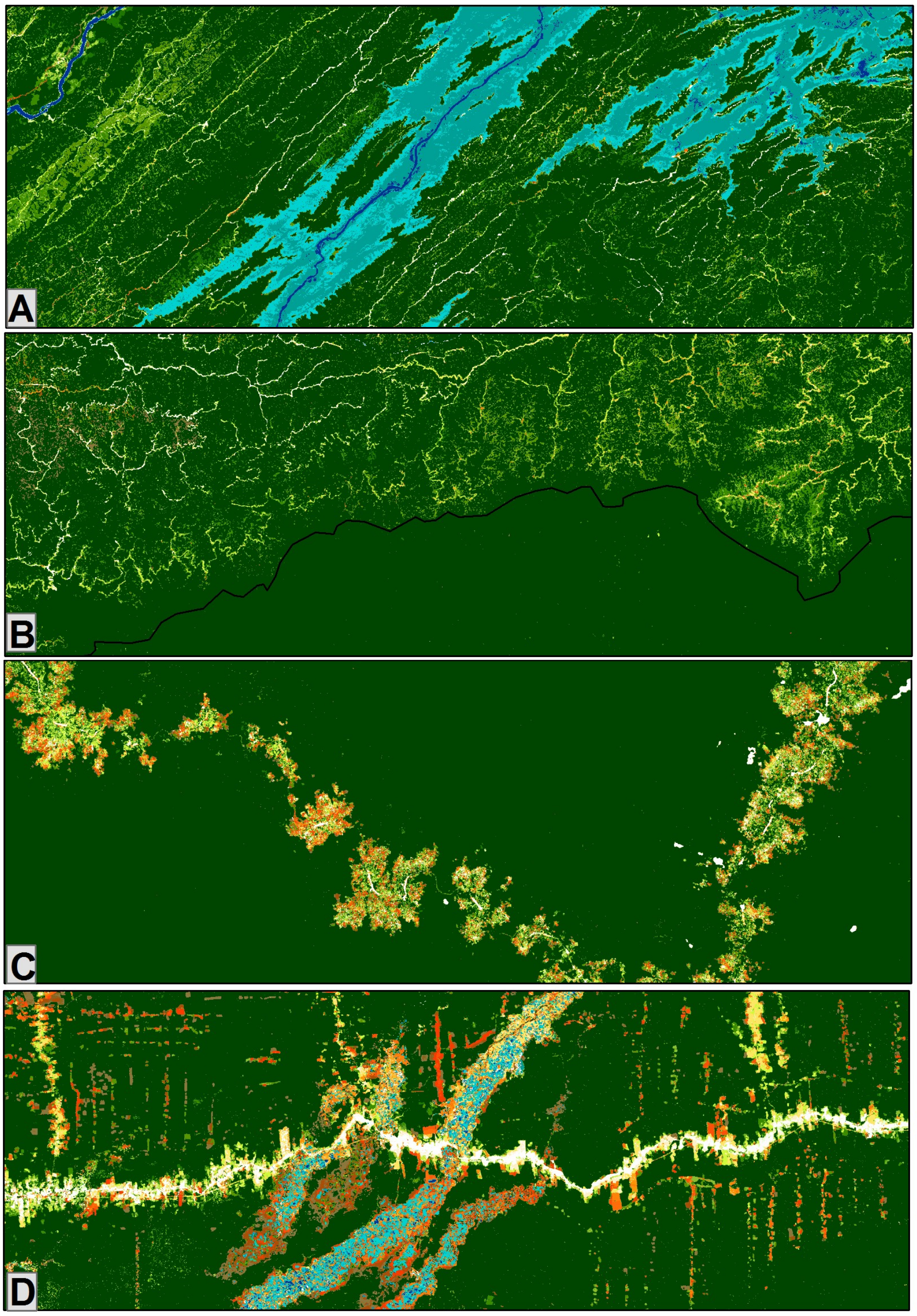

**Fig. S7** Subsets (20 X 50km) of the transition map capturing specific degradation patterns in tropical moist forests due to climatic events: A, Cyclone Debbie in 2017 in Northern Australia; B, droughts in 2016 due to ENSO events; C, Cyclone Hudah in Madagascar in April 2000 (north of Antalaha); D, fires related to droughts in the Amazon. Degraded forests appear in light green (if occurred before 2016) or brown (in 2016 or 2017).

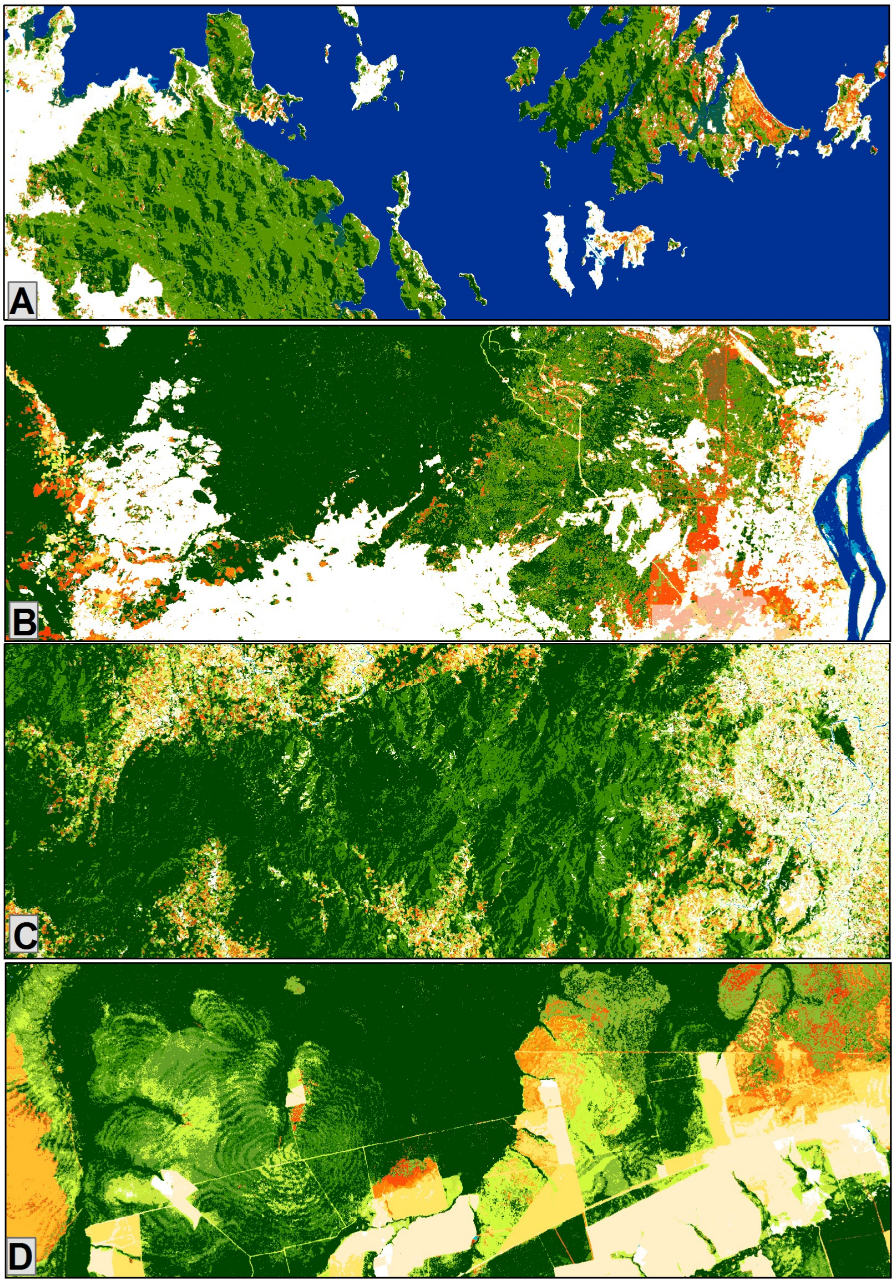

**Fig. S8** Sampling plots (5250) used for the validation and accuracy assessment.

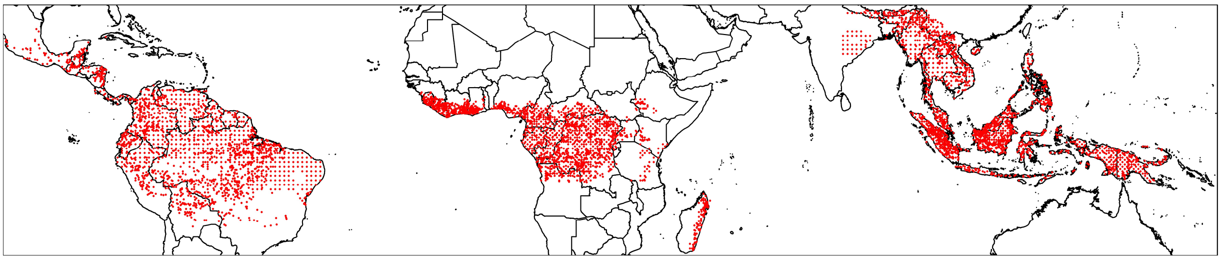

**Fig. S9** Validation response design: process of selection of dates of Landsat images within the seven strata and three periods, to be interpreted in the reference validation dataset.

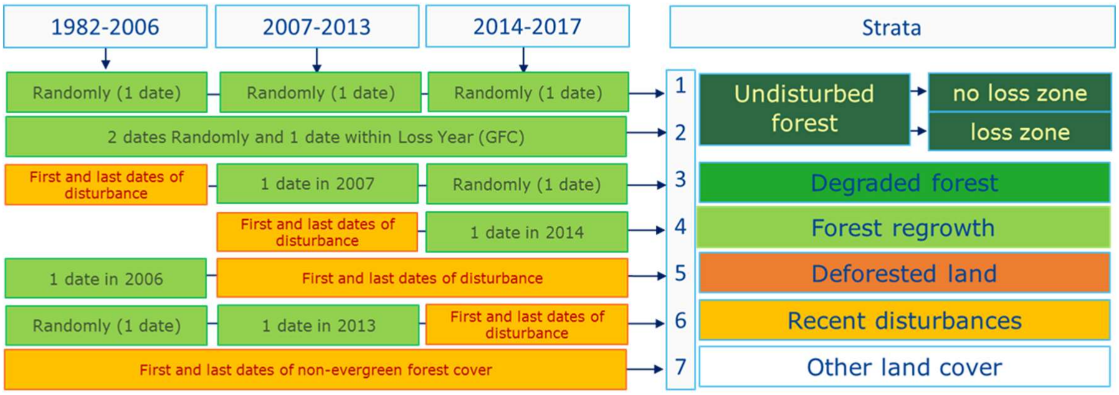

**Fig. S10** Example of systematic blocks of 1˚ by 1˚ longitude–latitude, and replicates 1 and 2 (one replicate is the set of points that occupy the same position in each block), over an area in the Democratic Republic of the Congo (around 350 km × 250 km in size and centred on 19.5˚E, 2˚N) with the transition layer in the background.

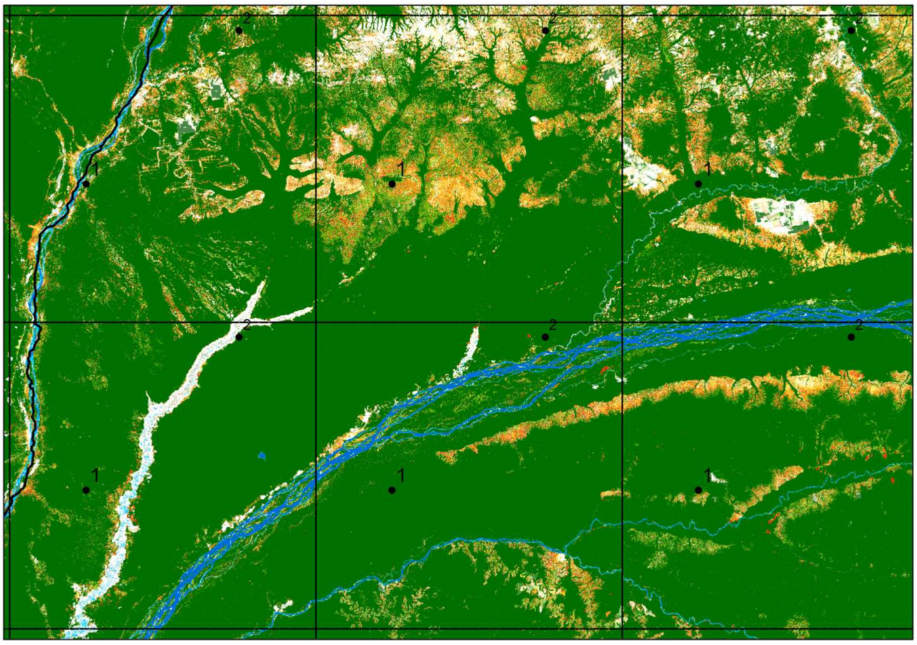

**Fig. S11** Annual disturbances from 1990 to 2019 for the seven subregions and the countries with an undisturbed forest area larger than 5 million ha in 1990.

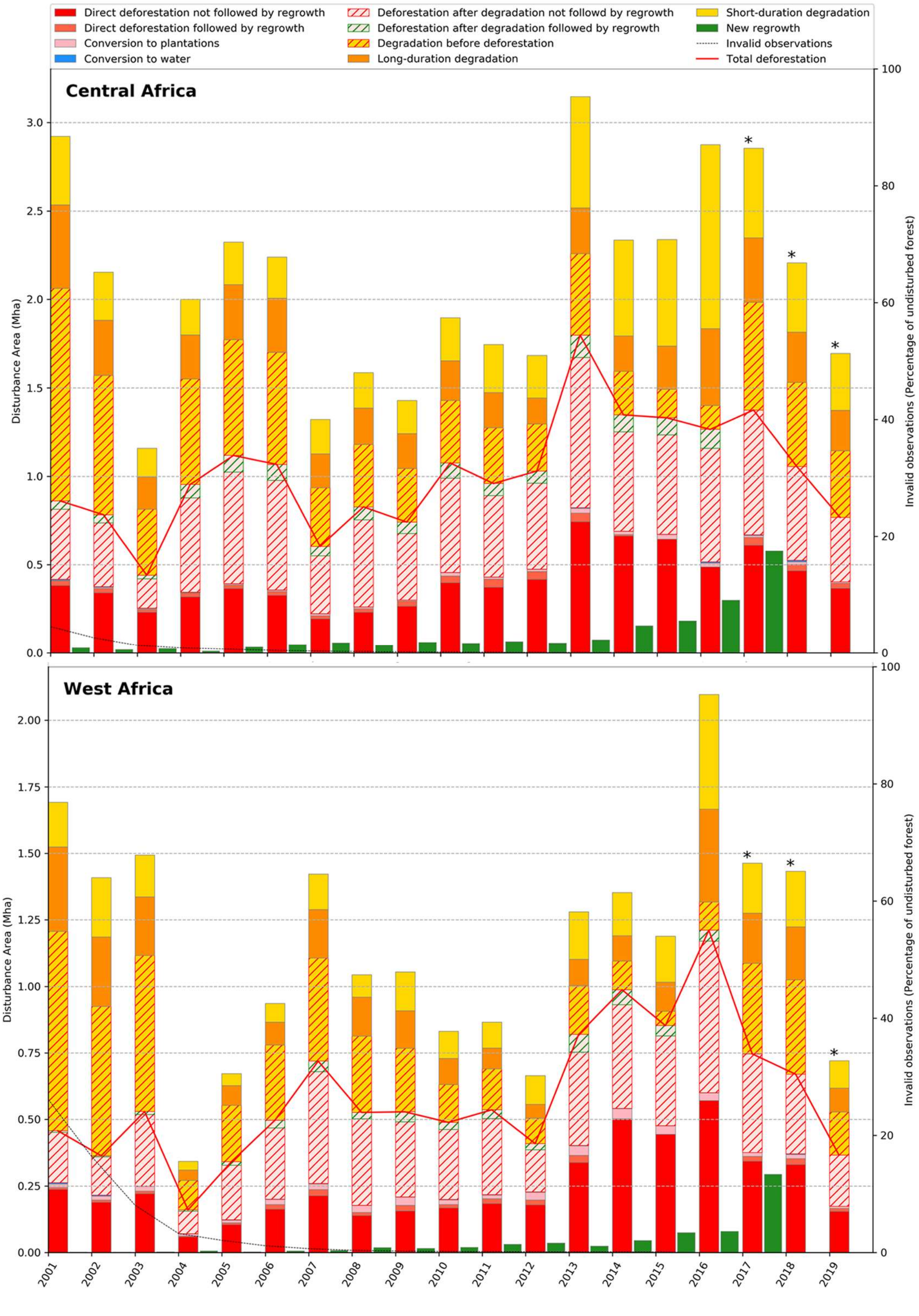

733

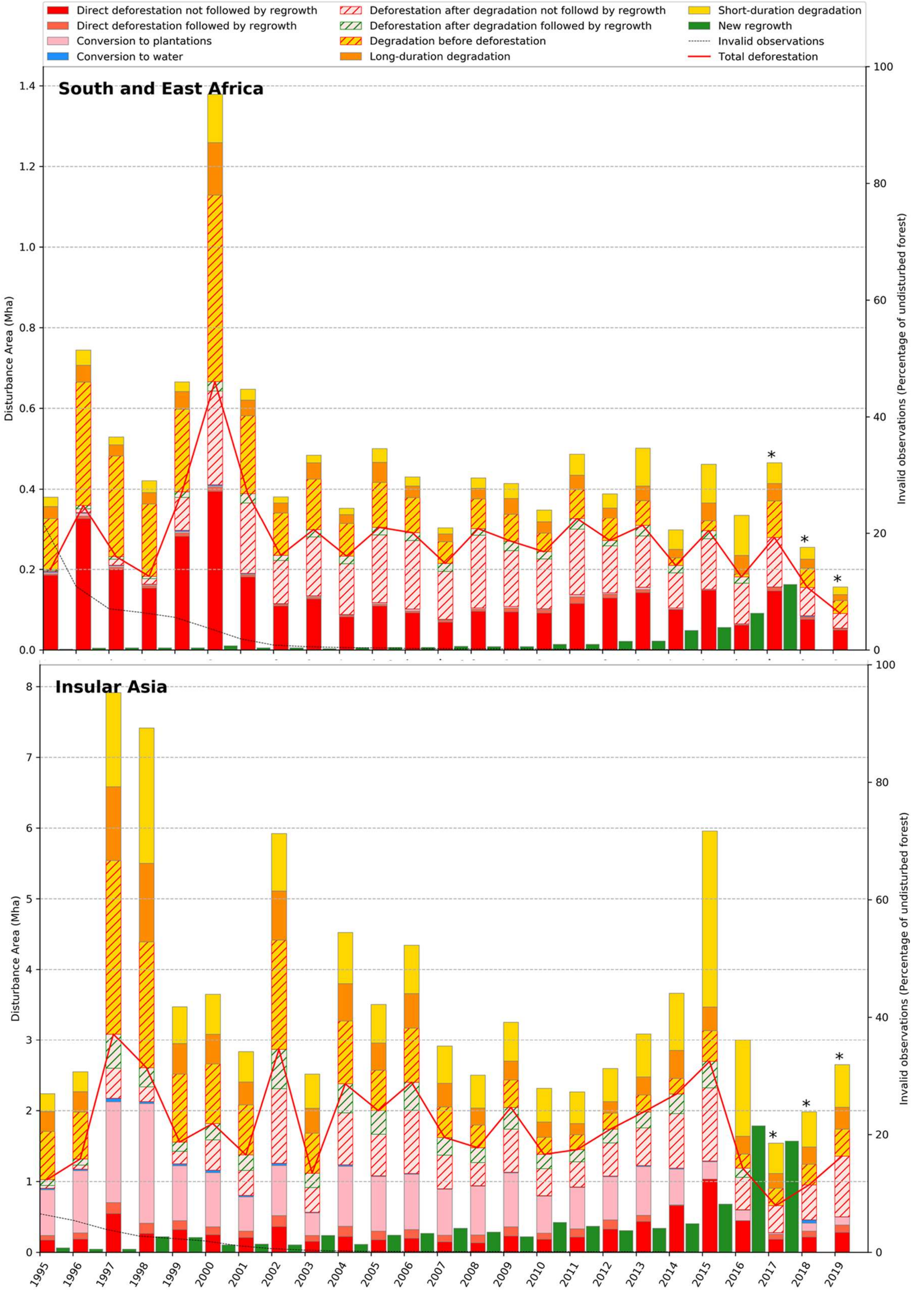

Direct deforestation not followed by regrowth
Direct deforestation followed by regrowth
Conversion to plantations
Conversion to water
Deforestation after degradation not followd by regrowth
Deforestation after degradation followed by regrowth
Degradation before deforestation
Long-duration degradation
Short-duration degradation
New regrowth
Invalid observations
Total deforestation
South and East Africa
Insular Asia
Disturbance Area (Mha)
Invalid observations (Percentage of undisturbed forest)

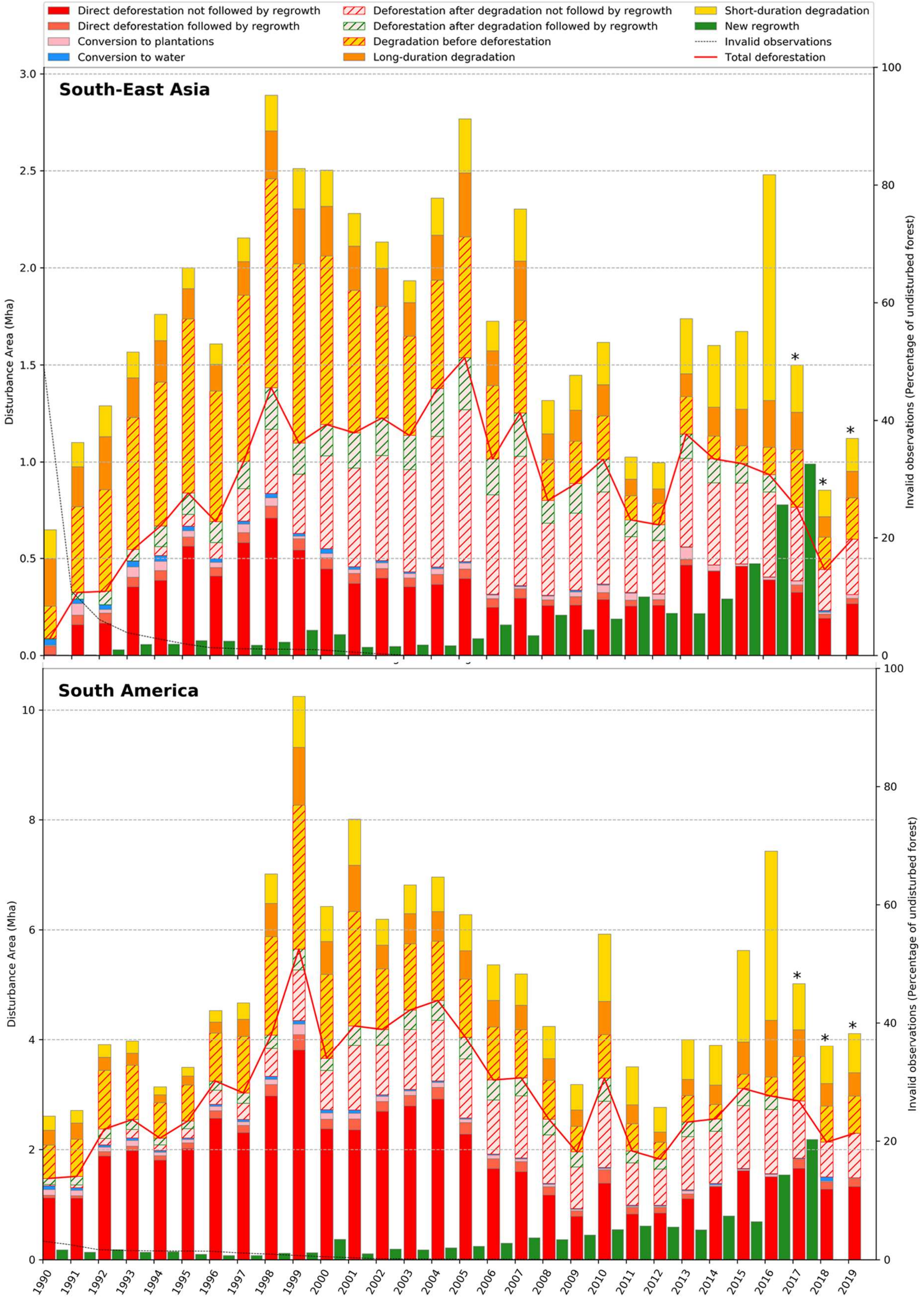

Direct deforestation not followed by regrowth
Direct deforestation followed by regrowth
Conversion to plantations
Conversion to water
Deforestation after degradation not followd by regrowth
Deforestation after degradation followed by regrowth
Degradation before deforestation
Long-duration degradation
Short-duration degradation
New regrowth
Invalid observations
Total deforestation
South-East Asia
South America
Disturbance Area (Mha)
Invalid observations (Percentage of undisturbed forest)


734

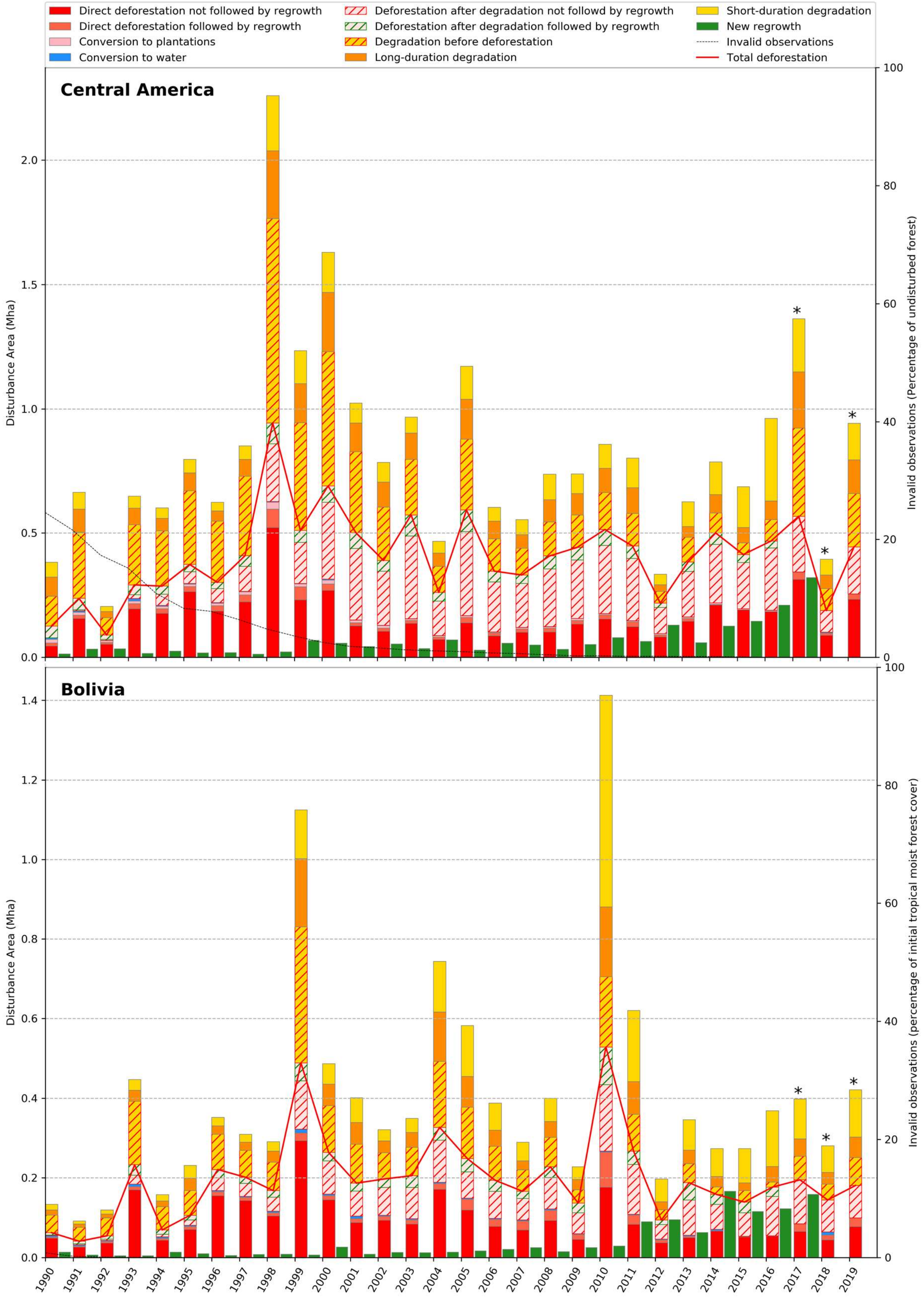
Direct deforestation not followed by regrowth
Direct deforestation followed by regrowth
Conversion to plantations
Conversion to water
Deforestation after degradation not followd by regrowth
Deforestation after degradation followed by regrowth
Degradation before deforestation
Long-duration degradation
Short-duration degradation
New regrowth
Invalid observations
Total deforestation
Central America
Disturbance Area (Mha)
Invalid observations (Percentage of undisturbed forest)
Bolivia
Disturbance Area (Mha)
Invalid observations (percentage of initial tropical moist forest cover)
1990 1991 1992 1993 1994 1995 1996 1997 1998 1999 2000 2001 2002 2003 2004 2005 2006 2007 2008 2009 2010 2011 2012 2013 2014 2015 2016 2017 2018 2019

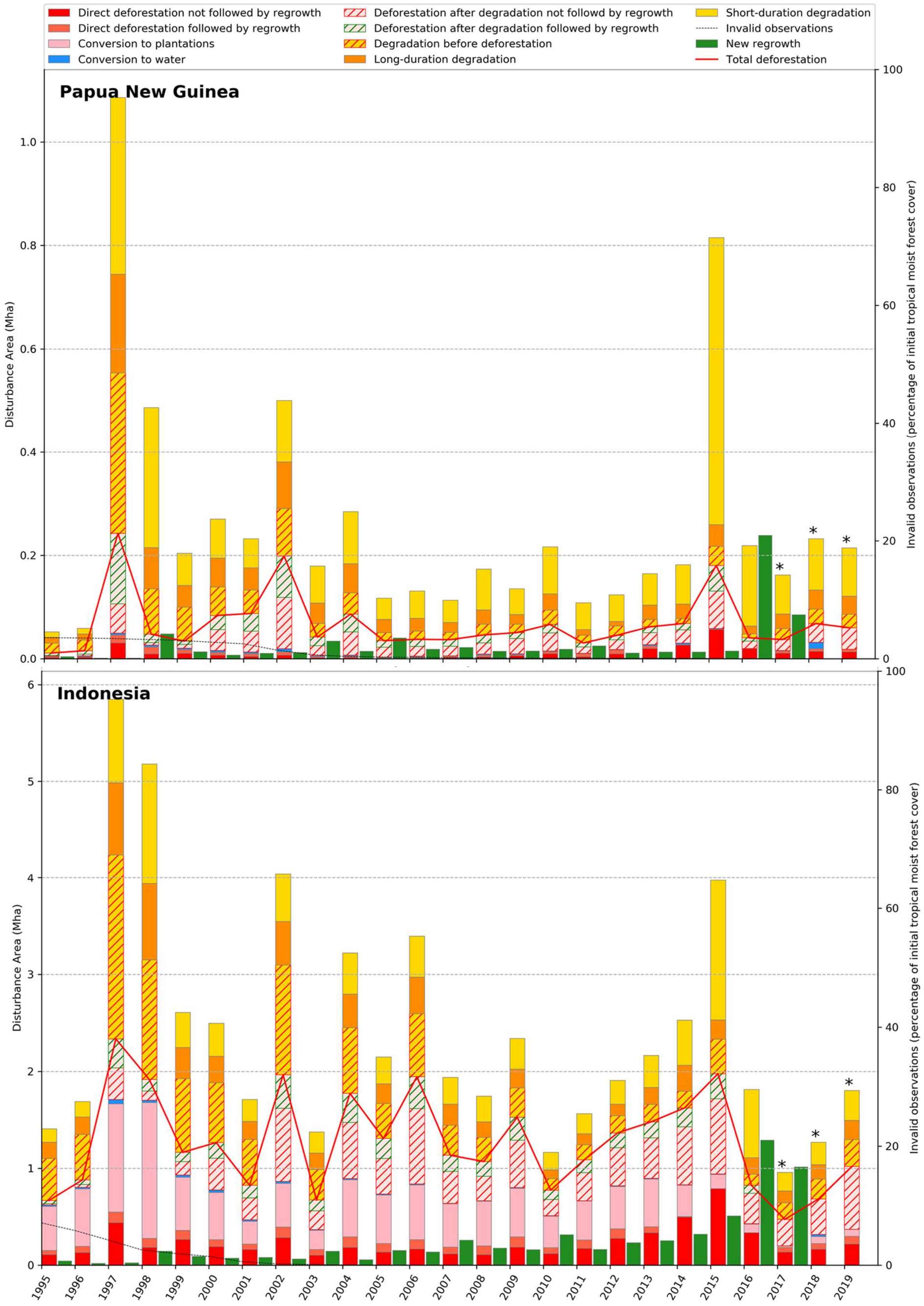

Direct deforestation not followed by regrowth
Direct deforestation followed by regrowth
Conversion to plantations
Conversion to water
Deforestation after degradation not followd by regrowth
Deforestation after degradation followed by regrowth
Degradation before deforestation
Long-duration degradation
Short-duration degradation
Invalid observations
New regrowth
Total deforestation
Papua New Guinea
Indonesia
Disturbance Area (Mha)
Invalid observations (percentage of initial tropical moist forest cover)


737

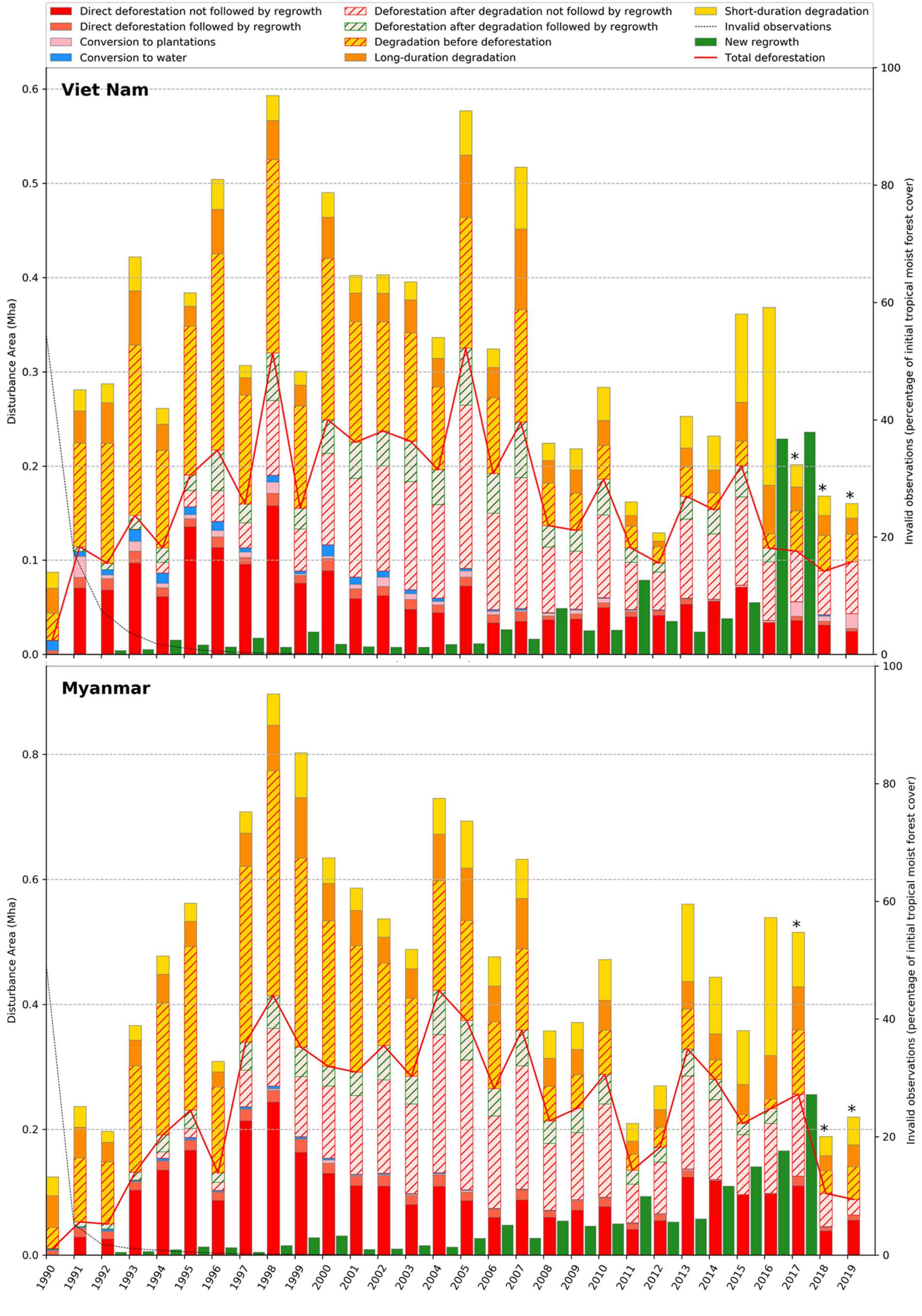
Direct deforestation not followed by regrowth
Direct deforestation followed by regrowth
Conversion to plantations
Conversion to water
Deforestation after degradation not followd by regrowth
Deforestation after degradation followed by regrowth
Degradation before deforestation
Long-duration degradation
Short-duration degradation
Invalid observations
New regrowth
Total deforestation
Viet Nam
Myanmar
Disturbance Area (Mha)
Invalid observations (percentage of initial tropical moist forest cover)
*
1990 1991 1992 1993 1994 1995 1996 1997 1998 1999 2000 2001 2002 2003 2004 2005 2006 2007 2008 2009 2010 2011 2012 2013 2014 2015 2016 2017 2018 2019

739

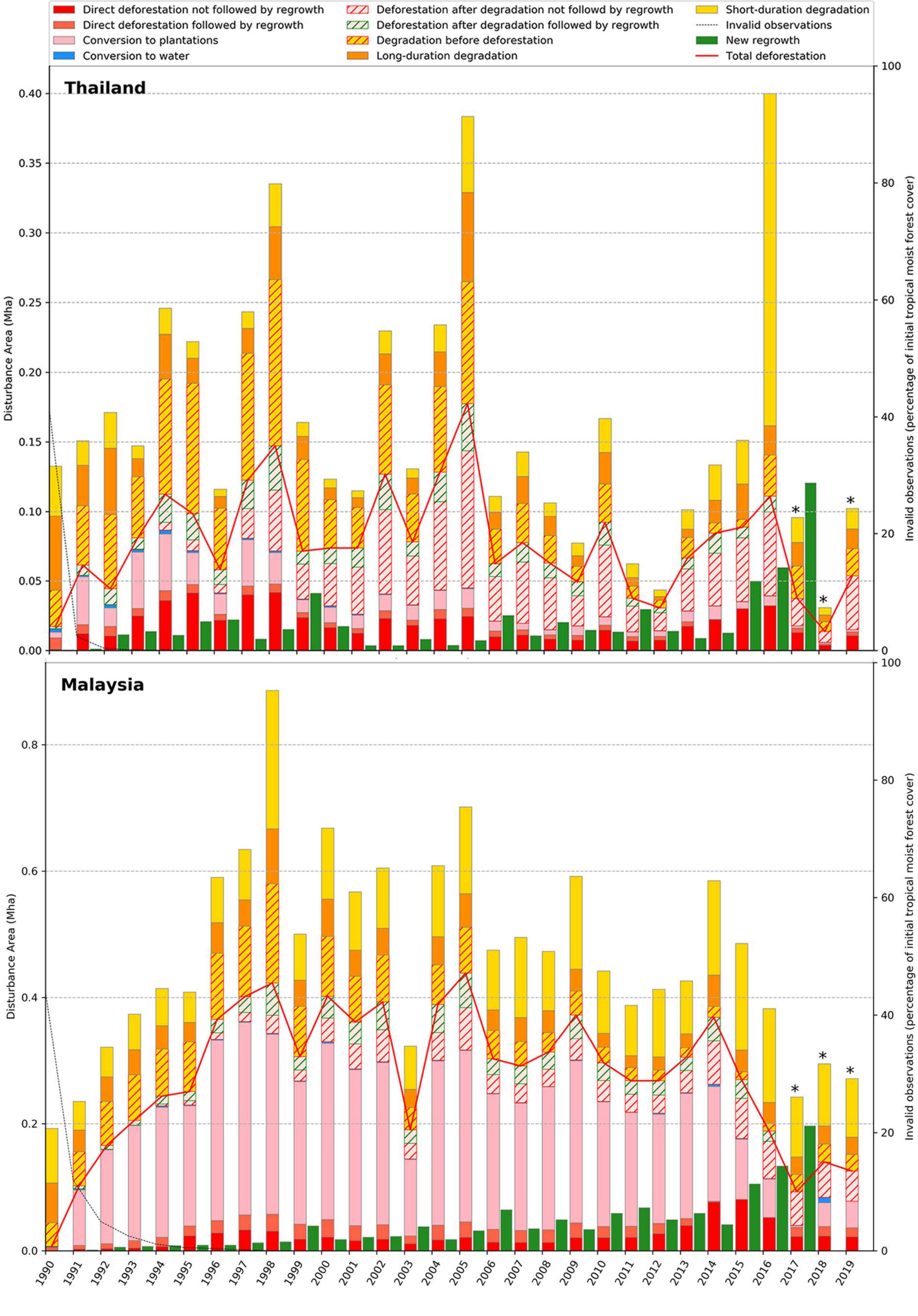

Direct deforestation not followed by regrowth
Direct deforestation followed by regrowth
Conversion to plantations
Conversion to water
Deforestation after degradation not followd by regrowth
Deforestation after degradation followed by regrowth
Degradation before deforestation
Long-duration degradation
Short-duration degradation
Invalid observations
New regrowth
Total deforestation
Thailand
Malaysia
Disturbance Area (Mha)
Invalid observations (percentage of initial tropical moist forest cover)

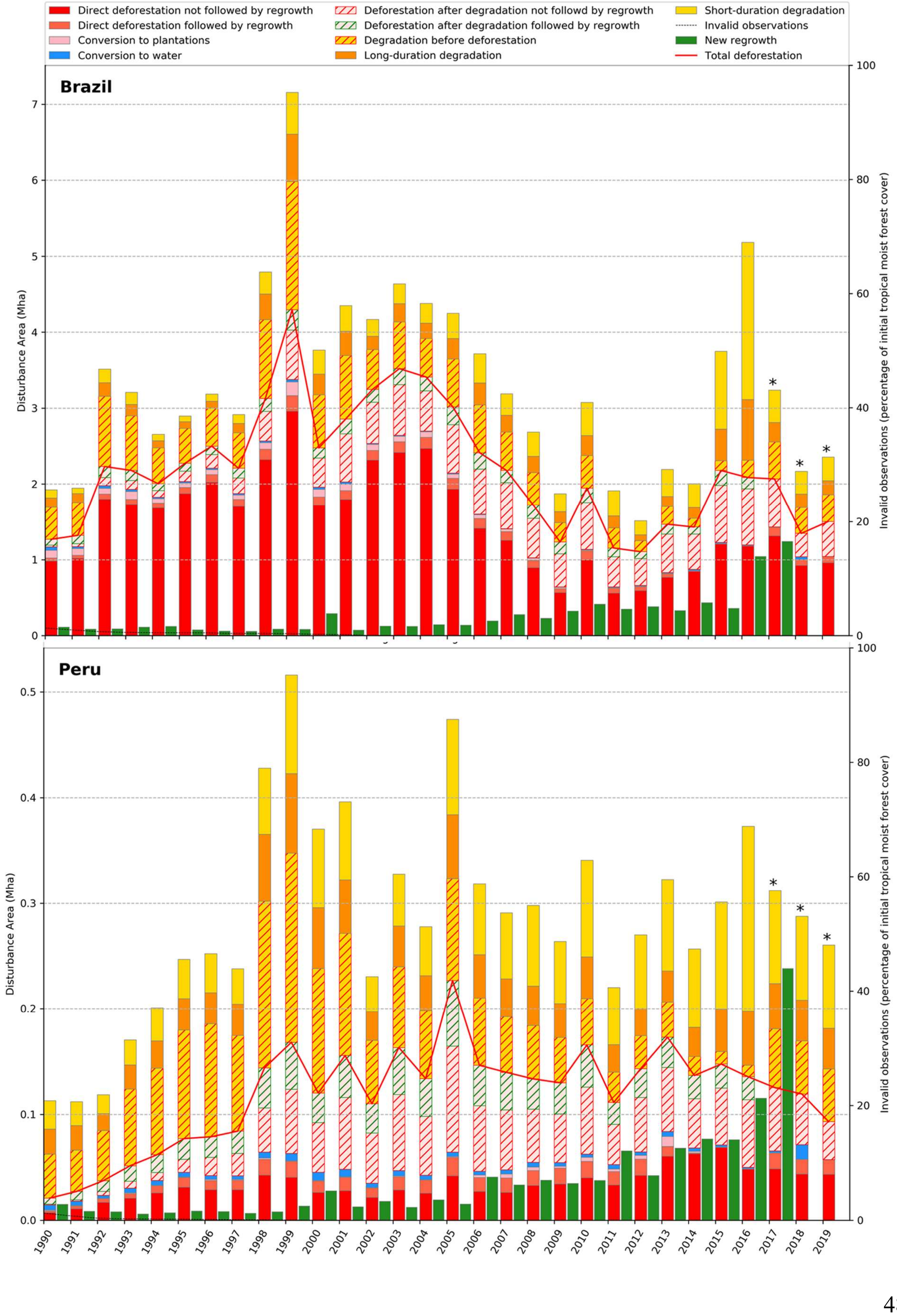

Direct deforestation not followed by regrowth
Direct deforestation followed by regrowth
Conversion to plantations
Conversion to water
Deforestation after degradation not followd by regrowth
Deforestation after degradation followed by regrowth
Degradation before deforestation
Long-duration degradation
Short-duration degradation
Invalid observations
New regrowth
Total deforestation
Brazil
Peru
Disturbance Area (Mha)
Invalid observations (percentage of initial tropical moist forest cover)


740

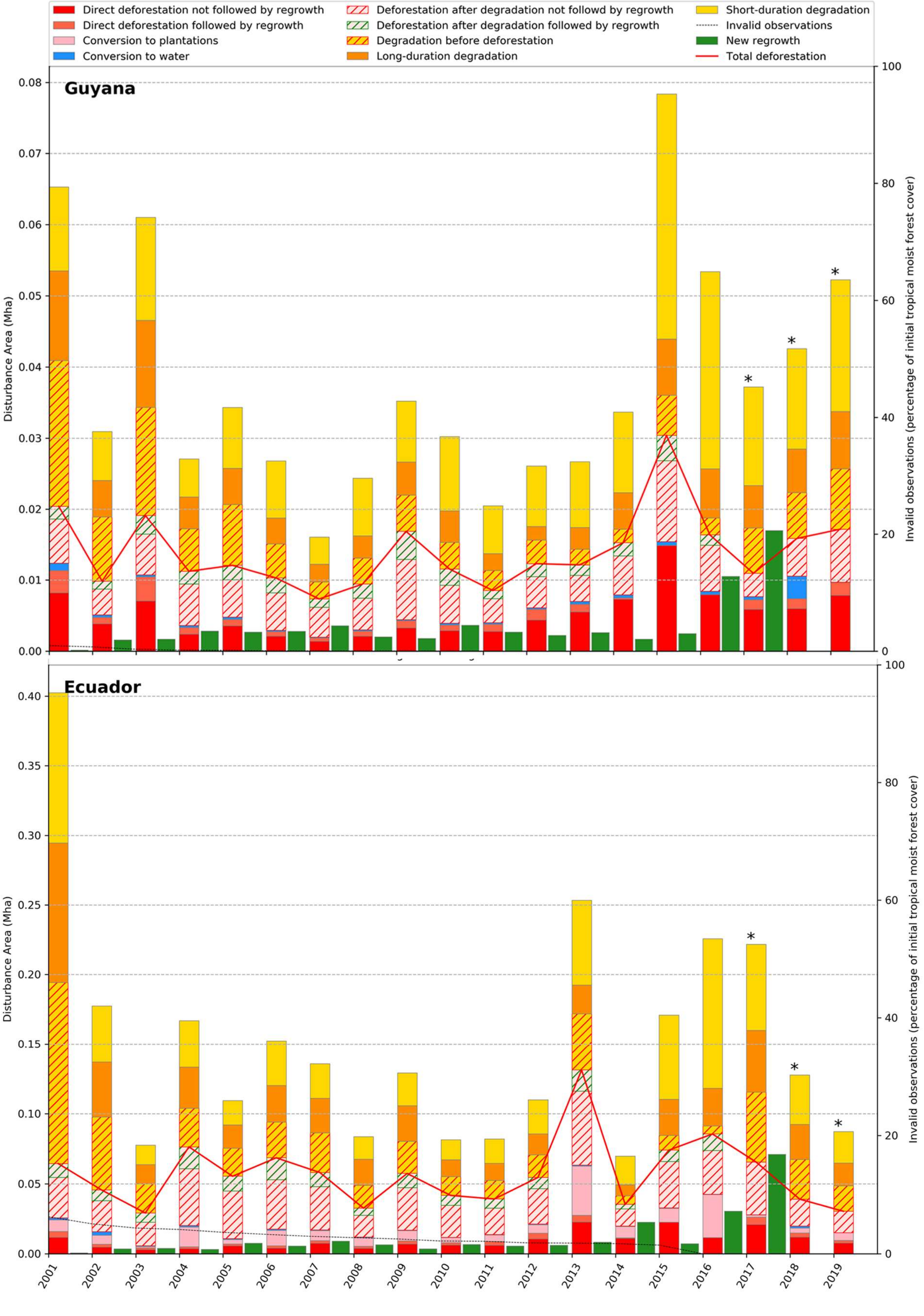

Direct deforestation not followed by regrowth
Direct deforestation followed by regrowth
Conversion to plantations
Conversion to water
Deforestation after degradation not followd by regrowth
Deforestation after degradation followed by regrowth
Degradation before deforestation
Long-duration degradation
Short-duration degradation
Invalid observations
New regrowth
Total deforestation
Guyana
Ecuador
Disturbance Area (Mha)
Invalid observations (percentage of initial tropical moist forest cover)
2001 2002 2003 2004 2005 2006 2007 2008 2009 2010 2011 2012 2013 2014 2015 2016 2017 2018 2019


741

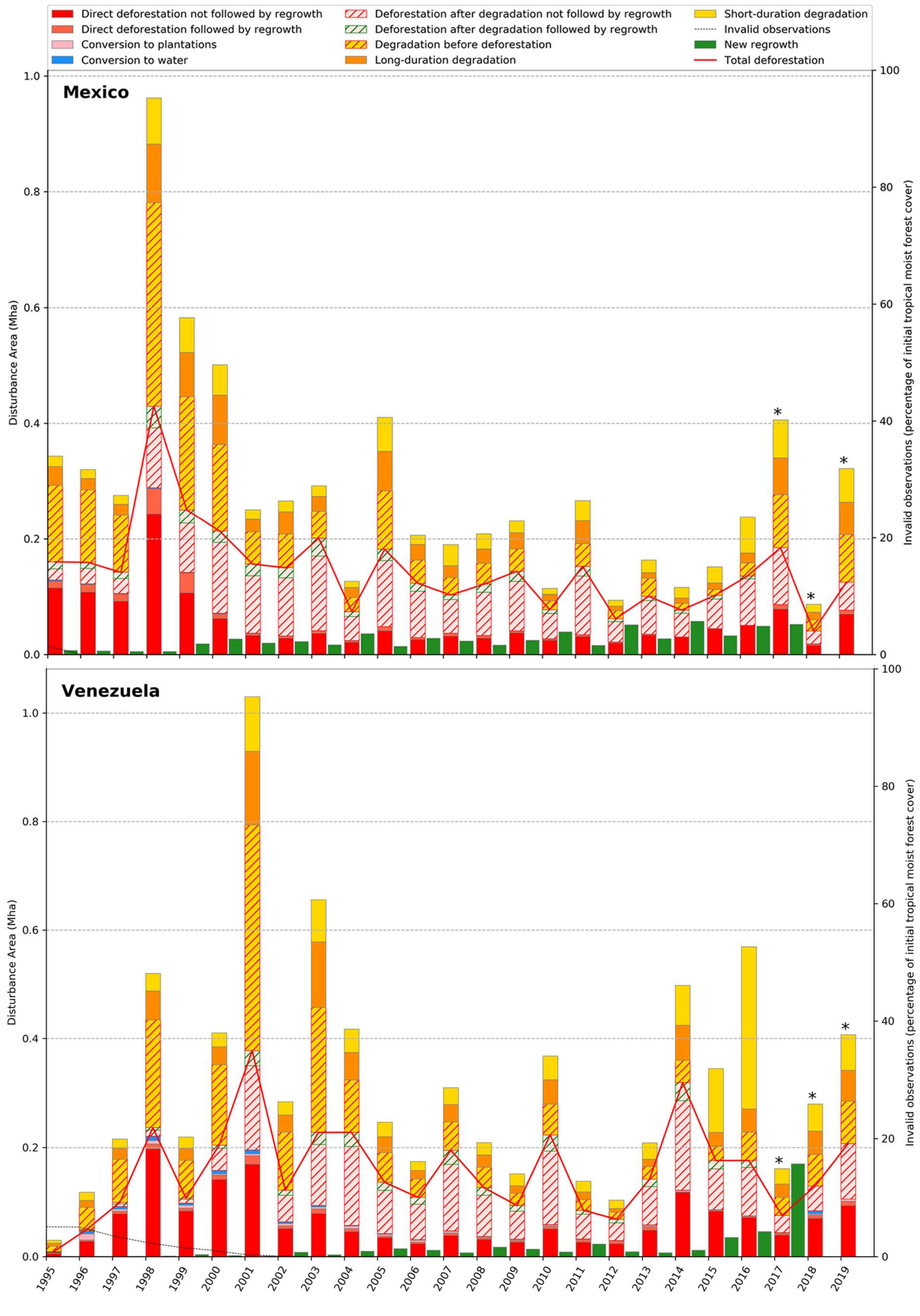

Direct deforestation not followed by regrowth
Direct deforestation followed by regrowth
Conversion to plantations
Conversion to water
Deforestation after degradation not followd by regrowth
Deforestation after degradation followed by regrowth
Degradation before deforestation
Long-duration degradation
Short-duration degradation
Invalid observations
New regrowth
Total deforestation
Mexico
Venezuela
Disturbance Area (Mha)
Invalid observations (percentage of initial tropical moist forest cover)


742

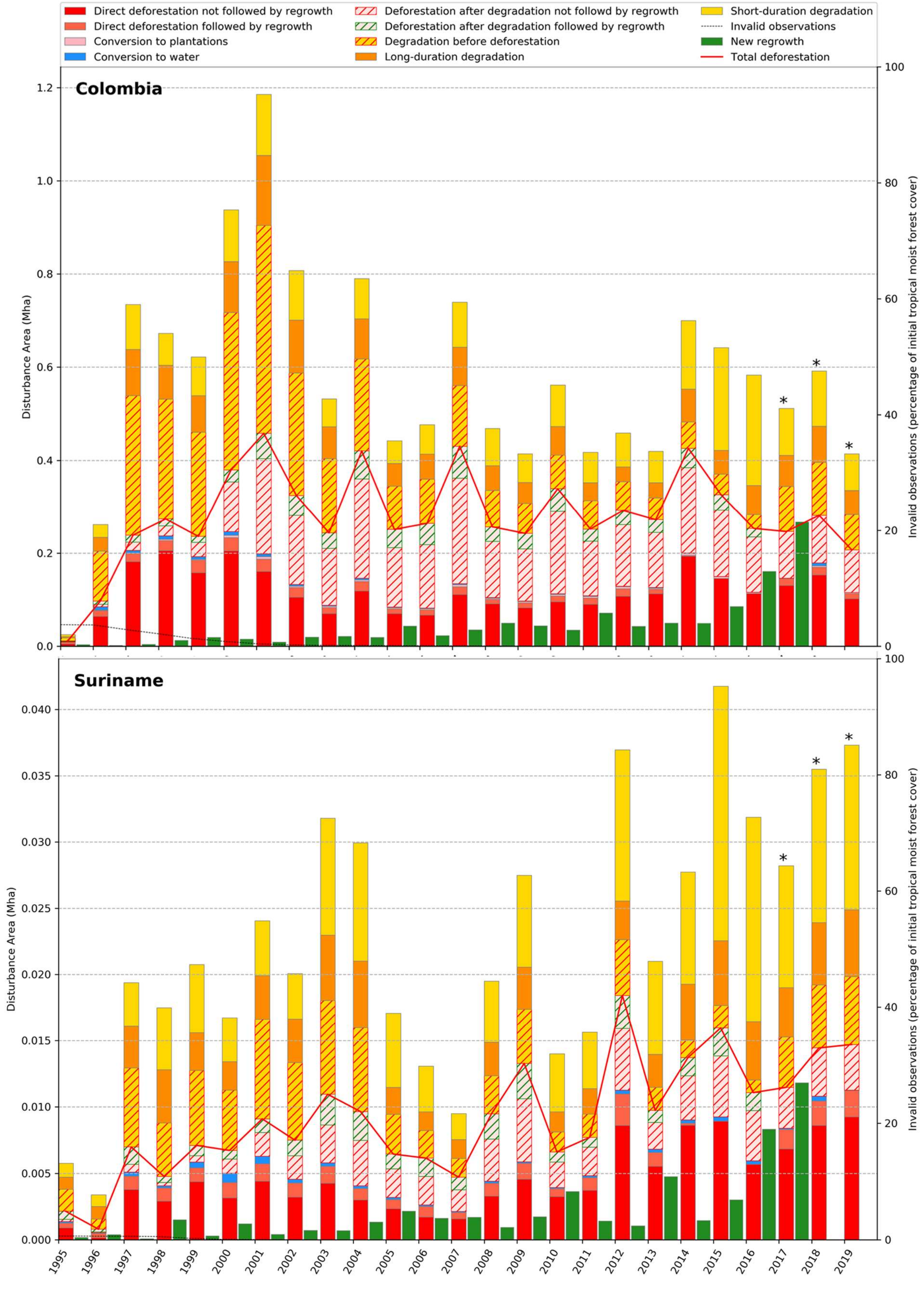
Direct deforestation not followed by regrowth
Direct deforestation followed by regrowth
Conversion to plantations
Conversion to water
Deforestation after degradation not followd by regrowth
Deforestation after degradation followed by regrowth
Degradation before deforestation
Long-duration degradation
Short-duration degradation
Invalid observations
New regrowth
Total deforestation
Colombia
Suriname
Disturbance Area (Mha)
Invalid observations (percentage of initial tropical moist forest cover)
*
1995
1996
1997
1998
1999
2000
2001
2002
2003
2004
2005
2006
2007
2008
2009
2010
2011
2012
2013
2014
2015
2016
2017
2018
2019

743

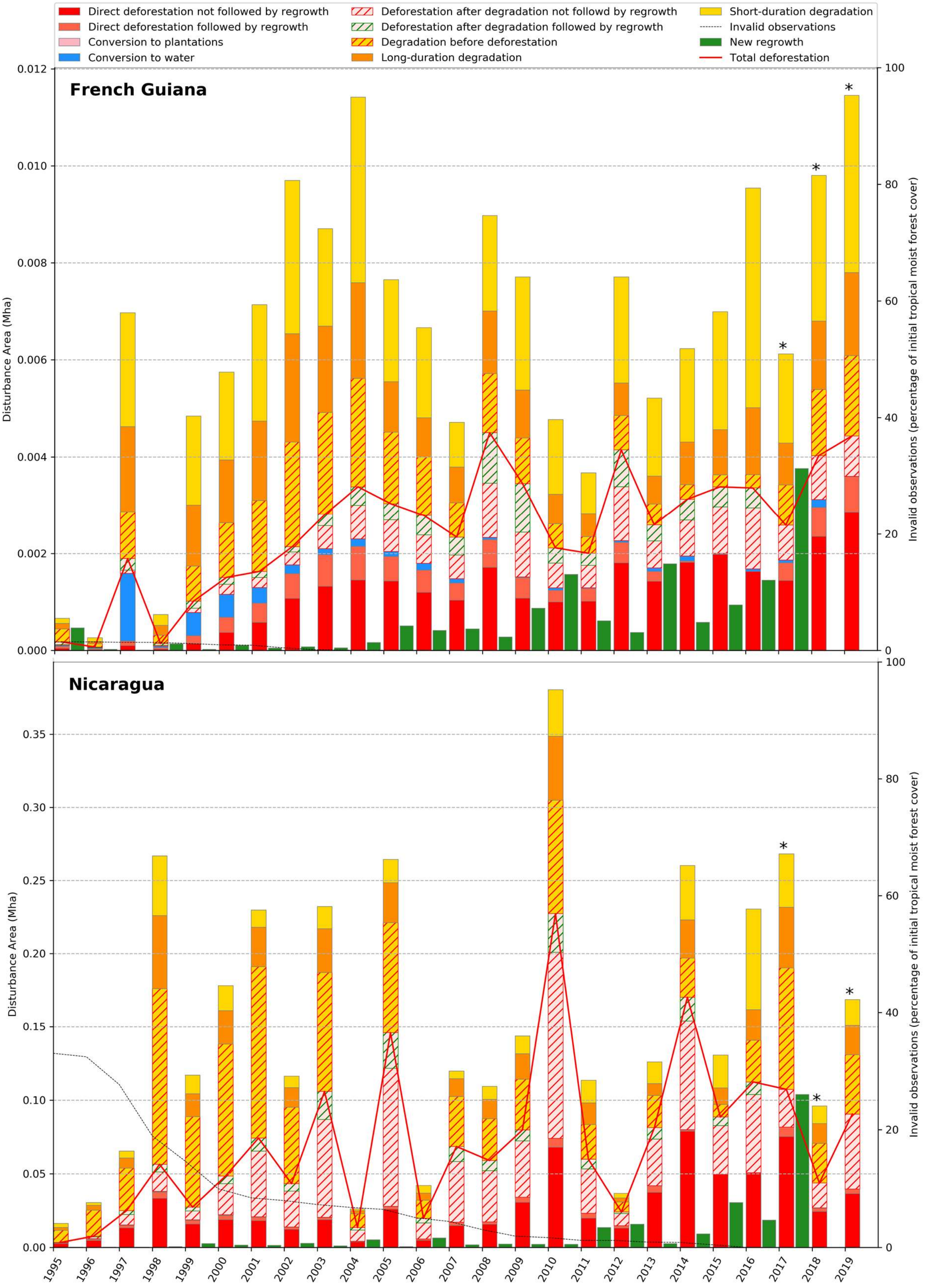
Direct deforestation not followed by regrowth
Direct deforestation followed by regrowth
Conversion to plantations
Conversion to water
Deforestation after degradation not followd by regrowth
Deforestation after degradation followed by regrowth
Degradation before deforestation
Long-duration degradation
Short-duration degradation
Invalid observations
New regrowth
Total deforestation
French Guiana
Nicaragua
Disturbance Area (Mha)
Invalid observations (percentage of initial tropical moist forest cover)
*

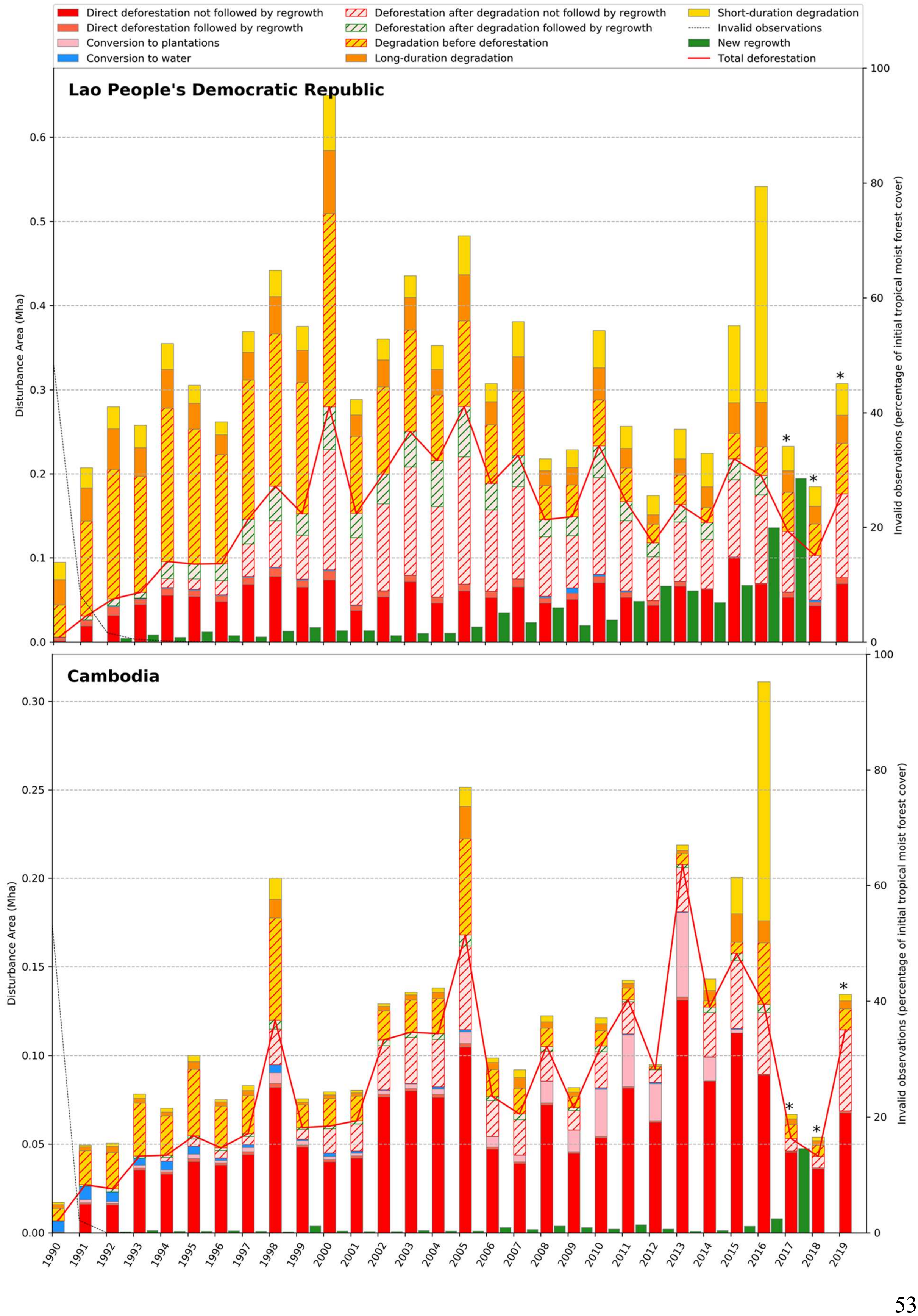

Direct deforestation not followed by regrowth
Direct deforestation followed by regrowth
Conversion to plantations
Conversion to water
Deforestation after degradation not followd by regrowth
Deforestation after degradation followed by regrowth
Degradation before deforestation
Long-duration degradation
Short-duration degradation
Invalid observations
New regrowth
Total deforestation
Lao People's Democratic Republic
Cambodia
Disturbance Area (Mha)
Invalid observations (percentage of initial tropical moist forest cover)

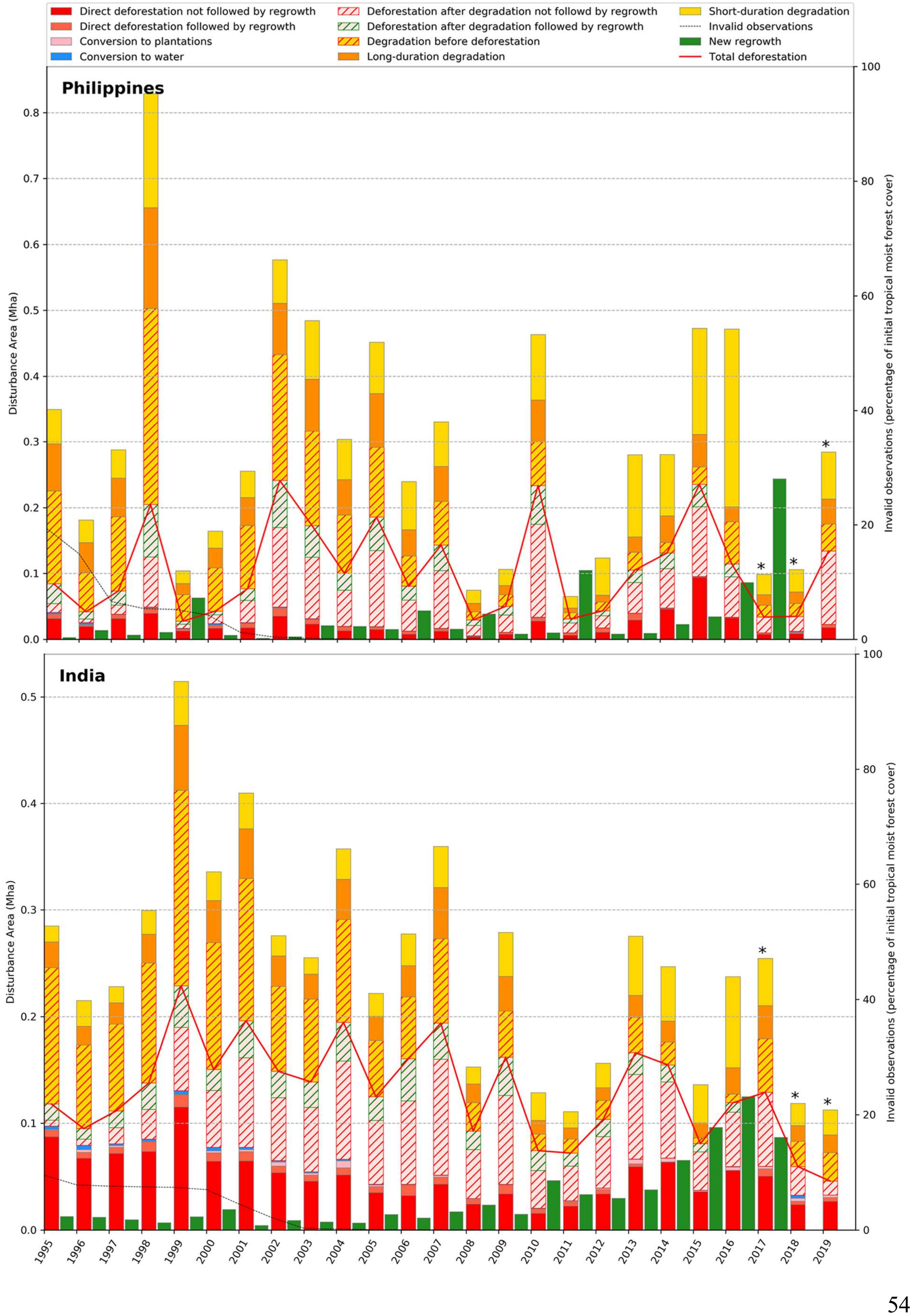

Direct deforestation not followed by regrowth
Direct deforestation followed by regrowth
Conversion to plantations
Conversion to water
Deforestation after degradation not followd by regrowth
Deforestation after degradation followed by regrowth
Degradation before deforestation
Long-duration degradation
Short-duration degradation
Invalid observations
New regrowth
Total deforestation
Philippines
India
Disturbance Area (Mha)
Invalid observations (percentage of initial tropical moist forest cover)


747

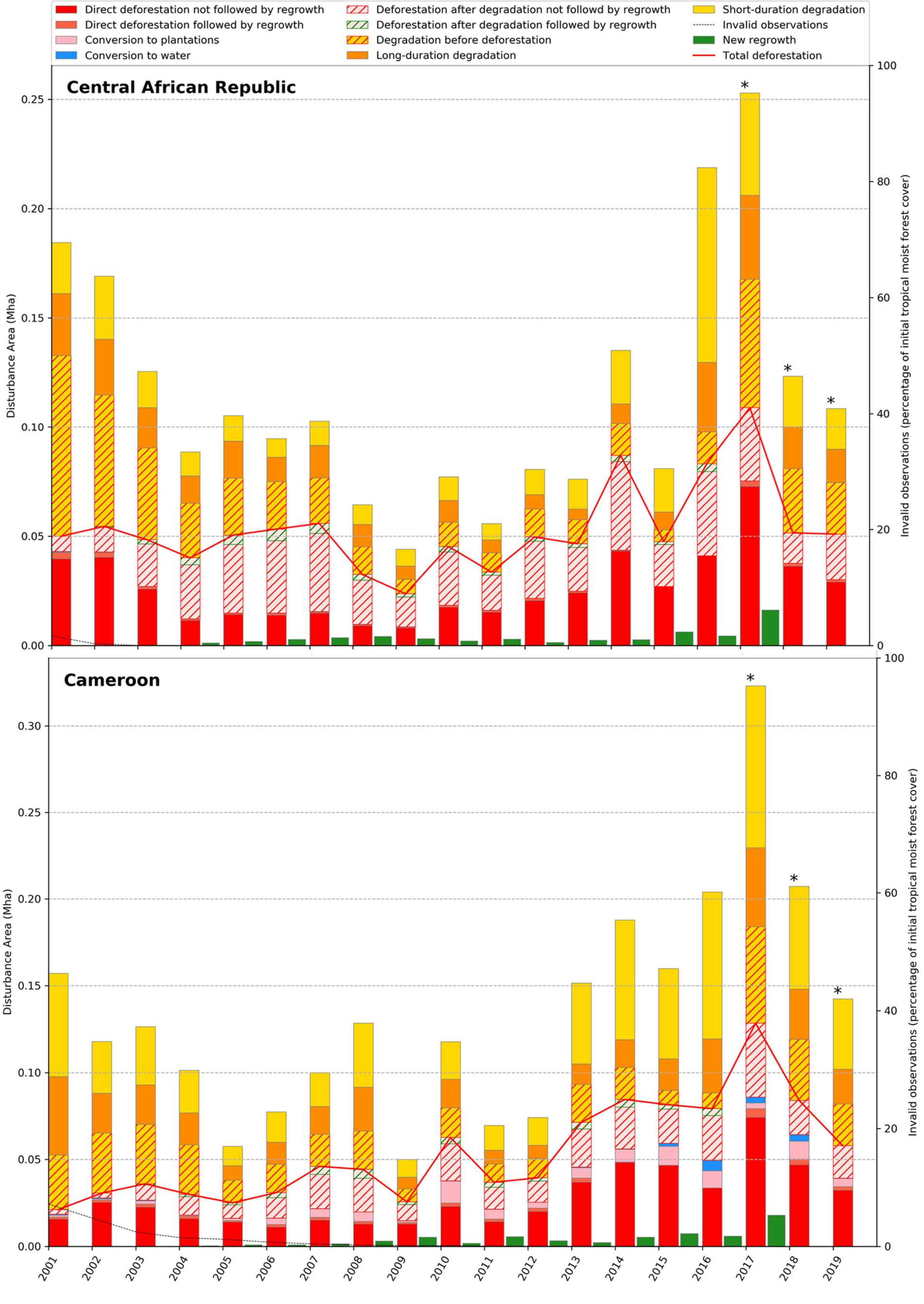

Direct deforestation not followed by regrowth
Direct deforestation followed by regrowth
Conversion to plantations
Conversion to water
Deforestation after degradation not followd by regrowth
Deforestation after degradation followed by regrowth
Degradation before deforestation
Long-duration degradation
Short-duration degradation
Invalid observations
New regrowth
Total deforestation
Central African Republic
Cameroon
Disturbance Area (Mha)
Invalid observations (percentage of initial tropical moist forest cover)
2001
2002
2003
2004
2005
2006
2007
2008
2009
2010
2011
2012
2013
2014
2015
2016
2017
2018
2019


748

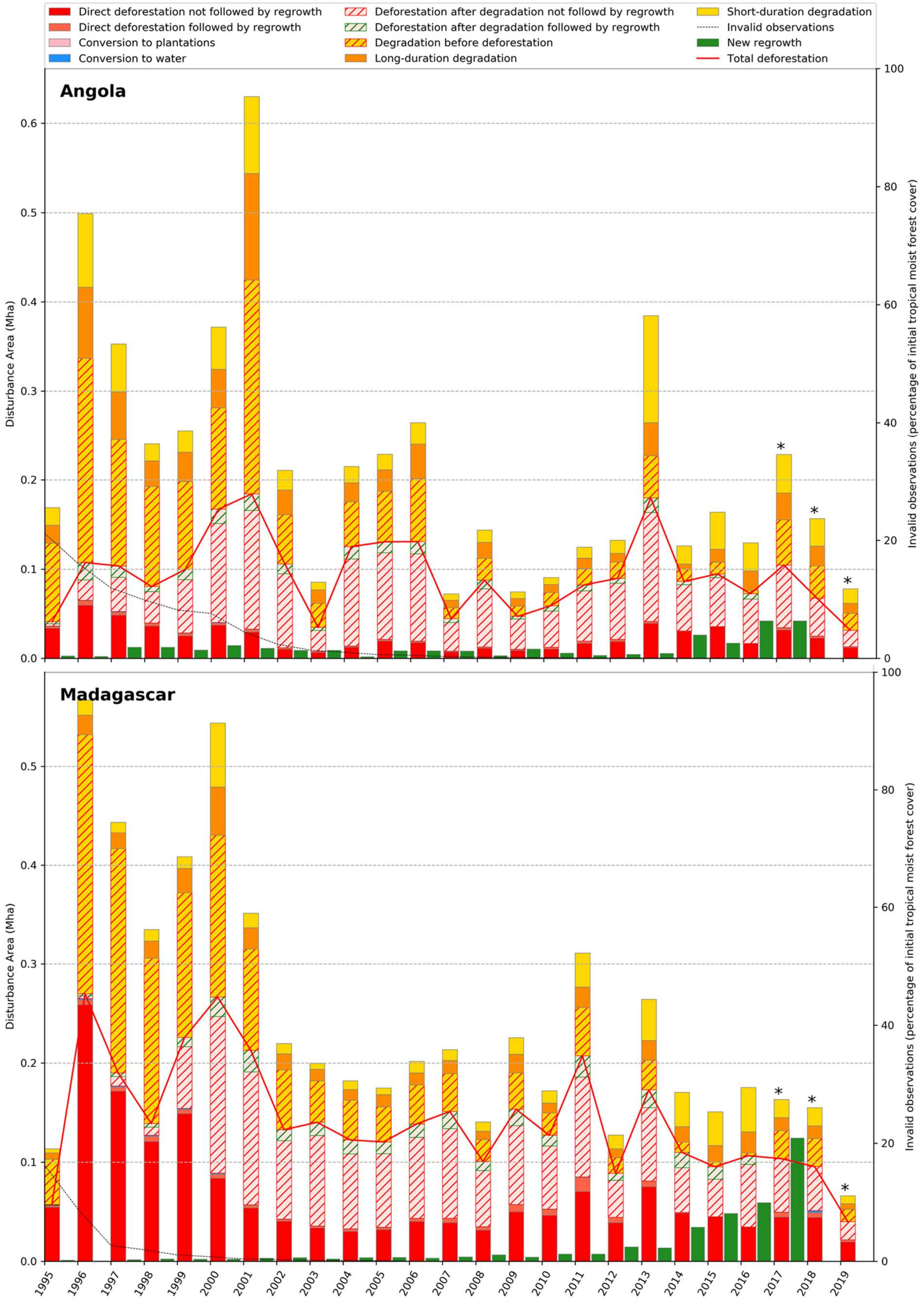
Direct deforestation not followed by regrowth
Direct deforestation followed by regrowth
Conversion to plantations
Conversion to water
Deforestation after degradation not followd by regrowth
Deforestation after degradation followed by regrowth
Degradation before deforestation
Long-duration degradation
Short-duration degradation
Invalid observations
New regrowth
Total deforestation
Angola
Madagascar
Disturbance Area (Mha)
Invalid observations (percentage of initial tropical moist forest cover)
1995
1996
1997
1998
1999
2000
2001
2002
2003
2004
2005
2006
2007
2008
2009
2010
2011
2012
2013
2014
2015
2016
2017
2018
2019

749

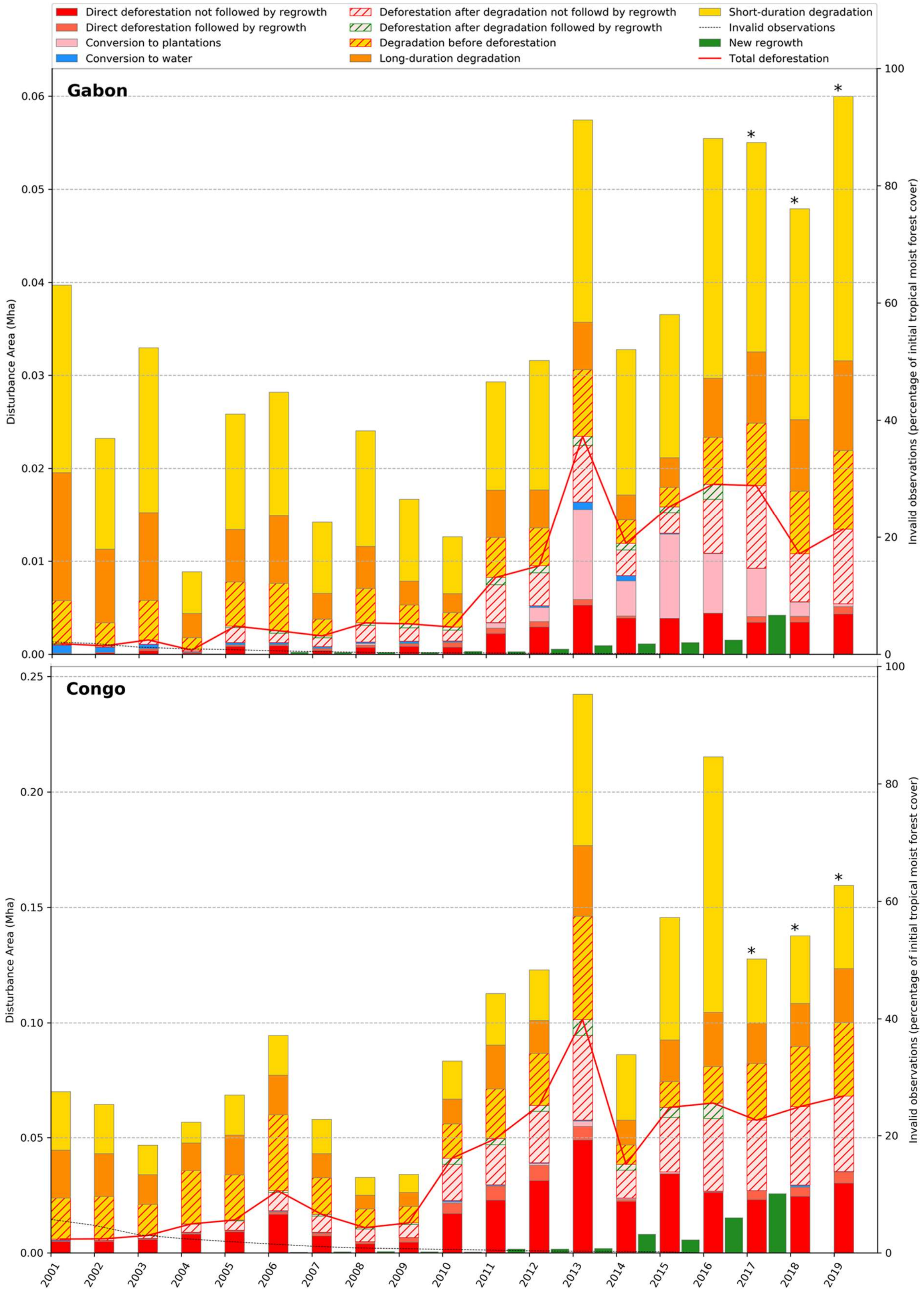
Direct deforestation not followed by regrowth
Direct deforestation followed by regrowth
Conversion to plantations
Conversion to water
Deforestation after degradation not followd by regrowth
Deforestation after degradation followed by regrowth
Degradation before deforestation
Long-duration degradation
Short-duration degradation
Invalid observations
New regrowth
Total deforestation
Gabon
Congo
Disturbance Area (Mha)
Invalid observations (percentage of initial tropical moist forest cover)
*


750

751 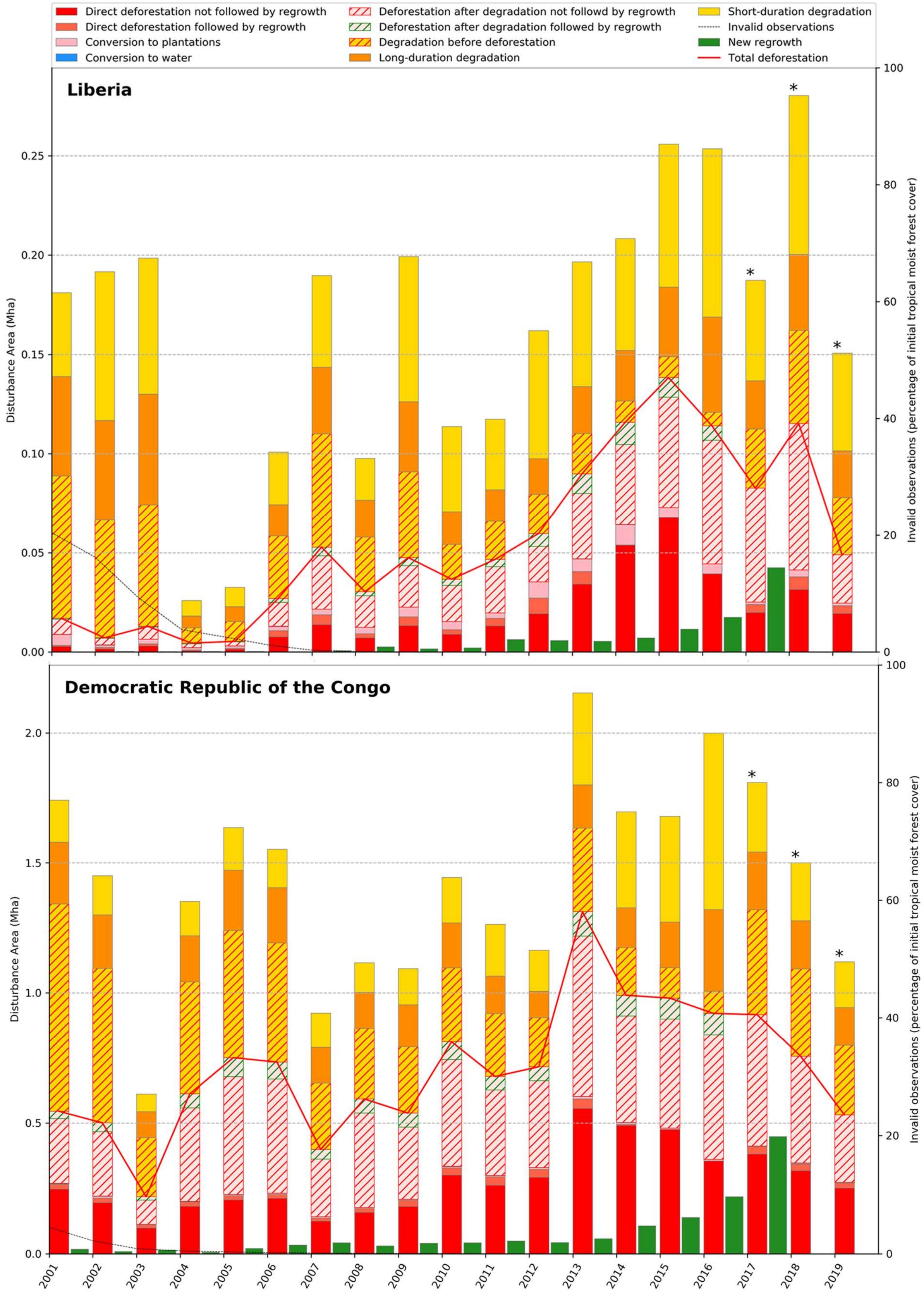
Direct deforestation not followed by regrowth
Direct deforestation followed by regrowth
Conversion to plantations
Conversion to water
Deforestation after degradation not followd by regrowth
Deforestation after degradation followed by regrowth
Degradation before deforestation
Long-duration degradation
Short-duration degradation
Invalid observations
New regrowth
Total deforestation
Liberia
Democratic Republic of the Congo
Disturbance Area (Mha)
Invalid observations (percentage of initial tropical moist forest cover)
2001
2002
2003
2004
2005
2006
2007
2008
2009
2010
2011
2012
2013
2014
2015
2016
2017
2018
2019

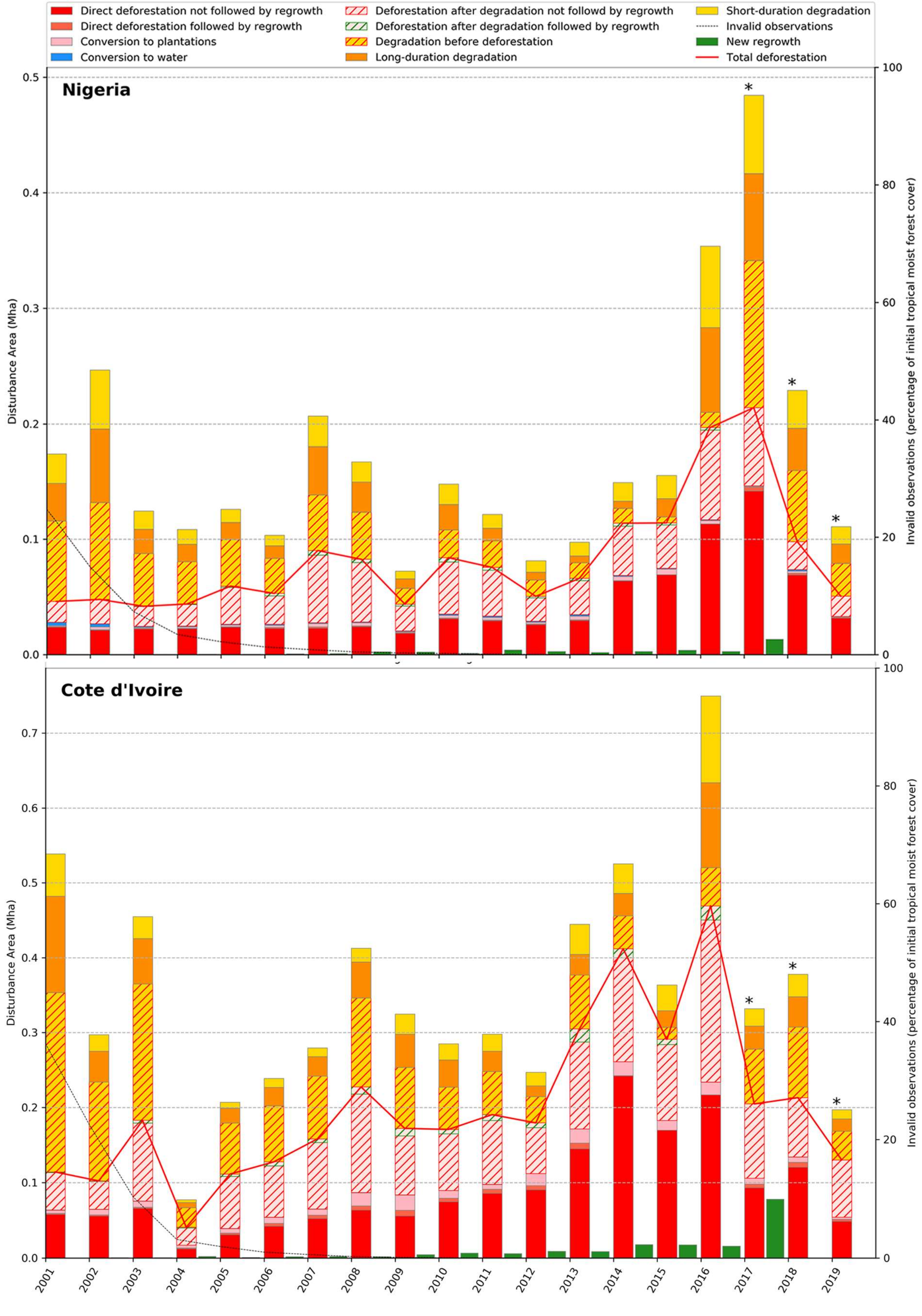
Direct deforestation not followed by regrowth
Direct deforestation followed by regrowth
Conversion to plantations
Conversion to water
Deforestation after degradation not followd by regrowth
Deforestation after degradation followed by regrowth
Degradation before deforestation
Long-duration degradation
Short-duration degradation
Invalid observations
New regrowth
Total deforestation
Nigeria
Cote d'Ivoire
Disturbance Area (Mha)
Invalid observations (percentage of initial tropical moist forest cover)

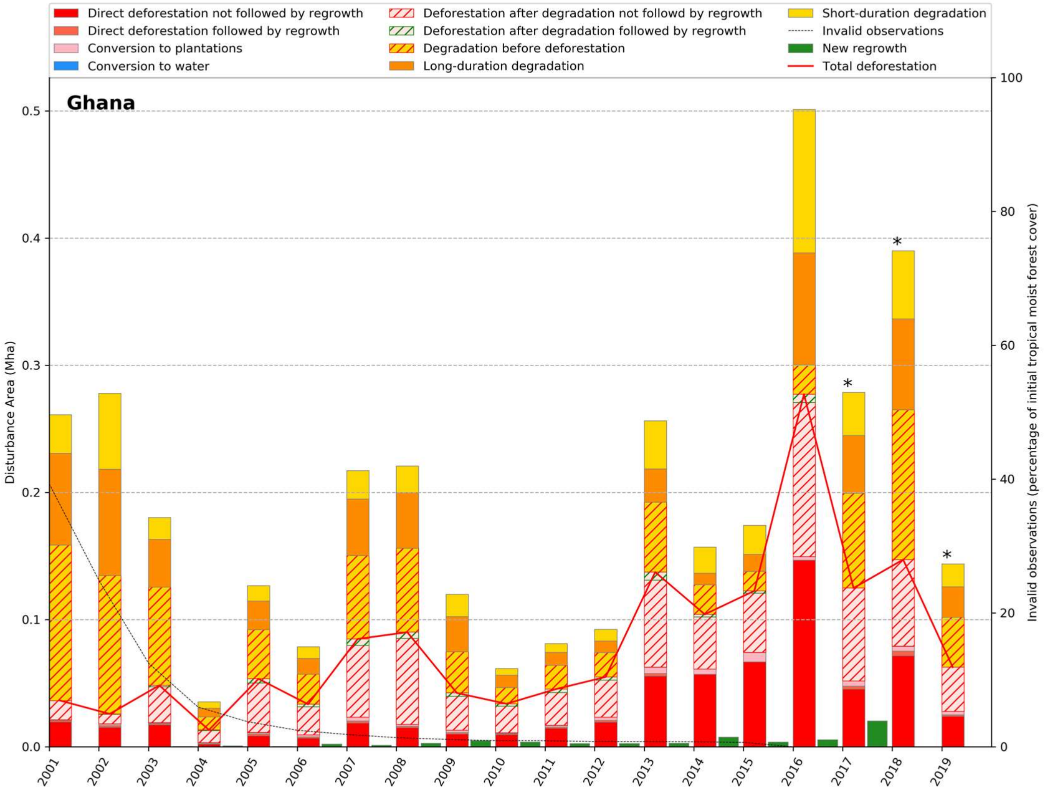


**Fig. S12** Forecasted year of full disappearance of undisturbed moist forest for countries with an undisturbed forest area larger than 1 million ha in January 2020, by applying the average disturbance rate for 2010-2019.

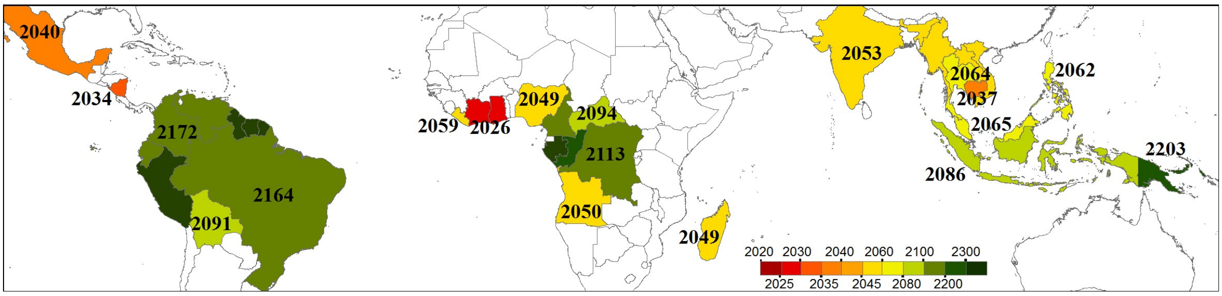

**Fig. S13** Extent of the study area. The study area is defined using the ecological zones adopted by the FAO and includes the following zones: 'Tropical rainforest', 'Tropical moist forest', 'Tropical mountain system' and 'Tropical dry forest'.

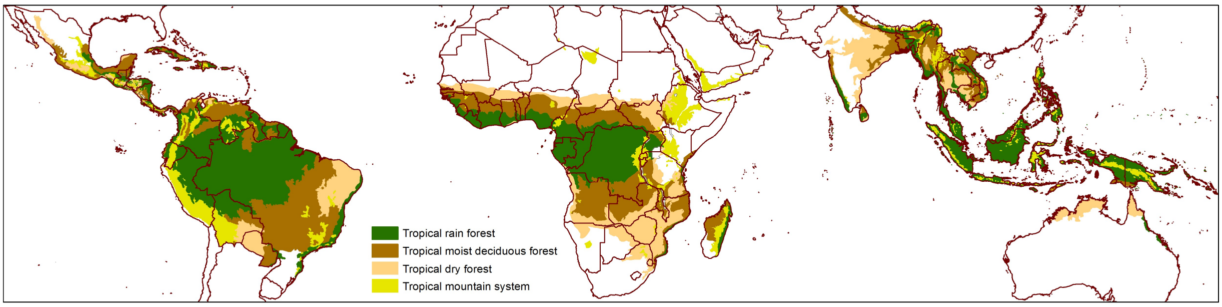


**Fig. S14** Total number of valid observations per pixel from the full Landsat archive (1982-2019) over the pan-tropical belt.

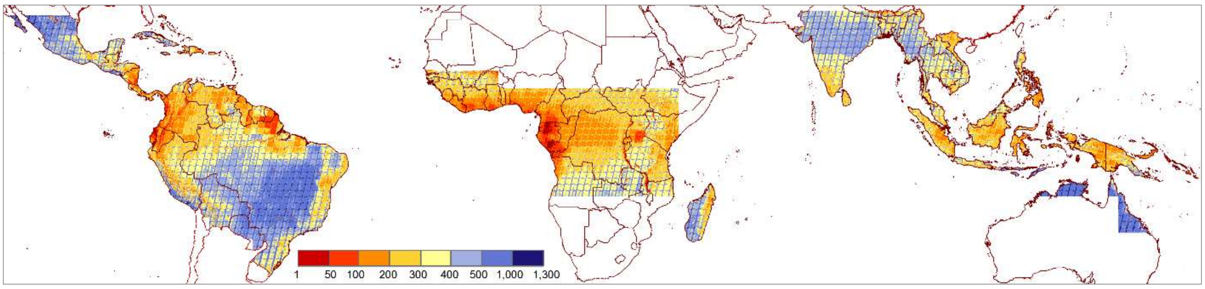


**Fig. S15** Year of first valid observation from the full Landsat archive (1982-2019), across the pan-tropical belt.

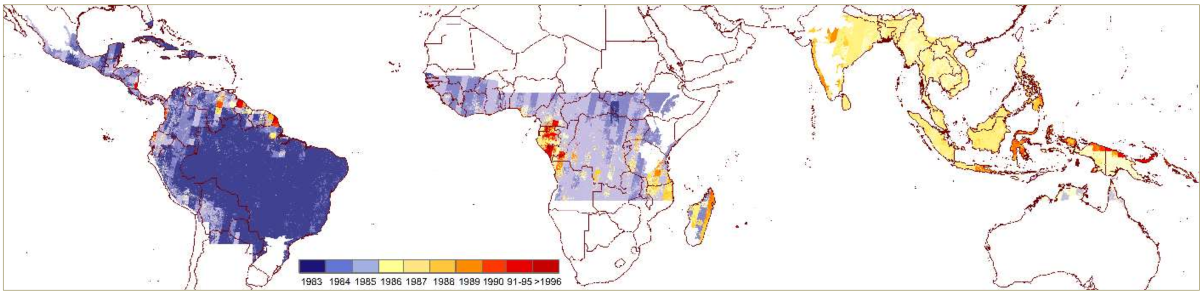

**Fig. S16** Annual average number of valid observations per pixel (by continent) over the period 1982-2019 Landsat archive for the tropical moist forest domain.

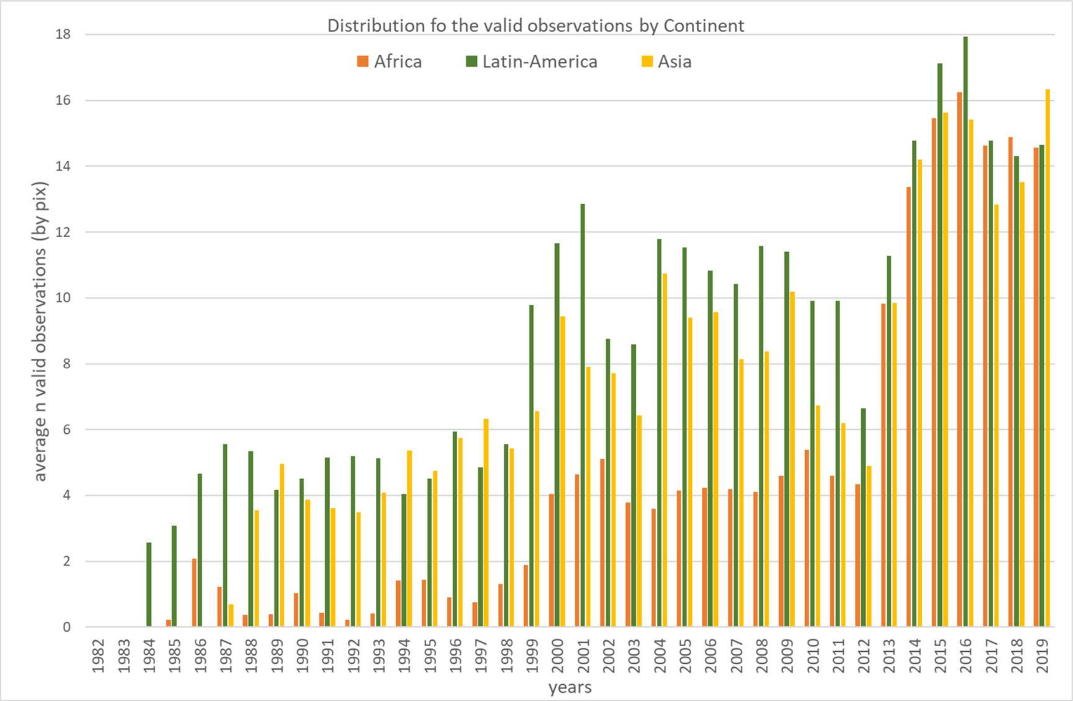

**Fig. S17** Multispectral feature space with following clusters: moist forest (dark green points), non-evergreen cover (orange for bare soils, brown for deciduous vegetation, light green for agriculture, blue for water) and invalid pixels (grey for shadows, purple for clouds and pink for haze): (a) hue versus saturation (both from SWIR2, NIR, red), (b) hue versus value (both from SWIR2, NIR, red), (c) hue (from SWIR2, NIR, red) versus TIR, (d) hue versus NDWI.

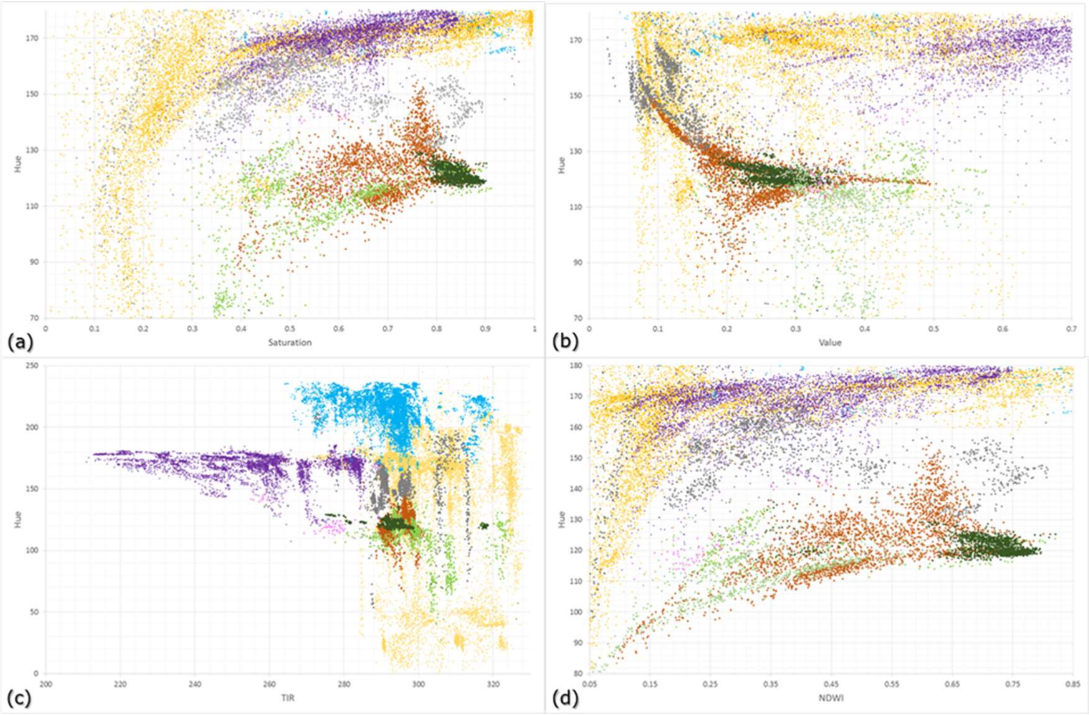

**Fig. S18** First year of the monitoring period used for changes analysis

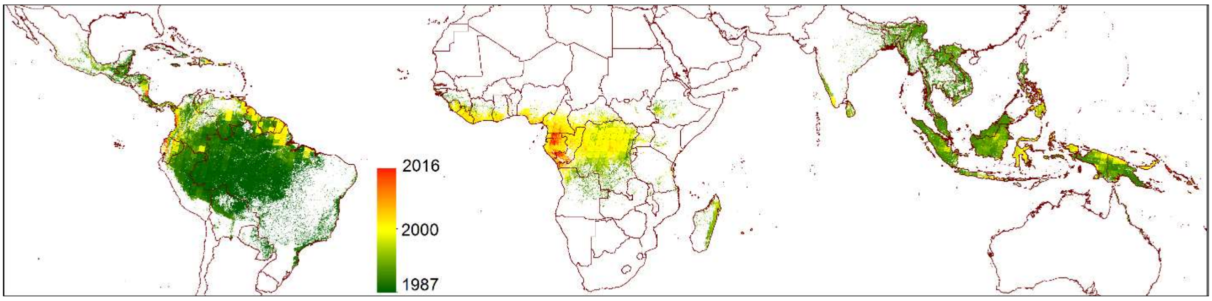


**Fig. S19** Distribution of the duration of disturbances recorded over the period 1990-2016 for each continent. The temporal thresholds used to define short-duration degradation, long-duration degradation and deforestation are represented as dashed lines (at 1 and 2.5 years, respectively).

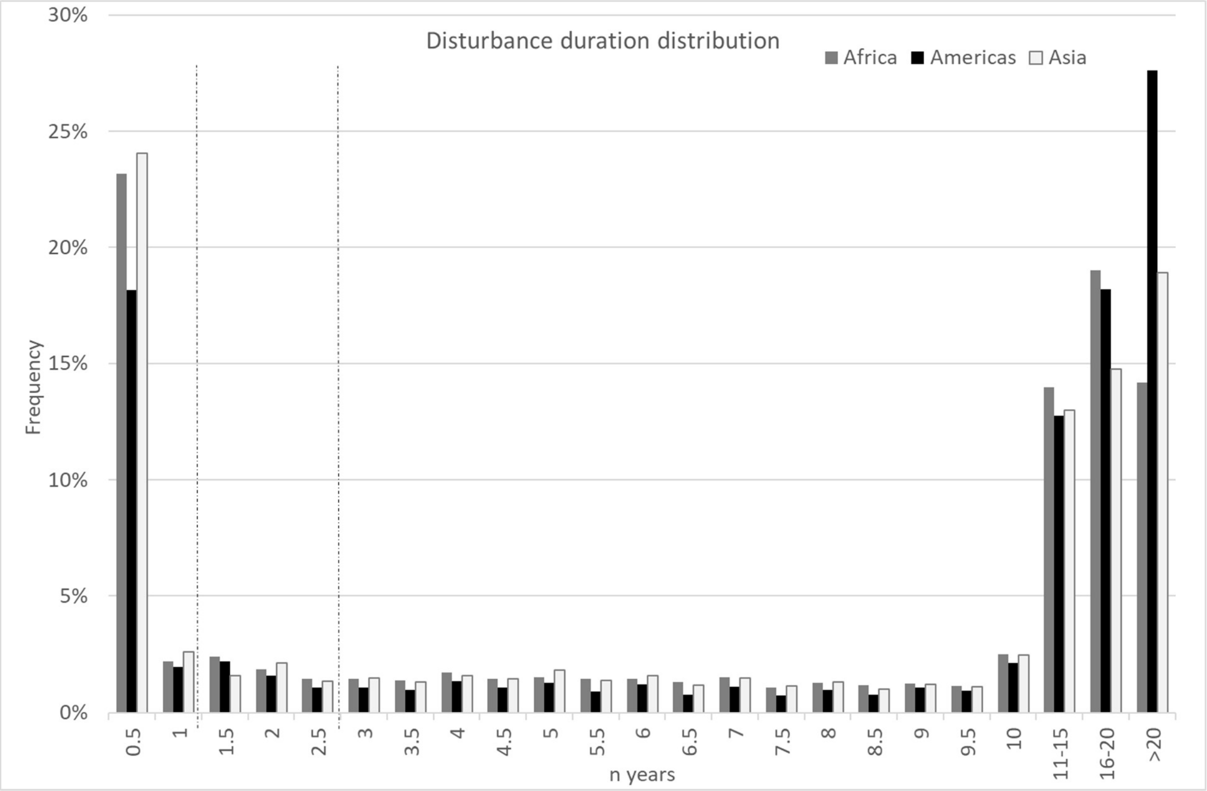

**Table S1.** Correspondence matrix between the main classes of our transition map and the GFC loss and gain products for the period 2001-2019 (areas in million ha) at the pan-tropical scale (A), and for the three continents (B-D).

| (A) Pan-tropical region | GFC | | | | | |
|---|---|---|---|---|---|---|
| Transition map | No loss, no gain | Gain, no loss | Loss, no gain | Loss & Gain | % agree | % disagree |
| *Undisturbed Jan 2020* | 958.3 | 0.4 | 5.6 | 0.2 | | |
| *Old Degradation or Regrowth (1982-2000)* | 52.4 | 1.1 | 2.9 | 0.3 | 97.2 | 2.8 |
| *Old Deforestation (1982-2000)* | 39.1 | 2.1 | 15.5 | 2.1 | | |
| Degradation 2001-2019 | 64.3 | 0.5 | 20.1 | 1.2 | 24.7 | 75.3 |
| Regrowth after deforestation (2001-2016) | 12.7 | 0.3 | 5.7 | 0.8 | 33.2 | 66.8 |
| Deforestation after degradation (2001-2019) | 35.3 | 0.3 | 22.8 | 1.4 | 40.6 | 59.4 |
| Direct deforestation (2001-2019) | 25.3 | 0.3 | 42.8 | 2.6 | 64.0 | 36.0 |
| Total without other LC | 1248.0 | 5.4 | 180.8 | 12.7 | | |
| % agreement | 83.8 | | 86.3 | | 84.2 | |
| % disagreement | 16.2 | | 13.7 | | | 15.8 |

| (B) Africa | GFC | | | | | |
|---|---|---|---|---|---|---|
| Transition map | No loss, no gain | Gain, no loss | Loss, no gain | Loss & Gain | % agree | % disagree |
| *Undisturbed Jan 2020* | 204.4 | 0.2 | 2.1 | 0.1 | | |
| *Old Degradation or Regrowth (1982-2000)* | 8.2 | 0.1 | 0.5 | 0.1 | 98.0 | 2.0 |
| *Old Deforestation (1982-2000)* | 4.4 | 0.0 | 1.3 | 0.1 | | |
| Degradation 2001-2019 | 15.4 | 0.2 | 6.1 | 0.3 | 29.2 | 70.8 |
| Regrowth after deforestation (2001-2016) | 1.7 | 0.0 | 0.8 | 0.1 | 33.4 | 66.6 |
| Deforestation after degradation (2001-2019) | 9.4 | 0.1 | 7.4 | 0.5 | 45.6 | 54.4 |
| Direct deforestation (2001-2019) | 8.4 | 0.0 | 6.2 | 0.1 | 43.1 | 56.9 |
| Total without other LC | 269.5 | 0.8 | 38.1 | 1.9 | | |
| % agreement | 80.3 | | 89.8 | | 81.5 | |
| % disagreement | 19.7 | | 10.2 | | | 18.5 |

| (C) Latin America | GFC | | | | | |
|---|---|---|---|---|---|---|
| Transition map | No loss, no gain | Gain, no loss | Loss, no gain | Loss & Gain | % agree | % disagree |
| *Undisturbed Jan 2020* | 563.0 | 0.0 | 1.4 | 0.0 | | |
| *Old Degradation or Regrowth (1982-2000)* | 23.0 | 0.3 | 0.9 | 0.1 | 97.8 | 2.2 |
| *Old Deforestation (1982-2000)* | 23.6 | 0.3 | 9.5 | 0.9 | | |
| Degradation 2001-2019 | 27.3 | 0.1 | 6.8 | 0.1 | 20.2 | 79.8 |
| Regrowth after deforestation (2001-2016) | 5.5 | 0.1 | 2.1 | 0.2 | 29.3 | 70.7 |
| Deforestation after degradation (2001-2019) | 13.4 | 0.0 | 8.8 | 0.3 | 40.3 | 59.7 |
| Direct deforestation (2001-2019) | 9.3 | 0.0 | 23.9 | 0.5 | 72.3 | 27.7 |
| Total without other LC | 687.7 | 0.9 | 86.0 | 2.8 | | |
| % agreement | 88.5 | | 85.7 | | 88.2 | |
| % disagreement | 11.5 | | 14.3 | | | 11.8 |

| (D) Asia - Oceania | GFC | | | | | |
|---|---|---|---|---|---|---|
| Transition map | No loss, no gain | Gain, no loss | Loss, no gain | Loss & Gain | % agree | % disagree |
| *Undisturbed Jan 2020* | 191.0 | 0.2 | 2.0 | 0.1 | | |
| *Old Degradation or Regrowth (1982-2000)* | 21.2 | 0.7 | 1.5 | 0.2 | 94.8 | 5.2 |
| *Old Deforestation (1982-2000)* | 11.2 | 1.7 | 4.7 | 1.1 | | |
| Degradation 2001-2019 | 21.6 | 0.2 | 7.2 | 0.7 | 26.6 | 73.4 |
| Regrowth after deforestation (2001-2016) | 5.5 | 0.2 | 2.8 | 0.5 | 36.5 | 63.5 |
| Deforestation after degradation (2001-2019) | 12.4 | 0.1 | 6.6 | 0.6 | 36.5 | 63.5 |
| Direct deforestation (2001-2019) | 7.7 | 0.2 | 12.7 | 2.0 | 65.2 | 34.8 |
| Total deforestation (2001-2019) | 20.1 | 0.3 | 19.3 | 2.7 | 51.8 | 48.2 |
| % agreement | 76.1 | | 85.1 | | 77.7 | |
| % disagreement | 23.9 | | 14.9 | | | 22.3 |

**Table S2.** Accuracy matrix for all the single-date interpretations (12 345 sample plots) by continent:

a) detailed matrix with all Landsat and HR classes;

| Reference | Landsat Mostly non-forest | | | | | | | | | | | | | | | | | | | | | | | |
|---|---|---|---|---|---|---|---|---|---|---|---|---|---|---|---|---|---|---|---|---|---|---|---|---|
| | *HR Mostly non-forest* | | | | *Minor non-forest* | | | | *Shurb* | | | | *Forest* | | | | *Invalid* | | | | *Total points User* | | | |
| User map | AFR | Asia | SAM | *Tot* | AFR | Asia | SAM | *Tot* | AFR | Asia | SAM | *Tot* | AFR | Asia | SAM | *Tot* | AFR | Asia | SAM | *Tot* | *AFR* | *Asia* | *SAM* | *Tot* |
| Mostly non-forest | 234 | 171 | 236 | *641* | 77 | 60 | 84 | *221* | 104 | 84 | 77 | *265* | 37 | 40 | 93 | *170* | 58 | 167 | 65 | *290* | *510* | *522* | *555* | *1587* |
| Minor non-forest | 53 | 64 | 58 | *175* | 36 | 16 | 16 | *68* | 23 | 21 | 11 | *55* | 9 | 9 | 17 | *35* | 27 | 64 | 21 | *112* | *148* | *174* | *123* | *445* |
| Forest | 0 | 29 | 14 | *43* | 2 | 4 | 1 | *7* | 3 | 6 | 5 | *14* | 3 | 1 | 1 | *5* | 2 | 7 | 0 | *9* | *10* | *47* | *21* | *78* |
| *Total points Producer* | *287* | *264* | *308* | *859* | *115* | *80* | *101* | *296* | *130* | *111* | *93* | *334* | *49* | *50* | *111* | *210* | *87* | *238* | *86* | *411* | *668* | *743* | *699* | *2110* |

| Reference | Landsat Minor non-forest | | | | | | | | | | | | | | | | | | | | | | | |
|---|---|---|---|---|---|---|---|---|---|---|---|---|---|---|---|---|---|---|---|---|---|---|---|---|
| | *HR Mostly non-forest* | | | | *Minor non-forest* | | | | *Shurb* | | | | *Forest* | | | | *Invalid* | | | | *Total points User* | | | |
| User map | AFR | Asia | SAM | *Tot* | AFR | Asia | SAM | *Tot* | AFR | Asia | SAM | *Tot* | AFR | Asia | SAM | *Tot* | AFR | Asia | SAM | *Tot* | *AFR* | *Asia* | *SAM* | *Tot* |
| Mostly non-forest | 139 | 94 | 70 | *303* | 171 | 95 | 200 | *466* | 149 | 84 | 129 | *362* | 40 | 34 | 69 | *143* | 58 | 130 | 42 | *230* | *557* | *437* | *510* | *1504* |
| Minor non-forest | 141 | 107 | 58 | *306* | 209 | 150 | 218 | *577* | 243 | 157 | 134 | *534* | 71 | 41 | 89 | *201* | 137 | 229 | 46 | *412* | *801* | *684* | *545* | *2030* |
| Forest | 23 | 39 | 18 | *80* | 56 | 61 | 76 | *193* | 34 | 28 | 8 | *70* | 9 | 8 | 5 | *22* | 9 | 120 | 5 | *134* | *131* | *256* | *112* | *499* |
| *Total points Producer* | *303* | *240* | *146* | *689* | *436* | *306* | *494* | *1236* | *426* | *269* | *271* | *966* | *120* | *83* | *163* | *366* | *204* | *479* | *93* | *776* | *1489* | *1377* | *1167* | *4033* |

| Reference | Landsat Forest | | | | | | | | | | | | | | | | | | | | | | | |
|---|---|---|---|---|---|---|---|---|---|---|---|---|---|---|---|---|---|---|---|---|---|---|---|---|
| | *HR Mostly non-forest* | | | | *Minor non-forest* | | | | *Shurb* | | | | *Forest* | | | | *Invalid* | | | | *Total points User* | | | |
| User map | AFR | Asia | SAM | *Tot* | AFR | Asia | SAM | *Tot* | AFR | Asia | SAM | *Tot* | AFR | Asia | SAM | *Tot* | AFR | Asia | SAM | *Tot* | *AFR* | *Asia* | *SAM* | *Tot* |
| Mostly non-forest | 4 | 0 | 1 | *5* | 7 | 5 | 14 | *26* | 9 | 9 | 50 | *68* | 1 | 5 | 17 | *23* | 0 | 19 | 2 | *21* | *21* | *38* | *84* | *143* |
| Minor non-forest | 5 | 3 | 6 | *14* | 20 | 20 | 33 | *73* | 11 | 28 | 64 | *103* | 6 | 26 | 47 | *79* | 2 | 46 | 20 | *68* | *44* | *123* | *170* | *337* |
| Forest | 62 | 86 | 63 | *211* | 219 | 224 | 244 | *687* | 244 | 499 | 169 | *912* | 824 | 434 | 1399 | *2657* | 252 | 691 | 310 | *1253* | *1601* | *1934* | *2185* | *5720* |
| *Total points Producer* | *71* | *89* | *70* | *230* | *246* | *249* | *291* | *786* | *264* | *536* | *283* | *1083* | *831* | *465* | *1463* | *2759* | *254* | *756* | *332* | *1342* | *1666* | *2095* | *2439* | *6200* |

b) simplified matrix with non-forest and forest classes.

| Reference | Non-forest | | | | Forest | | | | |
|---|---|---|---|---|---|---|---|---|---|
| User map | AFR | ASIA | Latin-Am | Tot | AFR | ASIA | Latin-Am | Tot | Tot User |
| Non-forest | 2016 | 1817 | 1733 | 5566 | 65 | 161 | 254 | 480 | 6046 |
| Forest | 141 | 303 | 133 | 577 | 1601 | 1934 | 2185 | 5720 | 6297 |
| *Total Producer* | 2157 | 2120 | 1866 | 6143 | 1666 | 2095 | 2439 | 6200 | 12343 |

**Table S3.** Accuracy results by continent and Landsat sensor.

| | Mostly non-forest | | | | Forest | | | |
|---|---|---|---|---|---|---|---|---|
| Continent | AFR | Asia | Latin-Am | Tot | AFR | Asia | SAM | Tot |
| % Prod Accuracy | 93.5 | 85.7 | 92.9 | **90.6** | 96.1 | 92.3 | 89.6 | **92.3** |
| % Ommission error | 6.5 | 14.3 | 7.1 | **9.4** | 3.9 | 7.7 | 10.4 | **7.7** |
| % User Accuracy | 96.9 | 91.9 | 87.2 | **92.1** | 91.9 | 86.5 | 94.3 | **90.8** |
| % Commission error | 3.1 | 8.1 | 12.8 | **7.9** | 8.1 | 13.5 | 5.7 | **9.2** |
| % Overall Accuracy | **94.6** | **89.0** | **91.0** | **91.4** | | | | |

| Sensor | LC8 | LE7 | LT5 | Tot | LC8 | LE7 | LT5 | Tot |
|---|---|---|---|---|---|---|---|---|
| % Prod Accuracy | 90.3 | 90.6 | 90.2 | **90.4** | 86.8 | 92.8 | 94.8 | **92.3** |
| % Ommission error | 9.7 | 9.4 | 9.8 | **9.6** | 13.2 | 7.2 | 5.2 | **7.7** |
| % User Accuracy | 89.1 | 92.2 | 93.9 | **91.9** | 88.2 | 91.3 | 91.7 | **90.8** |
| % Commission error | 10.9 | 7.8 | 6.1 | **8.1** | 11.8 | 8.7 | 8.3 | **9.2** |
| % Overall Accuracy | **88.7** | **91.8** | **92.7** | **91.4** | | | | |

**Table S4.** Area-weighted confusion matrix for the transition map and a validation reference dataset of 4 139 sample plots (%).

| | Reference | *Forest on Landsat & HR* | *Forest on Landsat & non-forest on HR* | At least 1 minor disruption on Landsat | | | | At least 1 major disruption on Landsat | | | | *total* |
|---|---|---|---|---|---|---|---|---|---|---|---|---|
| Transition map | *Max N disruptions* | *0* | *0* | *1* | *2* | *3* | *4* | *1* | *2* | *3* | *4* | |
| Undisturbed (0pix) | 0 | 48.0 | 0.6 | 2.6 | 0.2 | 0.0 | 0.0 | 0.2 | 0.0 | 0.0 | 0.0 | *52* |
| Mostly Undisturbed (1-4pix disturbed) | *1* | 1.5 | 0.1 | 0.6 | 0.1 | 0.0 | 0.0 | 0.1 | 0.0 | 0.0 | 0.0 | *2* |
| | *2-3* | 1.2 | 0.2 | 0.4 | 0.1 | 0.0 | 0.0 | 0.0 | 0.0 | 0.0 | 0.0 | *2* |
| | *>3* | 0.2 | 0.1 | 0.7 | 0.4 | 0.2 | 0.1 | 0.4 | 1.0 | 6.6 | 2.3 | *12* |
| Mostly changed (5-9pix disturbed) | *1* | 0.4 | 0.1 | 0.2 | 0.0 | 0.0 | 0.0 | 0.0 | 0.0 | 0.0 | 0.0 | *1* |
| | *2-3* | 0.6 | 0.1 | 0.6 | 0.2 | 0.0 | 0.0 | 0.2 | 0.3 | 0.0 | 0.0 | *2* |
| | *>3* | 0.2 | 0.1 | 2.2 | 1.4 | 1.1 | 0.0 | 3.3 | 9.4 | 11.2 | 0.4 | *29* |
| *Total* | | *52* | *1* | *7* | *2* | *1* | *0* | *4* | *11* | *18* | *3* | *100* |

**Table S5.** Area-weighted matrix showing the transition map versus the reference dataset and error estimation (million ha).

| | *Reference* | | |
|---|---|---|---|
| *Transition map* | Undisturbed | Forest Change | Area on the map |
| Undisturbed | 898.64 | 65.76 | 964.40 |
| Forest Change | 27.38 | 297.82 | 325.20 |
| Area cor (Mha) | 926.02 | 363.58 | 1289.60 |
| Producer Accuracy | *97.0%* | *81.9%* | |
| User Accuracy | *93.2%* | *91.6%* | |
| Commission error (Mha) | 65.8 | 27.4 | |
| Omission error (Mha) | 27.4 | 65.8 | |
| Difference (Mha) | -38.4 | 38.4 | |
| CI (95%) | 14.8 | 14.8 | |

**Table S6** Annual rate of loss of undisturbed TMF between 1990 and 2019 of 5 years, 30 year (1990-2019), and 10 years (1990-1999**,** 2000-2009, 2010-2019) by country and percentage of annual area loss versus initial TMF areas over the period 1990-2019 (A). Annual rate of direct deforestation, deforestation after degradation and degradation (followed or not by deforestation) and percentage of each disturbance's type over the total disturbances during the 30-year period, and areas of water conversion and plantations and percentage (over the total disturbances) (B).

**A.**

| Country | Undisturbed | | Annual decline of Undisturbed TMF (Mha) | | | | | | | | | | Decline |
|---|---|---|---|---|---|---|---|---|---|---|---|---|---|
| | 1990 | 2019 | [1990-1995[ | [1995-2000[ | [2000-2005[ | [2005-2010[ | [2010-2015[ | [2015-2020[ | [1990-2020[ | [1990-2000[ | [2000-2010[ | [2010-2020[ | 30y (%) |
| Brazil | 411.20 | 308.89 | 2.65 | 4.94 | 4.26 | 3.14 | 2.14 | 3.34 | 3.41 | 3.79 | 3.70 | 2.74 | 24.9% |
| Indonesia | 162.25 | 94.02 | 1.09 | 3.84 | 2.57 | 2.31 | 1.86 | 1.96 | 2.27 | 2.47 | 2.44 | 1.91 | 42.1% |
| DRC | 141.42 | 105.80 | 0.20 | 1.12 | 1.38 | 1.26 | 1.54 | 1.62 | 1.19 | 0.66 | 1.32 | 1.58 | 25.2% |
| Peru | 75.60 | 66.64 | 0.14 | 0.41 | 0.32 | 0.33 | 0.28 | 0.31 | 0.30 | 0.28 | 0.32 | 0.29 | 11.8% |
| Colombia | 74.20 | 58.14 | 0.14 | 0.65 | 0.85 | 0.51 | 0.51 | 0.55 | 0.54 | 0.40 | 0.68 | 0.53 | 21.6% |
| Venezuela | 47.26 | 38.49 | 0.06 | 0.30 | 0.56 | 0.22 | 0.26 | 0.35 | 0.29 | 0.18 | 0.39 | 0.31 | 18.6% |
| PNG | 42.08 | 34.61 | 0.15 | 0.43 | 0.29 | 0.13 | 0.16 | 0.33 | 0.25 | 0.29 | 0.21 | 0.24 | 17.7% |
| Bolivia | 36.80 | 24.28 | 0.19 | 0.56 | 0.46 | 0.38 | 0.57 | 0.35 | 0.42 | 0.37 | 0.42 | 0.46 | 34.0% |
| Malaysia | 29.85 | 15.20 | 0.31 | 0.74 | 0.55 | 0.55 | 0.45 | 0.33 | 0.49 | 0.52 | 0.55 | 0.39 | 49.1% |
| Congo | 24.37 | 22.21 | 0.01 | 0.02 | 0.06 | 0.06 | 0.13 | 0.16 | 0.07 | 0.01 | 0.06 | 0.14 | 8.9% |
| Myanmar | 24.15 | 9.57 | 0.28 | 0.78 | 0.59 | 0.51 | 0.39 | 0.36 | 0.49 | 0.53 | 0.55 | 0.38 | 60.4% |
| Gabon | 24.15 | 23.45 | 0.00 | 0.01 | 0.03 | 0.02 | 0.03 | 0.05 | 0.02 | 0.00 | 0.02 | 0.04 | 2.9% |
| Cameroon | 22.71 | 19.82 | 0.00 | 0.03 | 0.13 | 0.08 | 0.12 | 0.21 | 0.10 | 0.02 | 0.11 | 0.16 | 12.7% |
| Guyana | 18.87 | 17.95 | 0.01 | 0.03 | 0.04 | 0.03 | 0.03 | 0.05 | 0.03 | 0.02 | 0.03 | 0.04 | 4.9% |
| Philippines | 16.78 | 8.52 | 0.14 | 0.38 | 0.36 | 0.24 | 0.24 | 0.29 | 0.28 | 0.26 | 0.30 | 0.26 | 49.2% |
| Ecuador | 15.83 | 12.12 | 0.01 | 0.12 | 0.21 | 0.12 | 0.12 | 0.17 | 0.12 | 0.06 | 0.16 | 0.14 | 23.4% |
| Lao PDR | 15.70 | 5.48 | 0.24 | 0.48 | 0.42 | 0.32 | 0.26 | 0.33 | 0.34 | 0.36 | 0.37 | 0.29 | 65.1% |
| Viet Nam | 14.93 | 4.81 | 0.27 | 0.52 | 0.41 | 0.37 | 0.21 | 0.25 | 0.34 | 0.39 | 0.39 | 0.23 | 67.8% |
| Suriname | 13.76 | 13.15 | 0.01 | 0.02 | 0.02 | 0.02 | 0.02 | 0.03 | 0.02 | 0.01 | 0.02 | 0.03 | 4.5% |
| India | 11.62 | 4.20 | 0.17 | 0.38 | 0.33 | 0.26 | 0.18 | 0.17 | 0.25 | 0.27 | 0.29 | 0.18 | 63.9% |
| Mexico | 11.60 | 3.05 | 0.18 | 0.60 | 0.29 | 0.25 | 0.15 | 0.24 | 0.28 | 0.39 | 0.27 | 0.20 | 73.7% |
| Madagascar | 10.45 | 3.41 | 0.08 | 0.48 | 0.30 | 0.19 | 0.21 | 0.14 | 0.23 | 0.28 | 0.25 | 0.18 | 67.3% |
| CAR | 9.91 | 7.12 | 0.01 | 0.07 | 0.15 | 0.08 | 0.08 | 0.16 | 0.09 | 0.04 | 0.12 | 0.12 | 28.1% |
| Cote d'Ivoire | 9.74 | 1.80 | 0.01 | 0.16 | 0.36 | 0.29 | 0.36 | 0.40 | 0.26 | 0.08 | 0.33 | 0.38 | 81.5% |
| Angola | 9.52 | 3.13 | 0.12 | 0.38 | 0.30 | 0.16 | 0.17 | 0.15 | 0.21 | 0.25 | 0.23 | 0.16 | 67.1% |
| Liberia | 9.33 | 5.98 | 0.01 | 0.02 | 0.12 | 0.12 | 0.16 | 0.23 | 0.11 | 0.02 | 0.12 | 0.19 | 35.9% |
| Thailand | 9.06 | 4.07 | 0.17 | 0.24 | 0.17 | 0.16 | 0.10 | 0.16 | 0.17 | 0.21 | 0.17 | 0.13 | 55.1% |
| Nigeria | 8.44 | 4.50 | 0.02 | 0.06 | 0.19 | 0.14 | 0.12 | 0.27 | 0.13 | 0.04 | 0.16 | 0.19 | 46.7% |
| French Guiana | 8.14 | 7.96 | 0.00 | 0.00 | 0.01 | 0.01 | 0.01 | 0.01 | 0.01 | 0.00 | 0.01 | 0.01 | 2.1% |
| Nicaragua | 6.13 | 2.10 | 0.02 | 0.13 | 0.16 | 0.14 | 0.18 | 0.18 | 0.13 | 0.08 | 0.15 | 0.18 | 65.8% |
| Cambodia | 5.85 | 2.28 | 0.05 | 0.12 | 0.11 | 0.13 | 0.14 | 0.15 | 0.12 | 0.09 | 0.12 | 0.15 | 61.1% |
| Ghana | 5.84 | 1.71 | 0.00 | 0.05 | 0.19 | 0.15 | 0.13 | 0.30 | 0.14 | 0.03 | 0.17 | 0.21 | 70.8% |

B.

| Country | Annual rate for period [1990-2020[ (Mha) | | | % over the total disturbances | | | Conversion to Plantations | | Conversion to water | |
|---|---|---|---|---|---|---|---|---|---|---|
| | Direct defor | Defor. after degrad | Degrad | Direct defor | Defor. after degrad | Degrad | Mha | % | Mha | % |
| Brazil | 1.63 | 0.56 | 0.58 | 59% | 20% | 21% | 1.623 | 7.4% | 0.619 | 21.0% |
| Indonesia | 0.68 | 0.44 | 0.62 | 39% | 25% | 36% | 12.660 | 57.4% | 0.515 | 17.5% |
| DRC | 0.24 | 0.28 | 0.32 | 28% | 33% | 38% | 0.082 | 0.4% | 0.067 | 2.3% |
| Peru | 0.05 | 0.07 | 0.10 | 23% | 31% | 46% | 0.041 | 0.2% | 0.141 | 4.8% |
| Colombia | 0.12 | 0.12 | 0.14 | 31% | 32% | 38% | 0.081 | 0.4% | 0.102 | 3.5% |
| Venezuela | 0.00 | 0.00 | 0.01 | 2% | 1% | 96% | 0.071 | 0.3% | 0.103 | 3.5% |
| PNG | 0.02 | 0.04 | 0.13 | 9% | 22% | 69% | 0.006 | 0.0% | 0.131 | 4.5% |
| Bolivia | 0.11 | 0.09 | 0.12 | 34% | 27% | 39% | 0.000 | 0.0% | 0.111 | 3.8% |
| Malaysia | 0.22 | 0.05 | 0.14 | 53% | 13% | 33% | 5.257 | 23.8% | 0.131 | 4.4% |
| Congo | 0.01 | 0.01 | 0.03 | 24% | 19% | 57% | 0.006 | 0.0% | 0.013 | 0.4% |
| Myanmar | 0.11 | 0.12 | 0.10 | 33% | 36% | 31% | 0.020 | 0.1% | 0.113 | 3.8% |
| Gabon | 0.00 | 0.00 | 0.01 | 15% | 11% | 74% | 0.036 | 0.2% | 0.014 | 0.5% |
| Cameroon | 0.02 | 0.01 | 0.04 | 30% | 15% | 54% | 0.070 | 0.3% | 0.029 | 1.0% |
| Philippines | 0.02 | 0.07 | 0.11 | 12% | 33% | 55% | 0.003 | 0.0% | 0.042 | 1.4% |
| Ecuador | 0.02 | 0.02 | 0.05 | 18% | 26% | 56% | 0.000 | 0.0% | 0.031 | 1.0% |
| Lao PDR | 0.06 | 0.09 | 0.07 | 28% | 40% | 32% | 0.000 | 0.0% | 0.067 | 2.3% |
| Viet Nam | 0.07 | 0.07 | 0.08 | 31% | 32% | 37% | 0.003 | 0.0% | 0.175 | 6.0% |
| Suriname | 0.00 | 0.00 | 0.01 | 29% | 18% | 53% | 0.000 | 0.0% | 0.007 | 0.2% |
| India | 0.06 | 0.06 | 0.06 | 32% | 35% | 33% | 0.334 | 1.5% | 0.082 | 2.8% |
| Mexico | 0.06 | 0.07 | 0.06 | 33% | 37% | 31% | 0.000 | 0.0% | 0.029 | 1.0% |
| Madagascar | 0.06 | 0.06 | 0.03 | 40% | 39% | 21% | 0.061 | 0.3% | 0.025 | 0.9% |
| Cote d'Ivoire | 0.07 | 0.06 | 0.05 | 39% | 33% | 27% | 0.000 | 0.0% | 0.017 | 0.6% |
| CAR | 0.02 | 0.02 | 0.03 | 29% | 25% | 46% | 0.171 | 0.8% | 0.003 | 0.1% |
| Angola | 0.03 | 0.06 | 0.06 | 18% | 41% | 41% | 0.033 | 0.1% | 0.013 | 0.5% |
| Liberia | 0.02 | 0.02 | 0.05 | 18% | 21% | 61% | 0.146 | 0.7% | 0.003 | 0.1% |
| Thailand | 0.04 | 0.04 | 0.04 | 30% | 34% | 36% | 0.001 | 0.0% | 0.046 | 1.6% |
| Nigeria | 0.03 | 0.02 | 0.04 | 32% | 25% | 43% | 0.000 | 0.0% | 0.041 | 1.4% |
| French Guiana | 0.00 | 0.00 | 0.00 | 29% | 15% | 56% | 0.010 | 0.0% | 0.005 | 0.2% |
| Nicaragua | 0.03 | 0.03 | 0.03 | 28% | 39% | 33% | 0.000 | 0.0% | 0.005 | 0.2% |
| Cambodia | 0.07 | 0.02 | 0.01 | 69% | 19% | 13% | 0.000 | 0.0% | 0.065 | 2.2% |
| Ghana | 0.02 | 0.03 | 0.04 | 26% | 30% | 45% | 0.000 | 0.0% | 0.007 | 0.2% |

**Table S7** Correspondence between countries, subregions and continents.

| Africa | | Americas | | Asia-Oceania | |
|---|---|---|---|---|---|
| **Country Name** | **Region Name** | **Country Name** | **Region Name** | **Country Name** | **Region Name** |
| Angola | Central Africa | Anguilla | Central America | British Indian Ocean Territory | Insular Asia |
| Burundi | Central Africa | Antigua and Barbuda | Central America | Brunei Darussalam | Insular Asia |
| Cameroon | Central Africa | Aruba | Central America | Cocos (Keeling) Islands | Insular Asia |
| Central African Republic | Central Africa | Bahamas | Central America | Cook Islands | Insular Asia |
| Congo | Central Africa | Baker Island | Central America | French Polynesia | Insular Asia |
| Democratic Republic of the Congo | Central Africa | Belize | Central America | Indonesia | Insular Asia |
| Equatorial Guinea | Central Africa | British Virgin Islands | Central America | Malaysia | Insular Asia |
| Gabon | Central Africa | Cayman Islands | Central America | Maldives | Insular Asia |
| Rwanda | Central Africa | Clipperton Island | Central America | Micronesia (Federated States of) | Insular Asia |
| Sao Tome and Principe | Central Africa | Costa Rica | Central America | Palau | Insular Asia |
| Uganda | Central Africa | Dominica | Central America | Philippines | Insular Asia |
| Comoros | South and East Africa | Dominican Republic | Central America | Singapore | Insular Asia |
| Eritrea | South and East Africa | El Salvador | Central America | Tokelau | Insular Asia |
| Ethiopia | South and East Africa | Grenada | Central America | Australia | Insular Asia |
| Glorioso Island | South and East Africa | Guadeloupe | Central America | Guam | Insular Asia |
| Ilemi triangle | South and East Africa | Haiti | Central America | Kiribati | Insular Asia |
| Kenya | South and East Africa | Honduras | Central America | Nauru | Insular Asia |
| Madagascar | South and East Africa | Jamaica | Central America | Niue | Insular Asia |
| Malawi | South and East Africa | Jarvis Island | Central America | Samoa | Insular Asia |
| Mauritius | South and East Africa | Johnston Atoll | Central America | Tonga | Insular Asia |
| Mayotte | South and East Africa | Martinique | Central America | Tuvalu | Insular Asia |
| Mozambique | South and East Africa | Mexico | Central America | Arunachal Pradesh | South-East Asia |
| Reunion | South and East Africa | Midway Island | Central America | Azerbaijan | South-East Asia |
| Seychelles | South and East Africa | Netherlands Antilles | Central America | Bhutan | South-East Asia |
| South Sudan | South and East Africa | Palmyra Atoll | Central America | China | South-East Asia |
| United Republic of Tanzania | South and East Africa | Panama | Central America | Georgia | South-East Asia |
| Zambia | South and East Africa | Puerto Rico | Central America | India | South-East Asia |
| Benin | West Africa | Saint Vincent and the Grenadines | Central America | Iran (Islamic Republic of) | South-East Asia |
| Burkina Faso | West Africa | Turks and Caicos islands | Central America | Japan | South-East Asia |
| Cote d'Ivoire | West Africa | United States Virgin Islands | Central America | Lao People's Democratic Republic | South-East Asia |
| Gambia | West Africa | Bolivia | South America | Macau | South-East Asia |
| Ghana | West Africa | Brazil | South America | Marshall Islands | South-East Asia |
| Guinea | West Africa | Colombia | South America | Nepal | South-East Asia |
| Guinea-Bissau | West Africa | Ecuador | South America | Northern Mariana Islands | South-East Asia |
| Liberia | West Africa | French Guiana | South America | Pakistan | South-East Asia |
| Libya | West Africa | Guyana | South America | Paracel Islands | South-East Asia |
| Mali | West Africa | Paraguay | South America | Scarborough Reef | South-East Asia |
| Nigeria | West Africa | Pitcairn | South America | Senkaku Islands | South-East Asia |
| Senegal | West Africa | Saint Helena | South America | Thailand | South-East Asia |
| Sierra Leone | West Africa | Uruguay | South America | Viet Nam | South-East Asia |
| Togo | West Africa | | | | |

**Table S8.** Projections of forest cover in January 2050 (in million ha and percentage of forest cover in 2019) and year of decline considering the mean and confidence interval (minimum and maximum), for the main countries (with forest area greater than 1 million ha in 2020), considering the undisturbed forest (A) and the whole forest (undisturbed and degraded) (B).

**A.** Projections of the undisturbed forest

| Country | Observed forest area in 2020 | Predicted forest area in 2050 | Predicted relative forest decline in 2050 | *Year of decline mean* | *Year of decline min* | *Year of decline max* |
|---|---|---|---|---|---|---|
| Brazil | 308.9 | 243.2 | 21% | 2164 | 2126 | 2215 |
| DRC | 105.8 | 71.4 | 33% | 2113 | 2100 | 2129 |
| Indonesia | 94.0 | 51.3 | 45% | 2086 | 2069 | 2110 |
| Peru | 66.6 | 59.7 | 10% | 2316 | 2282 | 2353 |
| Colombia | 58.1 | 46.4 | 20% | 2172 | 2148 | 2200 |
| Venezuela | 38.5 | 31.4 | 18% | 2190 | 2123 | 2298 |
| PNG | 34.6 | 28.5 | 18% | 2195 | 2132 | 2292 |
| Bolivia | 24.3 | 13.9 | 43% | 2091 | 2064 | 2134 |
| Gabon | 23.5 | 22.3 | 5% | 2643 | 2454 | 2913 |
| Congo | 22.2 | 18.6 | 16% | 2211 | 2167 | 2268 |
| Cameroon | 19.8 | 15.4 | 22% | 2158 | 2113 | 2223 |
| Guyana | 18.0 | 16.9 | 6% | 2556 | 2405 | 2765 |
| Malaysia | 15.2 | 5.1 | 67% | 2065 | 2056 | 2076 |
| Suriname | 13.1 | 12.4 | 6% | 2541 | 2406 | 2723 |
| Ecuador | 12.1 | 8.7 | 28% | 2127 | 2091 | 2181 |
| Myanmar | 9.6 | 1.5 | 84% | 2055 | 2045 | 2069 |
| Philippines | 8.5 | 2.4 | 71% | 2062 | 2043 | 2095 |
| French Guiana | 8.0 | 7.8 | 2% | 3323 | 2996 | 3760 |
| CAR | 7.1 | 4.2 | 41% | 2094 | 2067 | 2135 |
| Liberia | 6.0 | 1.4 | 76% | 2059 | 2052 | 2068 |
| Lao PDR | 5.5 | 0.0 | 100% | 2046 | 2040 | 2056 |
| Viet Nam | 4.8 | 0.1 | 97% | 2050 | 2043 | 2061 |
| Nigeria | 4.5 | 0.0 | 100% | 2049 | 2037 | 2068 |
| India | 4.2 | 0.4 | 90% | 2053 | 2045 | 2064 |
| Thailand | 4.1 | 1.3 | 67% | 2064 | 2044 | 2103 |
| Madagascar | 3.4 | 0.0 | 100% | 2049 | 2042 | 2059 |
| Angola | 3.1 | 0.1 | 97% | 2050 | 2040 | 2066 |
| Mexico | 3.1 | 0.0 | 100% | 2040 | 2032 | 2053 |
| Cambodia | 2.3 | 0.0 | 100% | 2037 | 2031 | 2046 |
| Nicaragua | 2.1 | 0.0 | 100% | 2034 | 2028 | 2044 |
| Cote d'Ivoire | 1.8 | 0.0 | 100% | 2026 | 2024 | 2028 |
| Ghana | 1.7 | 0.0 | 100% | 2029 | 2025 | 2038 |

### **B.** Projections of the whole forest

| Country | Observed forest area in 2020 | Predicted forest area in 2050 | Predicted relative forest decline in 2050 | *Year of decline mean* | *Year of decline min* | *Year of decline max* |
|---|---|---|---|---|---|---|
| Brazil | 329.8 | 280.8 | 15% | 2221 | 2188 | 2262 |
| DRC | 116.9 | 91.0 | 22% | 2155 | 2132 | 2182 |
| Indonesia | 113.2 | 78.9 | 30% | 2118 | 2090 | 2157 |
| Peru | 70.4 | 66.3 | 6% | 2540 | 2475 | 2614 |
| Colombia | 63.8 | 55.1 | 14% | 2240 | 2210 | 2274 |
| Venezuela | 41.1 | 36.2 | 12% | 2272 | 2191 | 2391 |
| PNG | 38.9 | 36.9 | 5% | 2615 | 2429 | 2885 |
| Bolivia | 28.5 | 22.3 | 22% | 2157 | 2117 | 2214 |
| Gabon | 23.9 | 23.5 | 2% | 3740 | 3109 | 4738 |
| Congo | 23.3 | 21.4 | 8% | 2398 | 2329 | 2483 |
| Cameroon | 21.5 | 19.3 | 10% | 2313 | 2242 | 2406 |
| Malaysia | 19.6 | 12.5 | 36% | 2102 | 2078 | 2136 |
| Guyana | 18.4 | 18.0 | 2% | 3243 | 2968 | 3598 |
| Ecuador | 13.9 | 12.1 | 13% | 2252 | 2183 | 2350 |
| Suriname | 13.5 | 13.1 | 3% | 3098 | 2868 | 3391 |
| Myanmar | 13.2 | 6.8 | 49% | 2081 | 2063 | 2107 |
| Philippines | 12.2 | 8.7 | 29% | 2124 | 2074 | 2223 |
| CAR | 8.7 | 6.9 | 21% | 2163 | 2129 | 2207 |
| French Guiana | 8.0 | 7.9 | 1% | 4540 | 4078 | 5107 |
| Lao PDR | 7.9 | 3.0 | 62% | 2068 | 2059 | 2078 |
| Liberia | 7.8 | 5.2 | 33% | 2110 | 2083 | 2148 |
| Viet Nam | 7.0 | 3.0 | 57% | 2072 | 2062 | 2085 |
| India | 6.1 | 3.1 | 50% | 2080 | 2063 | 2102 |
| Nigeria | 6.1 | 2.9 | 53% | 2076 | 2058 | 2103 |
| Thailand | 5.6 | 3.7 | 35% | 2106 | 2071 | 2165 |
| Mexico | 5.3 | 2.1 | 60% | 2069 | 2055 | 2090 |
| Angola | 5.2 | 2.6 | 51% | 2079 | 2062 | 2103 |
| Madagascar | 4.4 | 0.9 | 79% | 2057 | 2047 | 2072 |
| Cote d'Ivoire | 3.6 | 0.0 | 100% | 2033 | 2030 | 2038 |
| Ghana | 3.2 | 0.0 | 100% | 2048 | 2037 | 2067 |
| Nicaragua | 3.2 | 0.1 | 98% | 2050 | 2038 | 2071 |
| Cambodia | 2.6 | 0.0 | 100% | 2041 | 2035 | 2051 |
